# Abstract representations emerge in human hippocampal neurons during inference behavior

**DOI:** 10.1101/2023.11.10.566490

**Authors:** Hristos S. Courellis, Juri Mixha, Araceli R. Cardenas, Daniel Kimmel, Chrystal M. Reed, Taufik A. Valiante, C. Daniel Salzman, Adam N. Mamelak, Stefano Fusi, Ueli Rutishauser

## Abstract

**Humans have the remarkable cognitive capacity to rapidly adapt to changing environments. Central to this capacity is the ability to form high-level, abstract representations that take advantage of regularities in the world to support generalization**^1^**. However, little is known about how these representations are encoded in populations of neurons, how they emerge through learning, and how they relate to behavior**^2,3^**. Here we characterized the representational geometry of populations of neurons (single-units) recorded in the hippocampus, amygdala, medial frontal cortex, and ventral temporal cortex of neurosurgical patients who are performing an inferential reasoning task. We find that only the neural representations formed in the hippocampus simultaneously encode multiple task variables in an abstract, or disentangled, format. This representational geometry is uniquely observed after patients learn to perform inference, and consisted of disentangled directly observable and discovered latent task variables. Interestingly, learning to perform inference by trial and error or through verbal instructions led to the formation of hippocampal representations with similar geometric properties. The observed relation between representational format and inference behavior suggests that abstract/disentangled representational geometries are important for complex cognition.**

## Introduction

Humans have a remarkable capacity to make inferences about hidden states that describe their environment^3–5^ and use this information to adjust their behavior. One core cognitive function that enables us to perform inference is the construction of abstract representations of the environment^5–7^. Abstraction is a process through which relevant shared structure in the environment is compressed and summarized, while superfluous details are discarded or represented so that they do not interfere with the relevant ones^8,9^. This process often leads to the discovery of latent variables that parsimoniously describe the environment. By performing inference on the value of these variables, frequently from partial information, the appropriate actions for a given context can rapidly be deployed^5,10^, thereby generalizing from past experience to novel situations. For example, a latent variable specifying being in a left- or right-driving country can be used by a pedestrian to infer which way to look for oncoming traffic when crossing a road, even in the absence of a sensory cue such as traffic moving in that pedestrian’s field of view, and when crossing roads they have never before encountered. Through abstraction, the common, underlying structure of the world is represented in a way that facilitates adaptive behavior.

How are latent variables encoded in the human brain? How are existing representations modified to accommodate them? Does their format matter for behavior? At the level of neuronal activity, the answers to such questions have remained elusive. Prior research has implicated the hippocampus in the implementation of abstraction and inference-related computations, both through neuroimaging in humans during tasks that require abstraction and generalization^10–13^, and through neurophysiology in rodents and non-human primates engaged in tasks with abstract spatial and non-spatial components^14–18^. To date, relatively few studies have explored the role that the format, or geometry, of task variable representations plays in shaping computation in the human brain at the single neuron level^19,20^, and no study has, to our knowledge, reported the emergence and manipulation of this geometry on the short timescales that would be required for rapid learning in humans.

We recorded the activity of populations of neurons in the brains of awake, behaving epilepsy patients to study the emergence of abstract representations. Patients performed a reversal learning task with two latent contexts, each requiring different responses to the same stimuli. We find that as patients learned to perform inference on the latent context, a representation of context emerged in an abstract format in the hippocampus, being disentangled from simultaneously represented stimulus information (i.e. represented in an approximately orthogonal subspace of the neural activity space). The emergence of the abstract context variable was correlated with an individual’s ability to rapidly perform inference on the state of the latent variable context and was absent during error trials. Furthermore, we found that this abstract hippocampal context representation could emerge in two ways: by learning through experience and through verbal instructions informing patients about the latent structure of the task.

## Results

### Humans perform inference in a context dependent task

Patients viewed a sequence of images and indicated for each whether they thought that the associated action was a “left” or “right” button press on a button box (Fig. 1A). Subjects discovered from the feedback provided after each response what the correct response is for a given image. There were two possible fixed mappings (“SRO maps”, see Fig. 1C) between each of the four stimuli, the associated correct response (Left/Right button press), and monetary reward given for a correct response (25¢ or 5¢). Which of the two fixed mappings should be used depended on which context a given trial was in (Fig. 1B,C, Context 1 or 2). The two contexts alternated every 15-32 trials. Context was a latent variable that had to be inferred by subjects because no information was provided on the screen on which context was presently active or whether it had changed. Critically, the two stimulus-response maps are systematically related: all stimulus-response pairings are inverted between the two contexts (see Fig. 1C). With this design, an individual performing inference can detect a change in latent context after receiving feedback that their response was incorrect in a single trial and immediately update their stimulus-response associations for the remaining stimuli even though they have not yet been encountered in the new context. We refer to the trials in which a given stimulus is encountered for the first time following a covert context switch as inference trials and to the remaining trials as non-inference trials.

**Figure 1.**
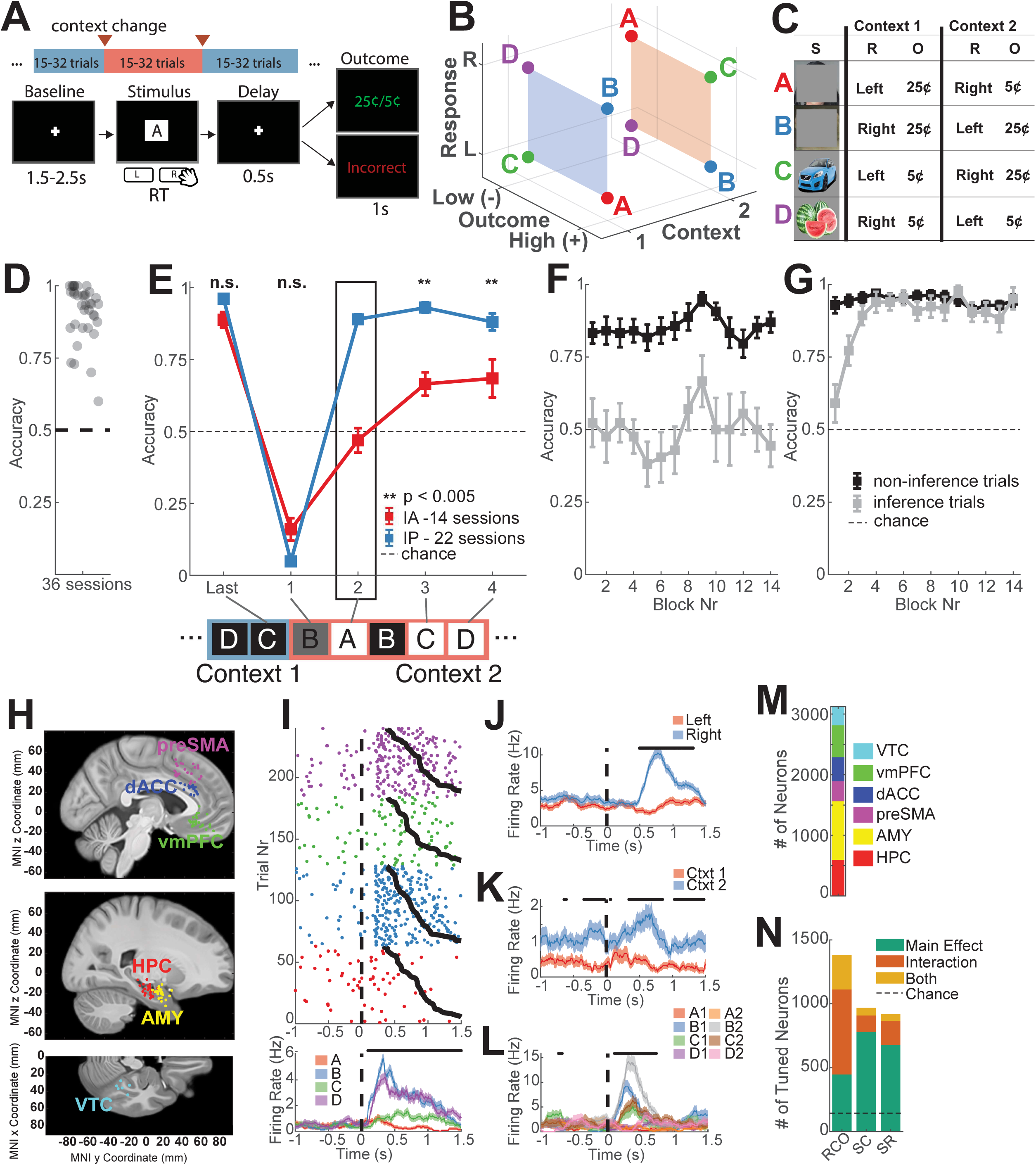
Task, behavior, recording locations, and single-neuron tuning. **(A)** The task consisted of variable-length blocks (15-32 trials) that alternated between two latent contexts (red and blue). Context changes (red arrows) were covert. Trials consisted of a pre-stimulus baseline followed by stimulus presentation during which patients executed the associated response (left or right button press) in a speeded manner. After button press, the stimulus was replaced with a fixation cross, followed by the outcome (either high/low reward or incorrect) was presented after a fixed 0.5s delay. **(B-C)** Illustration of the task structure. Each stimulus (A-D) is associated with a single correct response and results in either a high or low reward if the correct response is given. All stimulus-response-reward relationships are inverted between context 1 (blue) and 2 (orange). This visualization is reflective of the disentangled structure of the task variables, and does not necessarily reflect how neurons will organize their responses in neural state-space to each of these conditions. **(C)** Example images (left) associated with the stimuli A-D. Note: stimuli A and B are masked due to copyright. These associations are randomized for every session. **(D)** Patients exhibited high accuracy on non-inference (baseline) trials. Each dot corresponds to the average non-inference trial performance over a single session. Black dashed line indicates chance. Only sessions where patients exhibited above-chance accuracy on non-inference trials are shown (36/42 sessions, *p* < 0.05, Binomial Test on all non-inference trials). **(E)** Task performance split by whether inference was present (IP) or absent (IA) on the first inference trial following context switches in a given session. Sessions where inference performance was significantly above chance (22 sessions, *p* < 0.05, Binomial Test on inference trial 1) were deemed “inference present” (IP, blue), and those where inference performance was not above chance (14 sessions, *p* > 0.05, Binomial Test on inference trial 1) were considered “inference absent” (IA, red). Plot shows performance on the last trial before the context switch, the first trial after the context switch, and for the remaining three inference trials averaged over all trials in each session (mean ± s.e.m. across sessions). Dashed line marks chance. Black box indicates inference trial 1. **(F-G)** Task performance as a function of time in the task for the **(F)** inference absence and **(G)** inference present groups. Shown is the accuracy for the last non-inference trial before a switch (black) and the first inference trial after a switch (gray). Accuracy is shown block-by-block averaged over a 3-block window (mean ± s.e.m. across sessions). **(H)** Electrode locations. Each dot corresponds to a single microwire-bundle. Locations are shown on the same hemisphere (right) for visualization purposes only. Shown are pre-Supplementary Motor Area (preSMA, purple), dorsal Anterior Cingulate Cortex (dACC, blue), ventromedial Prefrontal Cortex (vmPFC, green), Hippocampus (HPC, red), Amygdala (AMY, yellow), and Ventral Temporal Cortex (VTC, teal). **(I-L)** Example neurons tuned to task variables. **(I**) Hippocampal neuron that encodes stimulus identity. Raster trials are reordered based on stimulus identity, and sorted by reaction time therein (black curves). Stimulus onset occurs at time 0. Black points above PSTH indicate times where 1-way ANOVA over the plotted task variables was significant (p < 0.05). **(J-L**) PSTH of three other example neurons that encode response **(J)**, context **(K)**, and mixtures of stimulus id and context **(L)**. **(M)** Number of single units recorded across all brain areas (3124 neurons recorded in total). **(N)** Number of single units across all brain areas exhibiting significant Main effects or interaction effects (n-way ANOVA with interactions, p < 0.05, see methods) to at least one of the principal task variables (R = Response, C = Context, O = Outcome, S = Stimulus ID) or to combinations of variables. A unit is linearly tuned if it has at least one significant main effect, and non-linearly tuned if it has at least one significant interaction term in the ANOVA model.

Patients (17 total, see Table 1) completed 42 sessions (180-320 trials/session, 10-16 blocks/session) of the task, typically in pairs of two back-to-back sessions on the same recording day (mean = 2.4 sessions per day, min = 2, max = 4, see Table 1). Novel stimuli were used in every session, thus requiring patients to re-learn the SRO maps through trial and error at the start of every session. Of the 42 sessions, 6 were excluded from analysis due to at-chance performance in non-inference trials (Binomial Test, *p* > 0.05). Performance on non-inference trials was well above chance for the remaining 36 sessions (Fig. 1D). Each of the 36 included sessions was classified as either a “Inference Present” (IP) or “Inference Absent” (IA) session depending on whether the patient performed significantly above chance on the first of the three possible inference trials occurring after context switches (Fig. 1E, trials marked 2-4).

Our task is designed such that by performing inference, patients can respond correctly the first time they see an image in a new context following the initial error trial (seen as significantly below chance performance in Fig. 1E at time point 1). This can be achieved by patients flexibly updating the currently active SRO map immediately after encountering an error, thereby allowing them to perform accurately for the remaining three stimuli that had not yet been seen in the new context. We took accuracy on the first of these three opportunities (the first inference trial) following a context switch as the behavioral signature of successful behavioral inference (timepoint 2 in Fig. 1E, Binomial Test, *p* < 0.05). Block-wise estimates of task performance for IA (Fig. 1F) and IP (Fig. 1G) sessions reveal that during IA sessions, patients exhibit poor inference performance after every context switch throughout the task, although the performance at the end of every block is high. In contrast, during IP sessions, inference performance rapidly rose over the first few blocks and remained high throughout the duration of the session. Note that within a given session, the two latent contexts had identical stimuli, responses, and outcomes; the only difference was which stimulus was associated with which response and outcome. Correspondingly, subject-level accuracies (Fig. S1B) and reaction times (Fig. S1C) for the two contexts (arbitrarily labeled 1 and 2 across sessions) were not significantly different, indicating that there was no systematic performance bias for one of the two contexts.

When first performing the task, patients were told that they needed to learn arbitrary stimulus-response associations that would change over time, but were not informed about the latent contexts and their related structure. Sessions were recorded in back-to-back pairs with verbal instructions (see Methods) detailing the latent structure of the task provided during the inter-session period (mean length of break = 241 s, range 102-524s), which was considerably shorter than the sessions themselves (mean = 1154 s, range 898-1900s). Importantly, the session following the verbal instructions required re-learning the SRO maps for new stimuli. We next considered whether patients could discover the latent task structure before receiving instructions, and if not, whether verbal instructions successfully shaped behavior. Patients were split into three groups: An “instruction successful” (IS) group, which is composed of patients who did not perform inference during pre-instruction sessions and who did perform inference during post-instruction sessions (5 patients, 10 sessions, Fig. S1D); An “instruction unsuccessful” (IU) group, which were patients who did not perform inference during both the pre-and post-instruction sessions (4 patients, 8 sessions, Fig. S1E); and a “session one interference” (S1) group, which were patients who exhibited inference behavior during both pre- and post-instruction sessions (3 patients, 6 sessions, Fig. S1F). Only patients who performed accurately in non-inference trials in both pre-and post instruction sessions were included in one of these three groups (Fig. S1D-F, “last”; 5 patients excluded, see Table 1). For each subject, no sessions after the first two were considered for this analysis. Thus, patients exhibited a variety of inference behaviors. Below, we contrast the neural representations between these groups of patients to examine how instructions shape neural representations on short timescales.

### Electrophysiology and analysis approach

Neural data recorded over the 36 (of 42) included sessions yielded 2694 (of 3124) well isolated single-units, henceforth neurons, distributed across the hippocampus (HPC, 494 neurons), amygdala (AMY, 889 neurons), pre-supplementary motor area (preSMA, 269 neurons), dorsal anterior cingulate cortex (dACC, 310 neurons), ventromedial prefrontal cortex (vmPFC, 463 neurons), and ventral temporal cortex (VTC, 269 neurons) (Fig. 1H,M). Only well isolated neurons as assessed by rigorous spike sorting quality metrics were included (see methods). Action potentials discharged by these neurons were counted during two 1s long trial epochs: during the baseline period (base, -1s to 0s prior to stimulus onset), and during the stimulus period (stim, 0.2s to 1.2s after stimulus onset). For the stimulus period, since patients would sometimes respond before 1.2s (reaction time = 1.08 ± 0.04s over sessions), we determined that 75.15% of all spikes occurred before a response was provided across all recorded neurons, indicating that analyses performed with these spike counts predominantly, but not exclusively, reflect pre-decision processing.

Single neuron responses during the two analysis periods were heterogeneous. During the stimulus period, some neurons exhibited selectivity to one or several of the four variables stimulus identity, response, (predicted) outcome, and context (Fig. 1I-K, example PSTHs of stimulus identity, response, and context coding neurons). Other neurons were modulated by combinations of these variables (Fig. 1L, PSTH of a neuron with conjunctive stimulus and context coding). Across all brain areas, 54% of units (1447/2694) were tuned to task variables, with 26% of units (706/2694) exhibiting only interaction effects, 17% (449/2694) exhibiting only main effects, and 11% (292/2694) exhibiting both when fitting a 3-Way ANOVA for Response, Context, and Outcome (Fig. 1N, RCO column, chance = 135/2694 units, factor significance at *p* < 0.05). When neurons were separated into those recorded from IA and IP sessions, 5-15% of neurons in IA and IP sessions were significantly tuned for each of the main and interaction effects of the 3-Way ANOVA, with significant reductions in the proportion of neurons tuned to Outcome, Response x Context, and Response x Context x Outcome (*p*_O_ = 0.0007, *p*_RxC_ = 0.0395, *p*_RxCxO_ = 0.0048, two sample z-test) in IP sessions compared to IA sessions (Fig. S1M). Similar analyses conducted on a separate 2-Way ANOVA for Stimulus Identity and Context (Fig. 1N, SC column, Fig. S1N, *p*_S_ = 0.0165, two sample z-test comparing IA with IP sessions), and for Stimulus Identity and Response (Fig. 1N, SR column, Fig. S1O, *p*_S_ = 0.0287), revealed a significant decrease in the fraction of neurons tuned to Stimulus Identity in IP compared to IA sessions, again with no significant changes in the proportion of neurons coding for Context, Response, or interactions. These findings indicate diverse tuning to many task variables simultaneously across all brain regions.

### Measures of Neural Population Geometry

Given the heterogenous nature of the response pattern at the single neuron level (also see Fig. S1G-L), we adopted a population-level approach. This approach allows us to assess which variables are encoded in the distributed neural patterns, considering also the correlations of the neural responses across multiple conditions. Most importantly, it enables us to examine how these variables are represented, and, in particular, to study the geometry of the neural representation, which has important computational implications. In our task, the geometry of a representation is defined by the arrangement of the eight points in the activity space that represent the experimental conditions. Low dimensional geometries (e.g., when the eight points define a cube) are typically the result of a process of abstraction and confer to a linear readout the ability to generalize. For example, consider a simplified situation with three neurons (the axes) and two stimuli in two contexts, each associated with one of two possible responses (Fig. 2A-C). Imagine that the 4 points (2 per context) are arranged on a relatively low dimensional square (the maximal dimensionality for 4 points is 3), with context encoded along one side and the response along one of the two orthogonal sides (Fig. 2A). Then, a linear decoder for context, trained only on the conditions for which the response is Right, can readily generalize to the Left response conditions (Fig. 2B). This ability to generalize is due to the particular arrangement of the points, which make the coding direction of context for the left responses parallel to the context coding direction for the right responses (Fig. 2C). Also, in this case, context and response are represented in orthogonal subspaces, and hence, they are called disentangled variables^21,22^. In the square example, the ability of a linear decoder to generalize across conditions (cross condition generalization) also applies to the variable response (i.e. a response decoder trained on context 1 conditions will generalize to context 2 conditions). In previous work^14^, we used the cross-condition generalization performance (CCGP) as the defining characteristic of an abstract representation of a variable. Indeed, when context CCGP is high, as in the square example, there is a subspace (the 1D space defined by the context coding direction) in which the representation of context does not depend on the response, or in other words, it is disassociated from the specific instances of the response (Left or Right). In this case we would say that context is represented in an abstract format, or more synthetically that the representation of context is abstract.

**Figure 2.**
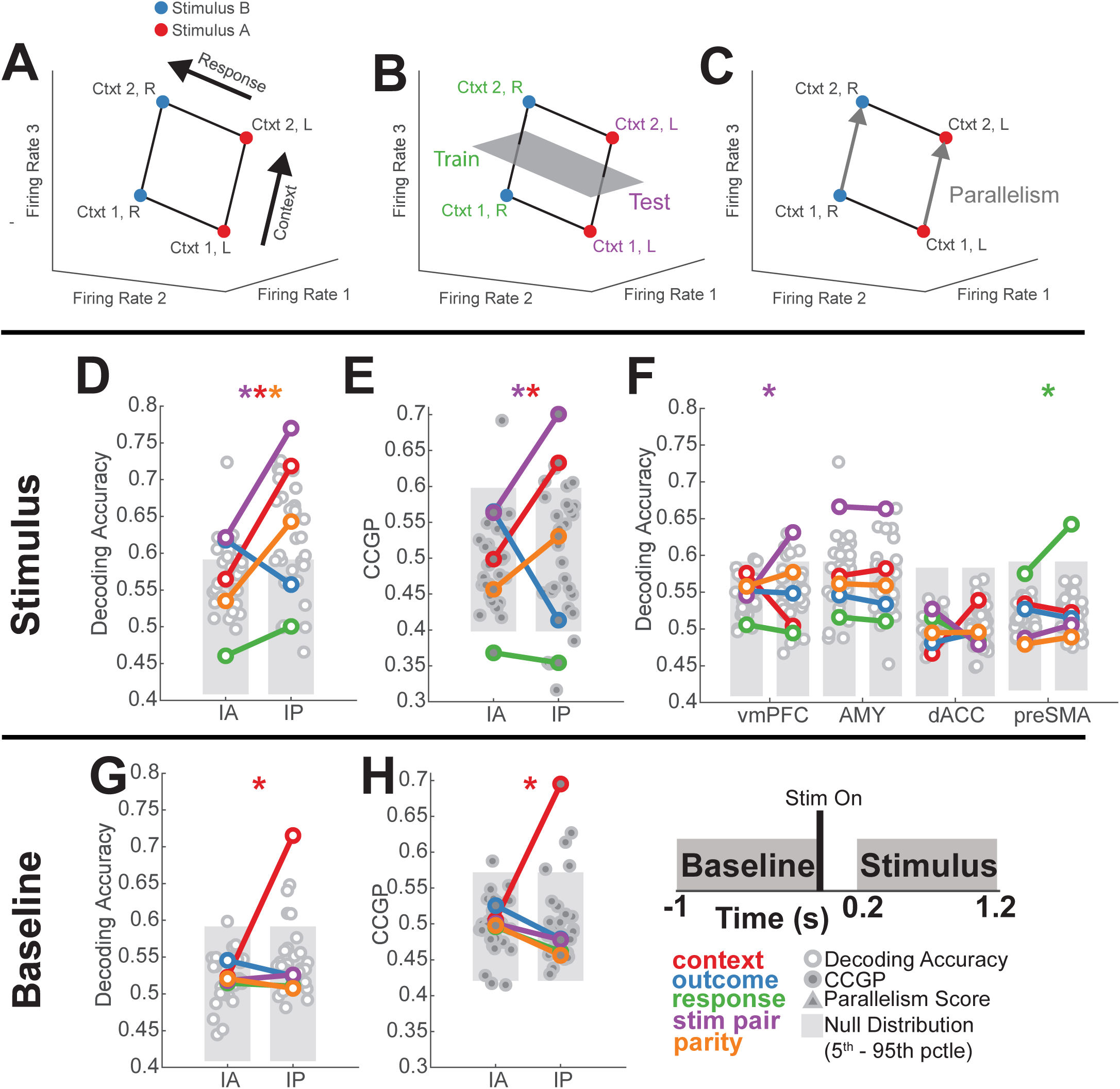
Emergence of multiple abstract variables in hippocampus supports inference. **(A)** Simplified example of a neural state space where each axis is the firing rate of one neuron. Points correspond to the response of the neurons to different task states, i.e. two stimuli (red and blue) that elicit two responses (R and L) in two contexts. Note the coding vectors for response and context (black arrows) are not aligned with the axes as each neuron might respond to mixtures of variables. The axes of this state space differ from those shown in Fig. 1B, with the latter being defined by experimenter-selected variables rather than neural firing rates. **(B)** Example of cross-condition generalization. A decoder is trained to classify context only on response “R” conditions (green) and is evaluated on its ability to decode context on response “L” conditions (purple). If context is represented in an abstract format (i.e. disentangled from response), then the decoder should generalize to the held-out response condition, yielding a high cross-condition generalization performance (CCGP) for context. **(C)** Example of context parallelism. Coding vectors for context (gray arrows) are parallel, indicating that the coding direction for context is identical for different responses, and thus that context and response are disentangled. Details for computing the parallelism score (PS) of a balanced dichotomy with 8 conditions are provided in the methods. **(D-F)** During the stimulus presentation period, context (red) and stimulus pair (purple) become decodable in the IP group in the HPC. Context is encoded in an abstract format. Decoding accuracy **(D)** and CCGP **(E)** are shown for all 35 balanced dichotomies during the stimulus period (0.2 to 1.2 s following stimulus onset, see inset). A subset of the dichotomies are named (color code) because they represent task specific variables (see Fig. S2). Swarm plots for decoding accuracy and CCGP are light circles and dark circles respectively. Gray bars denote the 5^th^-95^th^ percentile of the shuffle-null distribution for decoding accuracy and geometric null distribution for CCGP. Stars denote named dichotomies that are above chance in IP and are significantly different from their corresponding IA value (*p*_RS_ < 0.05/35, Ranksum Test, Bonferroni corrected for multiple comparisons across all dichotomies). **(F)** Identical analysis to **(D)** showing decodability of balanced dichotomies from neurons recorded in other brain regions (except VTC, which is shown in Fig. 4 and S8). **(G-H)** Same as for **(D-F)**, but for spikes counted during the baseline period prior to stimulus onset. Context (red) becomes decodable in IP sessions and is in an abstract format. Trials are labeled according to the current trial. Decoding accuracy **(G)** and CCGP **(H)** computed in HPC for all balanced dichotomies with spikes counted during the pre-stimulus baseline period (-1 to 0s prior to stimulus onset, see inset). All plotting conventions identical to those in **(D-F)**, except Baseline analysis is conducted with task variables from previous trial. Note: all reported geometric measures, decoding accuracies, or angles are the average of 1000 runs with condition-wise trial resampling as described in the methods. All null distributions are constructed from 1000 iterations of shuffled trial-resampling using either trial-label shuffling (shuffle null) or random rotations designed to destroy low-dimensional structure (geometric null). Also, neuron counts are balanced between IA and IP for every brain area to ensure that dimensionally-sensitive values (e.g. vector angles, decoding accuracies, etc..) are directly comparable. See methods for details.

Notice that if the 4 points of the example are at random locations in the activity space defining a tetrahedron, the representation is “unstructured” and does not have any of the generalization properties we described. On the other hand, these high dimensional representations allow a linear readout to separate (or shatter) the points in any arbitrary way, and hence confer to the readout the flexibility of implementing any possible task. We refer to the number of ways the points can be separated into two groups by a linear decoder (dichotomies) as shattering dimensionality (SD)^14,23^. Recorded neural representations can have both the generalization properties of the abstract representations, and the flexibility of the high dimensional representations^14^.

We characterized and compared the representational geometry between IA and IP sessions for neural pseudopopulations of all recorded neurons in each brain area. For each variable, or dichotomy, we computed the decoding accuracy, which tells us whether the variable is encoded, and the CCGP, which indicates whether the representation of that variable is disentangled from other variables. Both decoding accuracy and CCGP are reported in a cross-validated manner by training and testing decoders on single trials. We complemented this single-trial analysis with a third metric that we call the the parallelism score (PS). PS measures the parallelism of the coding directions of a specific variable. The coding directions are estimated using the average activity for each condition. A high PS strongly indicates that the variable is represented in an abstract format. The PS is also a more direct geometrical measure that focuses on the structure of the representation (the CCGP depends on the geometry but also on the noise and its shape).

To perform the analysis in an unbiased manner, we did not consider only the variables defining the task, but all the variables that correspond to the 35 possible dichotomies of the 8 conditions (see Fig. S2 for illustration of five dichotomies that correspond to specific task variables, Table 2; we refer to these as named dichotomies).

### Hippocampal neural geometry correlates with inference behavior

We first examined the decodability of each balanced dichotomy in different brain areas for sessions where inference was present (IP) and sessions where inference was absent (IA). Following stimulus onset, the hippocampal neural population exhibited a significant increase in average decodability across all balanced dichotomies in IP sessions relative to IA sessions, indicative of an increase in shattering dimensionality (Fig. 2D, IA vs IP *p*_RS_ = 2.7×10^-3^, Ranksum over dichotomies). Latent context (Fig. 2D, red, IA vs. IP *p*_RS_ = 2.9×10^-27^, *p*_lA_ = 0.12, *p*_lP_ = 5.1×10^-5^; p_IA_ and p_IP_ are significance tests vs. chance and p_RS_ a pairwise comparison between IA and IP) and stim pair (Fig. 2D, purple, IA vs. IP, *p*_RS_ = 5.0×10^-27^, *p*_lA_ = 0.015, *p*_lP_ = 7.9×10^-7^) emerged as the most strongly decodable named dichotomies in the IP sessions. The stim pair dichotomy corresponds to the grouping of stimulus identities that elicit the same response (D&B vs. A&C), a relationship that remains the same across both contexts (Fig. 1C, S2). This difference in representation between IA and IP sessions is unique to HPC, as no other recorded region exhibited a significant change in decodability of the variable context (Fig. 2F, red). Rather, in vmPFC stim pair (Fig. 2F, purple, IA vs. IP, *p*_RS_ = 6.9×10^-20^, *p*_lA_ = 0.21, *p*_lP_ = 0.0097) and in preSMA response (Fig. 2F, green, IA vs. IP, *p*_RS_ = 5.2×10^-1^, *p*_lA_ = 0.091, *p*_lP_ = 0.0057) increased significantly in decodability in IP compared to IA sessions, which is as expected (see discussion). The stim pair variable was also significantly decodable in AMY both during IA and IP, with no significant difference between the two (Fig. 2F, purple, IA vs. IP, *p*_RS_ = 0.88, *p*_lA_ = 0.0016, *p*_lP_ = 0.0018).

The expressiveness of the HPC representation can be quantified by the decodability of dichotomies that probe for non-linear interactions of variables in the population. The parity dichotomy (Fig. S2) is only decodable if variables are encoded with a high degree of non-linear interactions in a neural population (see methods). We observed that in the HPC but not in other brain areas, parity decodability increased significantly in IP relative to the IA sessions (Fig. 2D, orange, IA vs. IP, *p*_RS_ = 1.5×10^-21^, *p*_lA_ = 0.27, *p*_lP_ = 0.0055). Generalizing this finding, dividing different dichotomies into increasing levels of “difficulty”, with more difficult dichotomies requiring stronger non-linear interactions of task variables, reveals that average shattering dimensionality is highest for the most difficult dichotomies in the HPC (Fig. S7). Together, these findings suggest that non-linearities in the HPC population response in the IP relative to the IA group led to an increase in the number of dichotomies that could be decoded by a linear decoder. Notably, only the hippocampus exhibited significant parity decodability (Fig. 2F, orange, all *p* > 0.05), a significant increase in shattering dimensionality (Fig. 2F), and the emergence of multiple, simultaneously decodable dichotomies between IA and IP sessions.

We next examined the format of the decodable named dichotomies (context, stim pair, parity). During the stim period, CCGP (Fig. 2E, S3D) was significantly elevated for both the context (Fig. 2B, red, IA vs. IP, *p*_RS_ = 2.0×10^-28^, *p*_lA_ = 0.51, *p*_lP_ = 0.02) and stim pair (Fig. 2E, purple, IA vs. IP, *p*_RS_ = 2.0×10^-28^, *p*_lA_ = 0.17, *p*_lP_ = 0.0011) variables in IP but not in IA sessions. In addition, the PS was significantly larger than expected by chance for the variables context and stim pair in the IP but not IA sessions (Fig. S3E, red, *p*_lA_ = 0.55, *p*_lP_ = 1.4×10^-15^ *and* Fig. S3E, purple, *p*_lA_ = 0.17, *p*_lP_ = 1.7×10^-8^). CCGP larger than expected by chance was also observed for the stim pair variable in vmPFC during IP and not in IA (Fig. S3A, purple, *p*_lA_ = 0.45, *p*_lP_ = 0.014), the response variable in preSMA (Fig. S3A, green, *p*_lA_ = 0.050, *p*_lP_ = 0.0010), and the stim pair variable in AMY during both IA and IP (Fig. S3A, purple, *p*_lA_ = 0.050, *p*_lP_ = 0.039). Taken together, the increased CCGP and PS values in IP relative to IA sessions indicate that the context and stim pair variables are both simultaneously represented in an abstract format in the HPC in sessions where inference behavior was observed (IP sessions). This means that for example a decoder for context trained only on the conditions in which the subject responded Left, can readily generalize to all the conditions in which the subject responded Right. These two variables were not represented in an abstract format when inference behavior is absent (IA sessions). While other task variables were also represented in an abstract format in other brain regions, only the hippocampus simultaneously represented these two variables in an abstract format (Fig. S3A,B). These two disentangled variables are thus represented in approximately orthogonal subspaces.

We also conducted a parallel analysis during the pre-stimulus baseline (Fig. 2, inset), analyzing the geometry of persistent representations of the previous trial. We found that context alone was encoded in an abstract format in the HPC only in sessions in which subjects could perform inference (Supplement S.1). This finding indicates that the hippocampal context representation was persistently maintained in an abstract format across trial epochs.

Together, these findings indicate the emergence of an abstract representation of latent context in the HPC that is correlated with patient’s ability to perform inference. This context variable is encoded in the hippocampal representation alongside: (i) an abstract, behaviorally-relevant stimulus category variable (stim pair, but see below), and (ii) an increase in parity dichotomy decodability and increased shattering dimensionality from IA to IP sessions. Together, these findings indicate that the ability to perform inference correlates with the hippocampus explicitly encoding a latent context variable (whose state is not directly observable). The resulting representation simultaneously exhibited both features of high dimensionality (parity decodability, increased SD) and low dimensionality (two variables in an abstract format).

### Hippocampal representation of context is absent in incorrect trials

We next asked whether the presence of context as an abstract variable was associated with trial-level performance. To examine this question, we compared the decodability and geometry of all dichotomies between correct and incorrect (error) trials in inference present sessions. The first trial of every block was excluded from this analysis due to being necessarily incorrect by design (see Fig. 1E, time point 1). Contrasting correct with incorrect trials, we found that scattering dimensionality was significantly higher in correct trials (Fig. 3A, IP-c vs. IP-i, average SD 0.59 vs 0.54, *p*_RS_ = 0.0048), as was decodability of the parity dichotomy (Fig. 3A, IP-c vs. IP-i, orange, *p*_RS_ = 0.029). Furthermore, the dichotomies context and stimulus pairing were significantly more decodable in correct compared to incorrect trials (Fig. 3A, IP-c vs. IP-i, red, *p*_RS_ = 1.2×10^-20^, purple, *p*_RS_ = 1.0×10^-9^). Furthermore, PS for context but not other variables was significantly elevated in correct trials but not in incorrect trials (Fig. 3B, red, IP-c vs. IP-i, *p*_lP-c_ = 1.1×10^-1^, *p*_lP-i_ = 0.083) in IP sessions. Similarly, the baseline representation of context also showed this effect, being decodable during correct trials but not during incorrect trials (Fig. 3C, red, IP-c vs. IP-i, *p*_RS_ = 3.5×10^-12^, *p*_lP-c_ = 0.012, *p*_lP-i_ = 0.47). Context was present in an abstract format only in correct trials based on the PS for context being significantly larger than chance during correct but not incorrect trials (Fig. 3D, red, IP-c vs. IP-i *p*_lP-c_ = 0, *p*_lP-i_ = 0.94). The lack of decodability and parallelism of context in the baseline immediately prior to an incorrect trial indicates that the geometry of the representation at baseline is correlated with correct behavior in the upcoming trial. Together, these findings demonstrate that both the content and format of the hippocampal neural representation is correlated with behavior on a trial-by-trial basis. This effect is present during the stim period in incorrect trials, where shattering dimensionality, context decodability, and context PS significantly decreased. This effect is also present during the baseline period, where reduction in the decodability and parallelism of the context variable is associated with an error on an upcoming trial.

**Figure 3.**
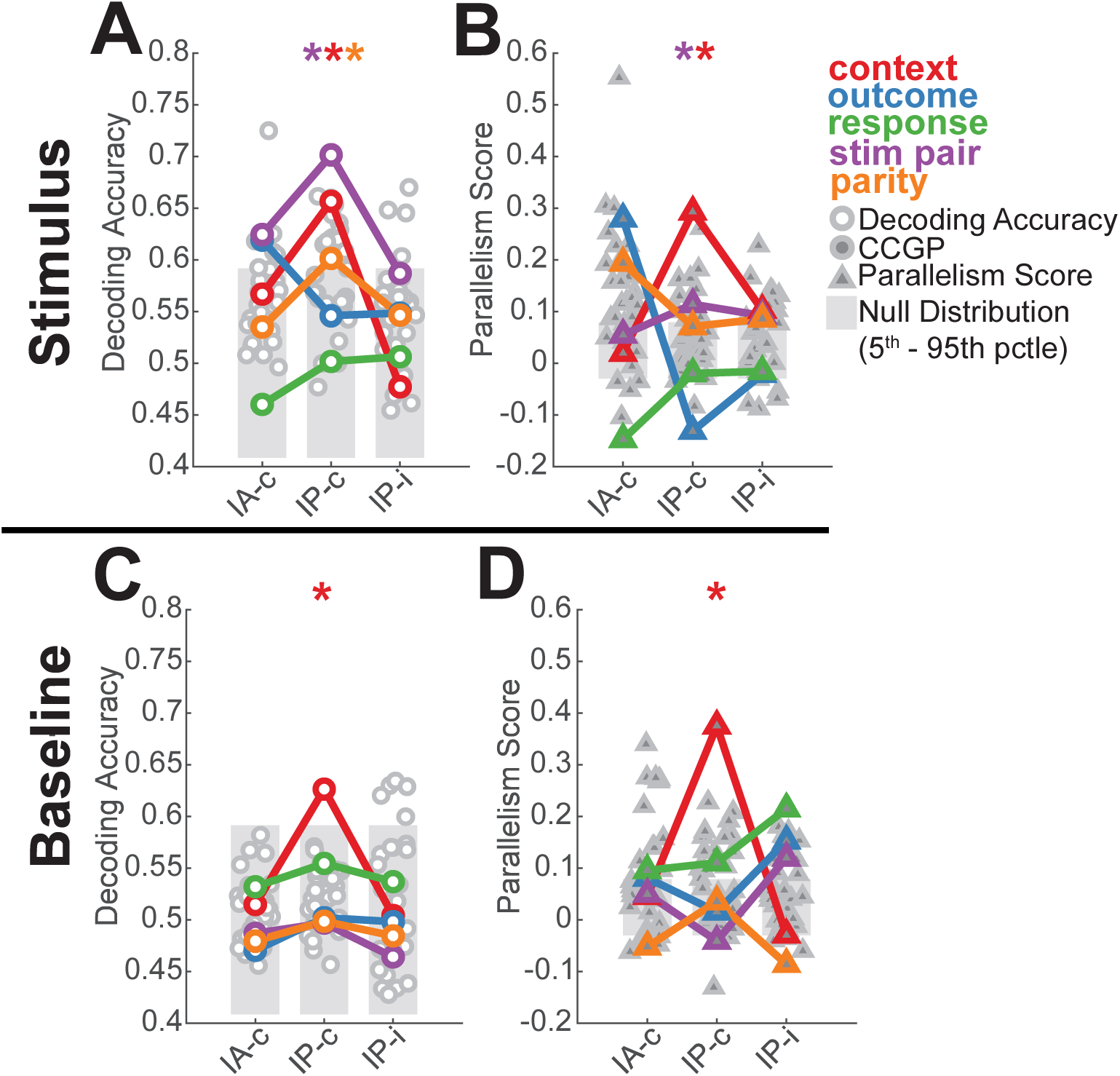
Geometry of hippocampal context representations is compromised during error trials. **(A-B)** Following stimulus onset, context (red) is not decodable (A) and not in an abstract format (B) in incorrect trials occurring during IP sessions. Decoding accuracy and parallelism scores are estimated separately during correct and incorrect trials in IP sessions (IP-c and IP-i, respectively) and in correct trials only during IA sessions (IA-c). Stars denote named dichotomies that are above chance in the IP-c trials and are significantly different from their corresponding IA-c value (*p* < 0.05/35, Ranksum Test, Bonferroni multiple comparison corrected across all dichotomies). **(C-D)** Same as **(A-B)**, but estimated for spikes counted during the baseline period and task states from previous trial. Note: IP-c values here are slightly lower than the IP values reported in Fig. 2 because additional neurons were removed by subsampling to equalize the number of neurons used for both correct and incorrect trials.

### Controls for univariately tuned neurons, seizure-onset zones, and non-inference performance

We performed three sets of control analysis. First, to determine the relationship of our population-level findings to classical (univariate) tuning, we repeated our analyses after removing subsets of neurons from the population. We removed neurons with significant main effects in a (i) a 2×2×2 ANOVA for Response, Context, and Outcome, and (ii) a 4×2 ANOVA for Stimulus Identity and Context (Supplement S.2). In both analyses, context remained significantly decodable and in an abstract format. Also, as expected, in (ii), stimulus pair representations were no longer decodable. These control analyses indicate that the representation of context in the hippocampus we identified did not arise only due to the emergence of classically context tuned neurons. Rather, context was represented by broadly distributed context modulation at the level of the population.

Second, to assess whether our results were influenced by pathology, we repeated our analysis after excluding hippocampal neurons that were located within clinically confirmed seizure onset zones (Supplement S.3). We found no quantitative changes in our results, suggesting that hippocampal pathology did not influence our results.

Lastly, we examined whether our results were sensitive to behavioral accuracy in non-inference trials (Fig. 1E, trial indicated as ‘last’). We repeated our analysis in a subset of IA and IP sessions that were chosen such that non-inference trial performance was matched (Supplement S.4). This control analysis revealed no qualitative changes in our results, suggesting that differences in non-inference trial performance cannot explain our results.

### Abstraction of stimulus coding across contexts uniquely increases in the hippocampus

Individual HPC neurons in humans prominently encode the identity of visual stimuli^24^. Visually tuned neurons, whose firing rate is strongly modulated by the identity of presented images, are a prominent example of such encoding^25^. We therefore next asked how the variable context, which we show above is encoded in the HPC, interacts with the stimulus identity variable. To examine this question, we conducted geometric analyses over pairs of stimuli (e.g. stimulus A vs B) in the two contexts. This approach is different from the analysis shown so far, in which we considered all 35 possible balanced dichotomies of task conditions. In each balanced dichotomy, different stimulus identities are grouped together (e.g. stimuli A&B vs stimuli C&D), thereby not allowing an examination of the representational geometry for individual stimuli. To examine how stimulus identity encoding relates to context encoding, we instead consider all possible stimulus pairs individually (A vs. B, A vs. C, A vs. D, B vs. C, B vs. D, and C vs. D) in both contexts. As a comparison, we concurrently performed this analysis on neurons recorded in ventral temporal cortex (VTC). VTC neurons were strongly modulated by stimulus identity (see below)^26,27^, but context was not decodable at the level of the balanced dichotomy analysis (basline period: Fig. S8A, red; compare with Fig. 2G; stimulus period: Fig. S8B, red, compare with Fig. 2D).

First, we verified that neurons in both areas encoded the identity of the four stimuli presented. This was the case in both HPC and VTC: 109/494 (22%) of neurons in HPC (Fig. 4A shows an example) and 195/269 (73%) of neurons in VTC (Fig. 4F shows an example) were significantly modulated by stimulus identity following stimulus onset (1×4 ANOVA, p < 0.05). Similarly to HPC, at the population level, VTC neurons encoded stimulus identity-related balanced dichotomies in an abstract format (Fig. S8B-D, purple, brown, pink, *p*_lA/lP_ < 10^-10^). Furthermore, error trial analysis revealed that stimulus-related dichotomies were also decodable during errors in VTC (Fig. S8E, purple, brown, pink, *p*_lP-i_ < 10^-10^). This finding contrasts with HPC (compare to Fig. 3A, stim pair AC vs. BD dichotomoy) and is consistent with the idea that neurons in VTC veridically represented the stimulus viewed on the screen by the patient during both correct and error trials. These analyses demonstrate that VTC and HPC neurons encode stimulus information, allowing us to next examine how context interacted with stimulus identity coding in these two brain areas.

**Figure 4.**
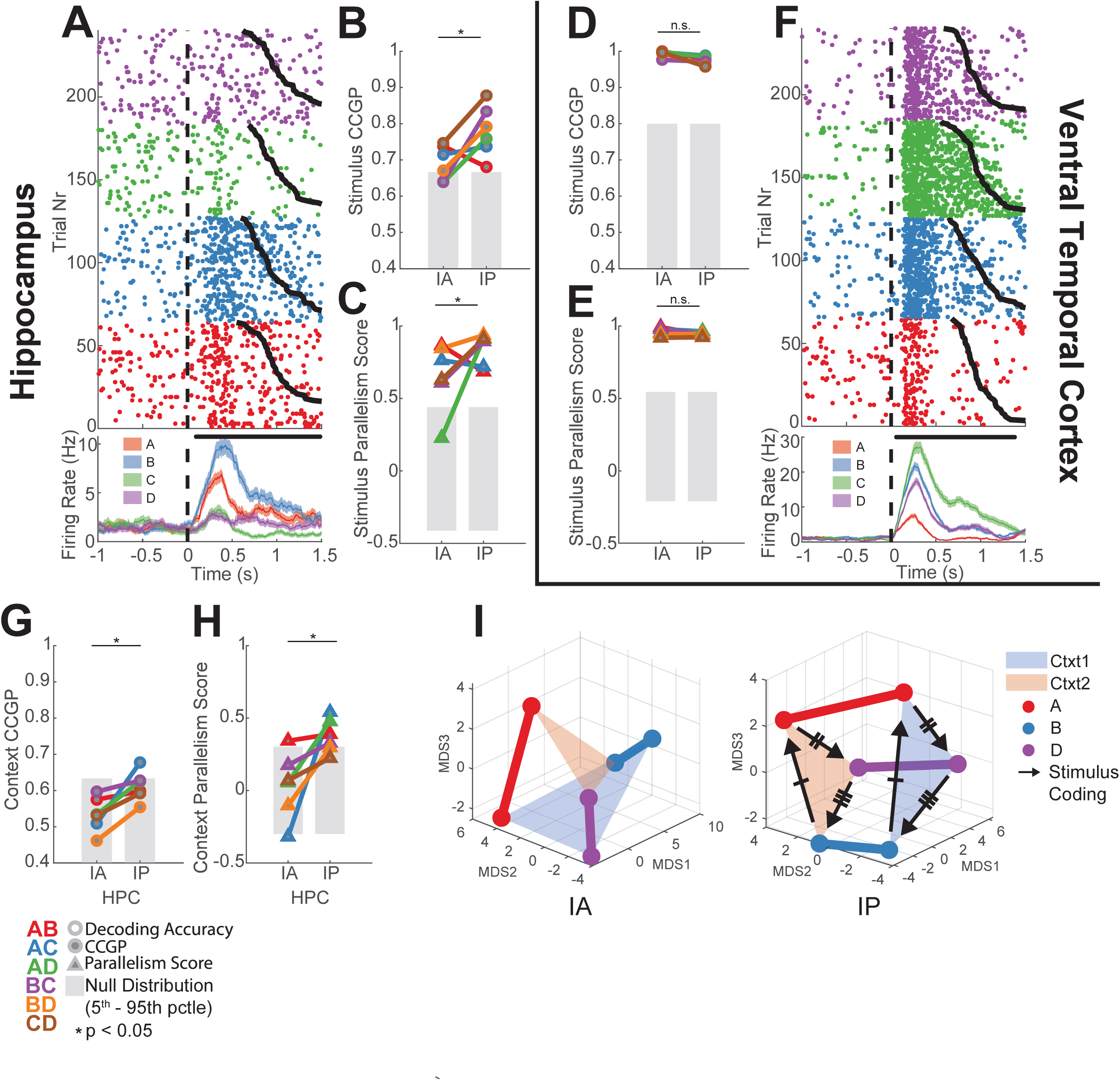
Stimulus representations become structured around context with inference in HPC but not VTC. **(A-C)** Responses in HPC following stimulus onset carry information about stimulus identity. **(A)** Example neuron in the HPC. Trials are reordered by stimulus identity, and are sorted by reaction time therein (black line). **(B,C)** Stimulus geometry across contexts, with geometric analysis conducted over pairs of stimuli in each context. Data points shown correspond to different stimulus pairs (color coded, see lower right for legend). Significance of differences is tested using RankSum comparing IA and IP over all stimulus pairs (* indicates *p* < 0.05, n.s. otherwise). All other conventions identical to those in Fig. 2,3. **(B)** CCGP (*p*_*RS*_ = 0.041) and **(C)** parallelism score (*p*_*RS*_ = 0.040) for stimulus coding across contexts significantly increased in IP compared to IA. **(D-F)** Same as **(A-C)**, but for VTC. **(D)** CCGP (*p*_*RS*_ = 0.15) and **(E)** parallelism score (*p*_*RS*_ = 0.39) for stimulus coding across contexts does not differ significantly between IA and IP. **(G-H)** Context encoding across stimulus pairs for HPC. Plotting conventions are identical to those in panels (B-E). **(G)** CCGP for context across stimuli (*p*_RS_ = 0.012) and **(H)** parallelism score for context coding vectors between pairs of stimuli (*p*_RS_ = 0.015) both significantly increase from IA to IP. **(I)** Changes in neural geometry in HPC. MDS of condition-averaged responses of all recorded HPC neurons shown for IA (left) and IP (right). Colored points are average population vector responses to a stimulus (point color) in each context (plane color). Stimuli of the same identity in either context are connected by a line of the same color. Black arrows represent the coding direction between two different stimuli in each context. Comparing left with right, note that context emerges as a decodable variable (red and blue planes separate), and both stimulus coding (black arrows) and context coding (colored lines) are parallel reflecting the abstract format of both variables. Note: only three stimuli are shown here since the full stimulus ID space is 3D (4 unique, unrelated stimuli) and context is encoded in its own independent dimension, thus requiring 4 dimensions to fully visualize the neural geometry of the hippocampus during inference. All 2D projections of stimulus pairs are shown in Fig. S12.

We next conducted a geometric stimulus-pair analysis to study the interaction of stimulus identity and context coding in the same neural population. The stimulus-pair analysis was designed to detect the presence of simultaneous abstract coding of stimulus identity across contexts (see Fig. S9A-C for illustration) and abstract coding of context across stimuli (see Fig. S9D-F for illustration).

The average stimulus decoding accuracy across all stimulus pairs in the HPC did not differ significantly between IA and IP sessions (0.73 vs. 0.76; Fig. S10C, *p*_RS_ = 0.13, RankSum over stimulus pairs), indicating that the decodability of stimulus information was not different when patients could perform inference vs. when they could not. In contrast, the geometry of the stimulus representation became disentangled from context: both the stimulus CCGP (Fig. 4B, *p*_RS_ = 0.041) and stimulus PS (Fig. 4C, *p*_RS_ = 0.040) were significantly increased in IP compared to IA sessions. This means that a decoder trained to differentiate between stimulus A and B in one context generalized better to the other context in IP compared to IA sessions (and vice-versa for all other possible pairs). This finding suggests that the stimulus responses reorganized with respect to the emerging context variable. Note that context was not decodable in IA sessions as a balanced dichotomy (Fig. 2D, red line). Nevertheless, stimulus decoders did not generalize well between the two contexts in IA sessions. This result indicates that context did modulate stimulus representations in the HPC, but in a way that was entangled with stimulus identity (see below). This effect was specific to the HPC: in VTC, the neural population geometry was unchanged, as indicated by no significant differences in stimulus decodability (Fig. S10D, *p*_RS_ = 0.15), stimulus CCGP (Fig. 4D, *p*_RS_ = 0.15) and stimulus PS (Fig. 4E, *p*_RS_ = 0.39) between IA and IP sessions. In VTC, CCGP was high even in the IA session, indicating that context was not entangled with stimulus identity like in HPC. These analyses demonstrate that the representational geometry for stimulus identity in HPC becomes significantly more disentangled from context in IP sessions compared to IA sessions in a manner that reflects an abstract format.

Having explored the generalization of stimulus encoding across contexts, we next turned our attention to study the generalization of the context code across stimuli. The presence of abstract coding for one variable (stimulus identity) does not necessarily imply that the other variable is also present in an abstract format, though we do have evidence that this is the case in hippocampus from the CCGP and PS analysis over balanced dichotomies (Fig. 2B, S4E). Context decoding analysis conducted over stimuli (e.g. considering only trials with stimuli A&B shown, decode context) in HPC revealed that context was decodable for many stimuli both during IA and IP sessions, without a significant difference between the two (0.63 vs. 0.67; Fig. S11A, *p*_RS_ = 0.065). However, despite being decodable, context encoding during IA sessions was not in an abstract format for stimulus pairs as indicated by low context CCGP (Fig. 4G, *p*_lA_ > 0.10 for all stim pairs) and low context PS (Fig. 4H, *p*_lA_ > 0.17 for all stim pairs except for AB, where *p*_lA_ = 0.033) values that were not significantly greater than chance. In contrast, in IP sessions, both context CCGP (Fig. 4G, *p*_RS_ = 0.012) and context PS (Fig. 4H, *p*_RS_ = 0.015) increased significantly relative to the IA group. This finding indicates that context emerged as an abstract variable at the level of individual stimulus pairs in the HPC.

We next contrast these findings with VTC. While context was decodable from some stimulus pairs during both IA and IP sessions (Fig. S11B, *p*_lA_ ∈ (0.013, 0.074), *p*_lP_ ∈ (0.020, 0.081) for all stim pairs), there was no significant change in context decodability between IA to IP sessions (Fig. S11B, *p*_RS_ = 0.18). Rather, there was a significant decrease in context CCGP (Fig. S11C, *p*_RS_ = 0.026) and no significant difference in context PS (Fig. S11D, *p*_RS_ = 0.39) from IA to IP. Together, these findings indicate that in the HPC, the context variable in the IP sessions is in an abstract format because context coding directions become aligned (i.e. parallel) across stimuli. For VTC, on the other hand, the lack of an abstract context representation across stimulus identities in both IA and IP sessions suggests that context, though decodable for individual stimulus pairs, is not organized in an abstract format and does not meaningfully correlate with inference behavior.

In summary, these findings indicate that the emergence of context as an abstract variable in the hippocampus when patients can perform inference is coupled with the reorganization of stimulus representations so they are also more disentangled, thereby forming a jointly abstracted code for stimuli and context. This transformation of the representation is visible directly in the data when projecting the neural representations in 3 dimensions using Multidimensional Scaling (Fig. 4I, S12, Supplementary Video 1). This reorganization occurs without the encoding of additional stimulus information since individual stimuli are equally decodable in the presence or absence of inference behavior. This change in geometry relies on the encoding of context in an abstract format and is unique to hippocampus. In contrast, we found no systematic reorganization of stimulus representations in VTC.

### The tuning changes of hippocampal neurons that reflect the geometry changes

We next examined what aspects of neuronal activity changed in the hippocampus to give rise to the abstract neural representations that we observed (i.e. the representations with elevated CCGP). We considered the following non-mutually exclusive possibilities (Fig. 5A-D). (i) The distances between conditions in state space could increase (Fig. 5A vs. Fig. 5B), either as a result of increased firing rate of variable-coding neurons, an increase in the fraction of tuned neurons, or an increase in the depth of tuning of these neurons. (ii) The variance of the population response projected along the coding direction could decrease (Fig. 5C). (iii) Parallelism could increase due to increases in the consistency of firing rate modulation in response to one variable over values of another (Fig. 5D).

**Figure 5.**
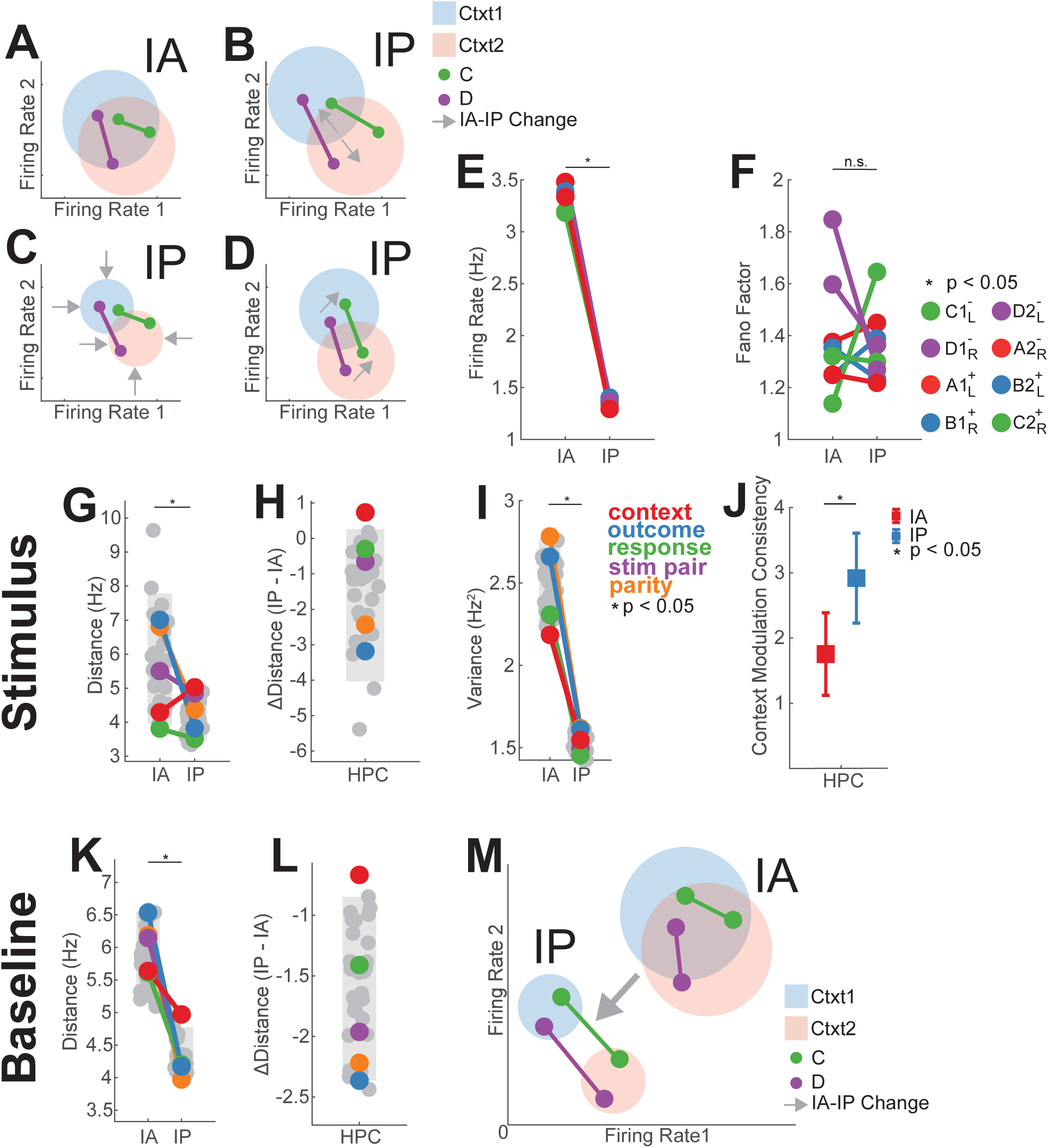
Firing rate properties underlying the observed changes of population-level hippocampal neural geometry. **(A-D)** Illustration of different hypothesized firing rate pattern changes that could give rise to the observed population level geometry changes. The four different hypotheses are illustrated with two stimuli (C and D), each present in two different contexts (blue and red). Condition responses are defined by the firing rates of two hypothetical neurons. The solid-colored points represent the condition average for each stimulus and the larger shaded circles represent the trial-by-trial variation of the two neurons for each context. Gray arrows signify changes that have occurred in IP plots **(B-D)** relative to the IA plot **(A)**. These response of the neurons to the stimuli during IA **(A)** can be shaped by **(B)** increasing the distance between the context centroids, **(C)** decreasing the variance along the coding direction in the absence of changes in distance, or **(D)** straightening the neural responses without changing distances/variances so that the geometry becomes more orthogonalized. **(E)** Changes in hippocampal firing rate from IA to IP. Points correspond to the average firing rate over neurons in the HPC for each of 8 unique task conditions, and are colored according to stimulus identity in that condition (e.g. task condition *C*1_L_^-^ describes: stimulus *C*, context 1, outcome −, response *L*). Neuronal firing rates were lower during IP compared to IA sessions (*p*_RS_ = 8.3×10^-5^, RankSum over conditions). **(F)** Same as **(E)**, but for condition-wise fano factors. Fano factor (FF) here is computed as the ratio of the condition-wise variance and the condition-averaged firing rate, computed by neuron and averaged over neurons. Points correspond to average FF over all hippocampal neurons. There was no significant difference (RankSum over conditions between IA and IP (*p*_RS_ = 0.99)). **(G)** Population distances between centroids for all 35 balanced dichotomies. The colored connected points represent distances for the named dichotomies indicated in the legend to the right. Gray bars indicate the 5^th^-95^th^ percentile of the geometric null distribution. Across all dichotomies, distances decreases from IA to IP (*p*_RS_ = 2.9×10^-8^, RankSum over dichotomies). **(H)** Context alone is the only dichotomy whose distance significantly increases from IA to IP (red, *p*_fJDist_ = 0.040). The null distribution shown H is the distribution of differences between the IP and IA null distributions shown in. **(I)** Average variance projected along the coding direction decreased on average between IA and IP sessions (*p*_RS_ = 6.5×10^-13^). Variance was computed in a cross-validated manner (see methods) resulting in a distribution of trial-by-trial population activity along each dichotomy coding direction from which the coding variance was computed. The 5^th^-95^th^ percentile of the geometric null distribution was also used here for the null distribution. **(J)** Change in the consistency of context-modulation for stimuli averaged over all neurons in HPC. Greater context modulation consistency for individual neurons results in greater parallelism score for context at the population level. HPC neurons on average exhibit a significant increase in context modulation consistency between IA and IP sessions (*p*_RS_ = 0.0039) during the stimulus period. **(K,L)** Same as **(G,H)**, but for spike counts during the baseline period and grouping trials by task state of the previous trial. Distance was significantly reduced across all dichotomies (**K**, *p*_RS_ = 6.4×10^-1^, RankSum over dichotomies) and context alone exhibits a distance reduction that is smaller than would be expected by chance (**L,** red, *p*_fJDist_ = 0.027) **(M)** Illustration of implementational changes to neural state space using the conventions introduced in **(A-D)**. We find that, when comparing IA with IP sessions, that (i) context dichotomy distance increased (indicated by the increased distance between the red and blue shaded circles), (ii) variance decreased due to a reduction in firing rate (indicated by decreased shaded circle radius and movement towards the origin of state space), and (iii) an increase in the consistency of stimulus modulation across contexts (indicated by lines becoming parallel).

We first examined whether mean firing rates across all recorded neurons differed between IA and IP sessions in the HPC. The firing rate across conditions decreased from 3.37±0.13 to 1.36±0.03 Hz, a 60% reduction on average during the stimulus period (Fig. 5E, *p*_RS_ = 8.3×10^-5^). Firing rates were also reduced during the baseline period (3.29±0.09 to 1.38±0.02 Hz, 58% reduction, Fig. S14G). This firing rate reduction was unique to the HPC, with every other recorded region exhibiting no significant differences or increases in firing rate between IA and IP during the stimulus (Fig. S13C) and baseline periods (Fig. S14H).

The firing rate reduction led to a decrease in the average distance between class centroids (separation) across all dichotomies in IP (5.77 ± 0.22 to 4.17 ± 0.07 Hz, *p*_RS_ = 2.9×10^-8^, Fig. 5G). However, the centroid distance for a single dichotomy, the context dichotomy, increased from IA to IP (4.3 vs 5.0 Hz, *p*_lA_ = 0.87, *p*_lP_ = 0.076, *p*_fJDist_ = 0.040, Fig. 5G,H,S13G). In fact, context was the dichotomy with the largest change in distance in firing rate space when comparing the IP and IA conditions (Fig. 5H). This isolated significant rise in context separability was not seen in any of the other recorded areas during the stimulus period (Fig. S13A,B). Similarly, during the baseline period, the centroids for context moved together the fewest in the HPC (5.6 vs 5.0 Hz, *p*_lA_ = 0. 68, *p*_lP_ = 0.0007, *p*_fJDis_ = 0.027, Fig. 5K,L) despite the significant decrease in distance over all dichotomies that was also observed here due to the firing rate reduction (5.85±0.08 to 4.25±0.04 Hz, *p*_RS_ = 6.5×10^-13^, Fig. 5K).

Next, we assessed changes in the variability of the population response along the coding direction of each dichotomy. The variance along the coding direction of neuronal responses in the HPC decreased for all dichotomies in IP when compared to IA sessions during both the stimulus period (2.51 ± 0.16 vs. 1.53 ± 0.06, *p*_RS_ = 6.5×10^-13^, Fig. 5I) and the baseline period (2.49 ± 0.09 vs. 1.58 ± 0.02, *p*_RS_ = 6.5×10^-13^, Fig. S14A,B). However, this decrease could be a simple consequence of the reduction in firing rates under the assumption of Poisson statistics. We conducted a condition-wise Fano-factor analysis to assess whether the variance reduction was beyond that expected for a the reduction in firing rates. This analysis revealed no significant differences in Fano factors between IA and IP sessions during the stimulus period (1.39±0.22 vs 1.36±0.14, *p*_RS_ = 0.99, Fig. 5F) and the baseline period (1.61±0.26 vs 1.45±0.11, *p*_RS_ = 0.19). Together, these two findings suggest that the decrease in variance along dichotomy coding directions is explained by the decreases in firing rate.

Though the increase in distances between dichotomy centroids for context appears to be a distributed, population-level phenomenon (see Fig. S5A-F), we sought to determine if a signature of this increase could be detected in the tuning of individual neurons (Supplement S.5). We found that the proportion of neurons exhibiting univariate context tuning increased from IA to IP sessions, thus partially explaining the increased representational distance. However, the hippocampal population geometry did not exclusively rely on these neurons because after excluding all neurons with significant univariate coding, we found no qualitative change in the population geometry (Fig. S5).

Finally, we examined the tuning of individual neurons to investigate what gave rise to the increases in parallelism for context across stimuli that we observed (see Fig. S11E-H for examples). A stimulus-tuned neuron also modulated by context could do so consistently across all stimuli (e.g. firing rate increased for all stimuli), or inconsistently (e.g. firing rate increased for some stimuli and decreased for others). Neurons consistently modulated by context would increase the context parallelism. Thus, we quantified the consistency in the direction with which stimulus representations were modulated (firing rate increased or decreased) across contexts for stimulus-identity tuned neurons. Context modulation consistency is computed for each neuron, and can take on values between 0 and 4, with 0 indicating no consistency in modulation and 4 indicating all stimuli exhibit the same firing rate modulation direction between contexts (see Methods for details). We find a significant increase in the consistency of context modulation in HPC from IA to IP sessions (Fig. 5J,11I, 1.8±0.2 vs 2.9±0.3, *p*_RS_ = 0.0049). This effect was specific to HPC: in VTC, this metric decrease significantly (Fig. S11I, 2.6±0.3 vs 1.6±0.2, *p*_RS_ = 0.0039). These findings indicate that, for HPC, context parallelism arises in part due to an increase in the consistency with which the firing rates of stimulus-tuned neurons are modulated by context.

The changes in neural state space responses for HPC are summarized in Fig. 5M, and feature aspects of our previous hypotheses: (i) condition averages for context increase in separation despite relaxing towards the origin (decrease in firing rate), (ii) are accompanied by decreases in variance along the coding direction, and (iii) neurons become increasingly consistent (parallel) in their modulation across stimulus and context dimensions. Together, these changes explain the implementation of the context coding dimension in the hippocampal representation and how it is able to emerge as a simultaneous, linearly encoded variable alongside stimulus identity.

### Context representations outside of the Hippocampus

The only other area of the brain that we examined in which we found a representation of latent context that correlated with inference behavior was in the dACC, but only during the baseline but not the stimulus period (Supplement S.6). Interestingly, context emerged in the task representation through a different implementation strategy, namely an increase in firing rates rather than a decrease, as was observed in HPC.

### Verbal instruction induces the representation of context in an abstract format

In all analyses discussed thus far, we compared sessions in which patients performed inference (IP) with those in which patients did not (IA), without regard to how patients transitioned from the IA to the IP group. We provided verbal instructions detailing the latent task structure following the first session (Fig. 6, inset) to all patients, allowing us to examine whether instructions lead to changes in neural representations. As shown above, patients were divided into three types based on behavior: those who exhibited inference behavior in the very first session (S1 group), those who did so after being given verbal instructions (IS group), and those who did not perform inference even after being provided with verbal instructions (IU group). We next compared the neural representation of context between these three groups of patients.

**Figure 6.**
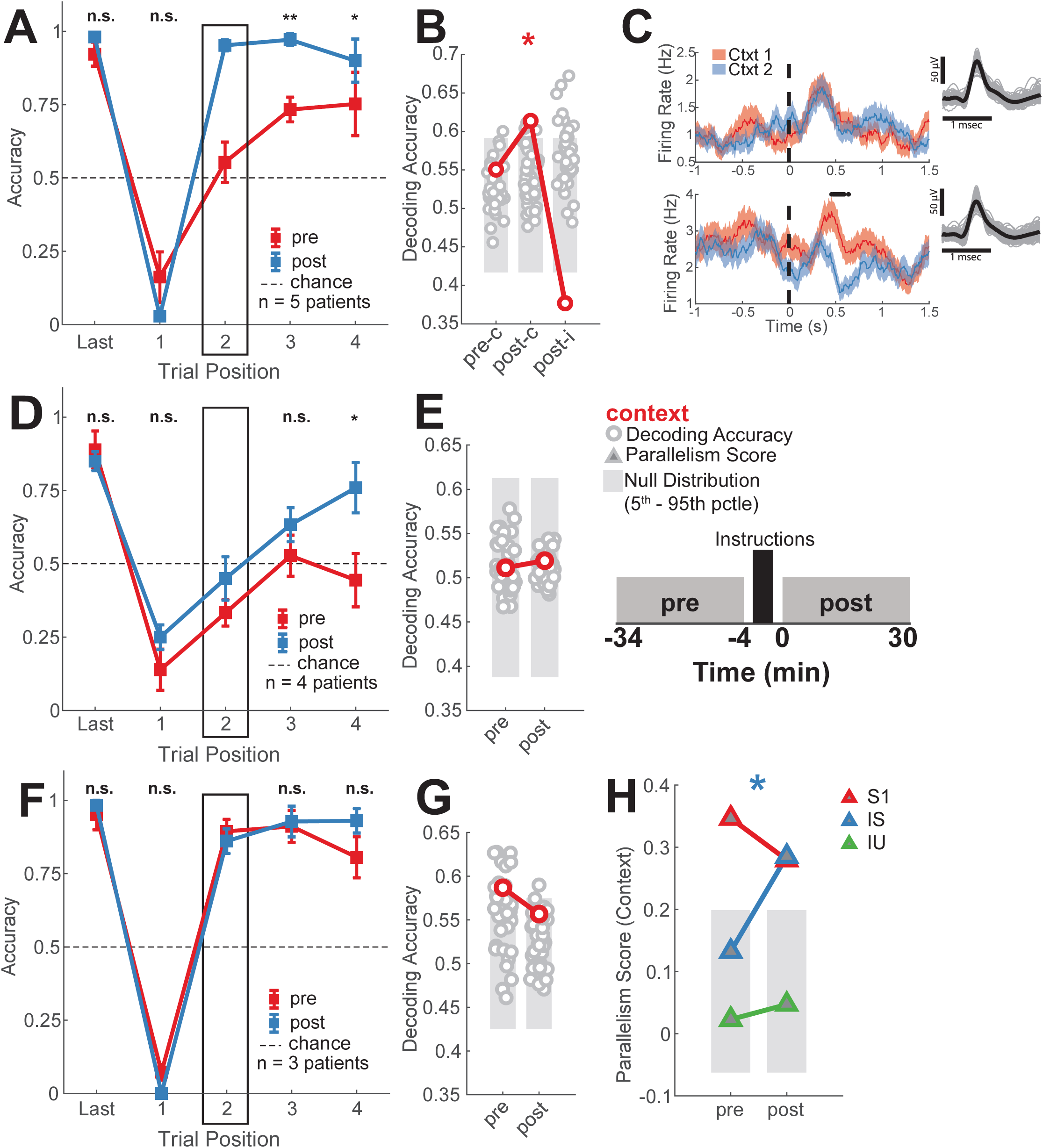
Abstract hippocampal representation of context is present following successful verbal instructions about latent context. **(A,D,F)** Behavioral performance shown separately for pre-instruction and post-instruction sessions (pre and post). **(A)** Inference performance for the instruction-successful (IS) patient group (5 patients, 10 sessions). **(C)** Example hippocampal neuron with univariate context encoding in the post (bottom) but not the pre instruction (top) session. (one-way ANOVA, *p*_pre_ = 0.40, *p*_post_ = 0.010). **(D)** Inference performance for the instruction-unsuccessful (IU) patient group (4 patients, 8 sessions). **(F)** Inference performance for the session one inference (S1) patient group (3 patients, 6 sessions). **(B,E,G)** Encoding of context in the stimulus period in pre-and post-instruction sessions during pre-instruction correct trials (pre-c), post-instruction correct trials (post-c) and post-instruction incorrect trials (post-i). The first trial following a switch is excluded from this analysis. **(E, inset)** Schematic of the recording procedure, showing a pre- and post-instruction session shaded in gray (30 min duration), with a 4 minute inter-session break (mean duration = 241 s, range 102-524s) during which instructions detailing task structure were provided. **(B)** Context emerges as significantly decodable post-instructions but not pre-instructions in the IS group in a task-performance-dependent manner (*p*_pre-c_ = 0.17, *p*_post_ = 0.016, *p*_RS_ = 3.1×10^-19^, *p*_post-_ = 0.99). **(E)** Context is not significantly decodable in the IU group neither pre- nor post-instructions (*p*_pre_ = 0.44, *p*_post_ = 0.42). **(G)** Context is decodable both pre- and post-instructions in the S1 group (*p*_pre_ = 0.014, *p*_post_ = 0.17). **(I)** Summary of changes in PS for context for all three subject groups (S1 red, IS blue, IU green). Neuron counts are sub-sampled to match across all groups so that PS values are directly comparable. Significant increase in context PS from pre to post is indicated for the IS group (*p*_lS,pre_ = 0.20, *p*_lS,post_ = 0.0028), but not for the IU (*p*_lU,pre/post_ < 0.5) and S1 (*p*_S1,pre/post_ < 0.005) groups.

The instructions provided to these three groups were identical, and all patients acknowledged receipt of the instructions. All included patients exhibited high performance on non-inference trials both before and after being given instructions, indicating that they understood the task and learned the SRO maps. The principal difference between the IS (Fig. 6A, S15A,B) and IU (Fig. 6D, S15C,D) groups is therefore their ability to perform inference following the verbal instructions, with both groups performing the task accurately otherwise. The S1 group, on the other hand, exhibited above-chance inference performance during both pre and post-instruction sessions (Fig. 6F, S15E,F).

In the IS group, context was decodable in the HPC during the stimulus period on correct trials in the session following the verbal instructions (Fig. 6B, *p*_pre-c_ = 0.17, *p*_post-_ = 0.016, *p*_RS_ = 3.1×10^-19^). This representation of context was in an abstract format, as indicated by significant increases in both CCGP (Fig. S16A; *p*_pre_ = 0.28, *p*_post_ = 0.047, *p*_RS_ = 8.4×10^-1^) and PS (Fig. S16B; *p*_pre-c_ = 0.023, *p*_post-_ = 1.2×10^-6^). Successful performance in the task was associated with context being represented abstractly in HPC, as both the decodability (Fig. 6B, *p*_post-_ = 0.99, post-c vs. post-i, *p*_RS_ = 4.3×10^-2^) and PS (Fig. S16B, *p*_post-i_ = 1.1×10^-4^) of context decreased significantly on incorrect trials in the post-instruction session. Context was also encoded in an abstract format during the baseline period in the same performance dependent manner as context in the stimulus period (Fig. S16C-E). At the single neuron level, this effect can be appreciated by an increase in the proportion of neurons that are significantly linearly tuned to context (*p* < 0.05, one-way ANOVA for context) during both the stimulus (8% (6/75 neurons) vs. 18% (17/93 neurons), *p* = 0.027) and baseline (7% (5/75 neurons) vs. 16% (15/93 neurons), *p* = 0.029) periods in the post-instruction session compared to the pre-instruction session (Fig. 6C shows an example). Thus, the ability of IS group patients to perform inference following instructions was associated with the rapid emergence of an abstract context variable in their hippocampus.

In contrast to the IS group, in patients in the IU group, context was not encoded by HPC neurons neither during the stimulus (Fig. 6E, S16F,G, all *p*_pre/post_ > 0.05) nor the baseline (Fig. S16H-J all *p*_pre/post_ > 0.05) periods in the post-instruction session. Furthermore, there was no significant change in tuning to context at the single-neuron level in the hippocampus of patients in the IU group both during the stimulus period (6% pre vs 6% post, *p* = 0.41) and the baseline period (8% pre vs 5% post, *p* = 0.27). These data indicate that receiving verbal instructions describing the latent context alone is insufficient to generate an abstract context representation in the hippocampus. Instead, context was only represented abstractly in the subset of subjects that performed inference following the instructions.

For the S1 patient group, context was decodable already during the pre-instruction session (Fig. 6G, *p*_pre_ = 0.014), and dropped slightly below significance (*p*_post_ = 0.17) during the post-session, likely due to the small number of patients and neurons present in this analysis. The CCGP was not significance (Fig. S16K, *p*_pre_ = 0.21, *p*_post_ = 0.31), but the PS in both the pre and post-sessions was highly significant and near the top of the dichotomy rank order in both cases (Fig. S16L, *p*_pre_ = 1.5×10^-9^, *p*_post_ = 1.7×10^-6^). This finding suggests that the context variable these patients learned experientially during the pre-instruction session was in an abstract format as assessed by PS but not CCGP, and that receiving the instructions did not significantly alter the behavior or the neural representation of context. A similar trend was observed with the baseline context representation for these patients (Fig. S16M-O). Note that such discrepancies between CCGP and PS are not unexpected since, when less data is available (fewer neurons, decreased firing rates), single trial measures (Decoding, CCGP) become less sensitive to representational structure than measures that operate on condition averages (PS).

We also examined firing rate changes of HPC neurons separately for the IS, IU, and S1 groups (Fig. S13D). This analysis revealed significant reductions in firing rate across all conditions from pre- to post-sessions for the IS patients alone (-0.39 ± 0.15 Hz, *p*_lS_ = 1.4×10^-4^), confirming that reductions in hippocampal firing rate in the same neurons recorded across adjacent sessions were associated with increases in inference performance. HPC neurons from IU patients exhibited an increase in firing rate (0.10 ± 0.04 Hz, *p*_lU_ = 1.2×10^-4^) and S1 firing rates did not significantly change (0.06 ± 0.08 Hz, *p*_S1_ = 0.08).

Lastly, we compared the geometry of the context representations formed by each of these patient groups using the PS (balancing number of neurons, see methods). PS for context increased significantly in the IU group, from levels not different from chance during pre-instructions (*p*_lS,pre_ = 0.20, *p*_lU,pre_ = 0.58, Fig. 6H) to a level comparable to S1 post-instructions (*p*_lS,post_ = 0.0028, *p*_S1,post_ = 0.0035, Fig. 6H). PS in the S1 group, on the other hand, did not change significantly and was higher than chance already in the pre-instruction group. This finding suggests that HPC neurons in the S1 group carried an abstract representation of context before receiving high-level instructions, and retained that geometry after receiving instructions. Hippocampal neurons in the IS group, on the other hand, did not encode an abstract representation of context before receiving instructions. However, upon receiving instructions, subjects in the IS group could perform inference, and neurons in their HPC started to encode a task representation whose geometry resembled that of the S1 group. This result indicates that a similar representational geometry can be constructed through either experience or instruction. Lastly, subjects in the IU group could not leverage the information provided in the instructions to perform inference, and accordingly, their hippocampi never encoded an abstract representation of context.

## Discussion

The ability to perform inference in our task was associated with the hippocampus forming an abstract representation of the environment. This representation encoded latent context and stimulus identity in approximately orthogonal subspaces, was behaviorally relevant on the level of individual trials, and emerged with learning. To implement this representation, the context coding directions for different visual stimuli became more parallel, the distance between contexts in neural state space increased, and the overall variance in firing was reduced due to a reduction in mean firing rates. This representation could emerge quickly, with some patients spontaneously learning the latent task structure during their first session and others exhibiting abstract representations within minutes of receiving verbal instructions explaining the task structure despite having no previous experience before the data shown here was recorded. Abstract representations of context and stimulus identity following stimulus onset were only present in the hippocampus and not in the other brain areas we examined. Together, this data reveals that hippocampal population codes can be restructured by learning and verbal instructions within minutes to support inference in a new task.

How can a neural or biological network efficiently encode multiple variables simultaneously^14,28^? One solution is to encode variables in an abstract format so they can be re-used in novel situations to facilitate generalization and compositionality^21,29–33^. Here, we show that in the human brain, such a disentangled representation emerged as a function of learning to perform inference in our task. The format by which latent context and stimulus identity was represented was predictive of the ability to perform behavioral generalization that relies on contextual inference. Crucially, patients performed well on non-inference trials in all sessions included in the analysis, indicating that they understood the task and successfully learned the stimulus-response associations in both contexts. Therefore, the difference between the inference present and absent sessions was only in whether they performed inference following the covert context switch. For those sessions where patients did not perform inference, there was no systematic relationship between context coding vectors across stimuli. For sessions where patients performed inference, there was alignment of the context coding direction across stimuli (making them parallel), indicating that the context variable had been disentangled from the stimulus identity variable in the hippocampi of these patients. As a result, the two variables became disentangled, thereby allowing for generalization.

Inferential reasoning is thought to rely on cognitive maps, which have been observed in the hippocampus and other parts of the brain^15,34–38^. Cognitive maps are thought to underlie inferential reasoning in various complex cognitive and spatial domains^3,10,34,35,39,40^. However, little is known about how maps for cognitive spaces emerge at the cellular level in the human brain as a function of learning. Here, we show that a cognitive map that organizes stimulus identity and latent context in an ordered manner emerges in the hippocampus. The cognitive map emerges because task states in one context, indexed by stimulus identity, become systematically related to the corresponding task states in the other context through a dedicated context coding direction that is disentangled from stimulus identity. Furthermore, the relational codes between task states (stimuli) in each context are preserved across contexts.

However, hippocampal cognitive maps observed in other studies are often different from those that we observed. Indeed, the encoded variables are often observed to non-linearly interact, which is a signature of high dimensional representations. These representations are believed to be the result of a decorrelation of the neural representations (recoding) that is aimed at maximizing memory capacity^41–43^. This form of pre-processing leads to widely observed response properties, like those of place cells^44^. However, there is some evidence of hippocampal neurons that encode one task variable independently of others^17,45–51^. In these studies, no correspondence was shown between different representational geometries in the hippocampus and differences in behavior. Here, the task representations generated when patients cannot perform inference (but can still perform the task) are systematically different from the abstract hippocampal representations of context and stimulus identity that correlate with inference behavior^14^. Finally, it is important to stress that we also observed an increase in the shattering dimensionality (SD). High SD can be compatible with the low dimensionality of disentangled representations^14,51^.

We found that there are other brain regions where stimulus identity is encoded, but does not undergo any major reorganization as a function of learning to perform inference. This is the case in the ventral temporal cortex, a region analogous to macaque IT cortex, in which neurons construct a high-level representation of visual stimuli^52–54^. Some studies conducting unit recordings in this general region in humans show that neurons exhibit strong tuning to stimulus identity ^27,55,56^. We similarly find that VTC neurons encode visual stimulus identity. However, these responses were not modulated by latent context in a systematic manner. As a result, despite being decodable for individual stimulus pairs, context was not represented in an abstract format. Rather, in VTC, context was only weakly decodable for a subset of the stimuli, context decodability did not change between IA and IP sessions (Fig. S11B,C), and stimulus identity geometry was not reorganized relative to context in IP sessions (Fig. 4D,E). Our study therefore shows that disentangled context-stimulus representations emerged in the hippocampus, but not in the upstream visually responsive region VTC.

The hippocampus was not unique in its representation of context. A weaker representation of context was also found in the dACC, but only during the baseline period. This finding is in line with work implicating the dACC in the representation of task rules and task sets ^57–62^. Following stimulus onset, however, dACC did not contain a representation of latent context (Fig. 2). In contrast, previous studies in tasks with explicitly cued context switches^19,20,63,64^ find that neurons in the medial frontal cortex (dACC and preSMA) are tuned to task context following stimulus onset. We hypothesize that this might be due to differences in task demands: in our task, context switches were uncued and had to be inferred from outcomes. It remains an open question to examine whether cued vs. inferred context switches engage different mechanisms of switching between contexts and/or different context encoding schemes in the hippocampus.

The focus of our study was to examine how representations of context, stimulus identity, response, and predicted outcome change as a function of learning in the human brain. In a prior study in macaques^14^, the representation of the same variables in very well trained animals was examined in HPC, dlPFC, and ACC in a similar task. Several notable differences exist between the two studies. First, context was encoded in an abstract format at baseline and was decodable after stimulus onset in all three brain areas examined in the macaques. In contrast, in humans, context is only strongly decodable in the HPC. We hypothesize that the wide-spread encoding of context in the macaque study was due to the extensive training the animals received before recordings commenced. In contrast, our patients had no prior task experience. It is possible that early on during learning, latent context representations are present only in the hippocampus, and are propagated to cortex (dACC) with extensive task experience. This hypothesis is supported by prominent direct and indirect projections from the hippocampus to dACC in primates^65–67^, and flexible, context-dependent interactions between medial frontal cortical neurons and hippocampal outputs^19,68^. Second, the human hippocampus exhibited abstract stimulus representations, unlike the abstract response or “choice” representation in the macaques. Notably, the abstract stimulus pair and response dichotomies are constructed such that high CCGP for one will necessarily lead to below-chance CCGP for the other, which was indeed the case for both our study (high stim pair, low response) and the primate study (low stim pair, high response). One potential reason for these differences is a species difference: human HPC neurons are strongly modulated by the identity and semantic category of presented images^19,24,69–71^, making it natural to organize representations of context relative to this existing representation. Similarly, representations of choices are not prominent in the human HPC^19^. Another potential reason is a difference in task construction: our task employed semantically identifiable images, whereas the prior experiment with macaques used fractals. Third, unlike macaque HPC, human HPC did not encode predicted outcome. We note that in our task, outcome prediction was not necessary to perform the task because context switches were signaled by the accuracy of the response (correct or incorrect), which was independent of predicted outcome received for a correct response. Furthermore, all possible task-states were uniquely indexable using stimulus identity and context, rendering outcome prediction representation unnecessary for unambiguously defining the current task state. Finally, another possibility is that the reward for the macaques (juice volume) was more motivationally salient than the small monetary reward (25¢ or 5¢) patients received. We hypothesize that these reasons obviated the need for a predictive representation of outcome to complete the task in our patients. It remains an important open question whether representations similar to those seen in macaques emerge in the other brain areas we examined following extensive training. Our data indicates that on short experiential timescales, the human hippocampus generates a representation that encodes the minimum set of variables required to solve the task at hand.

In our study, verbal instructions resulted in changes in hippocampal task representations that correlated with behavioral changes. The emergence of this representation in the session immediately following the instructions in the IS group is correlated with their newfound ability to perform inference and suggests that hippocampal representations can be modified on the timescale of minutes through verbal instructions. This change in representation is qualitatively different from the standard neurophysiological approach of studying the emergence of a “learning set”, wherein a low-dimensional representation of abstract task structure emerges slowly over days through trial-and-error learning ^46,72,73^. Our finding of similar representational structure in the hippocampus in subjects who learned spontaneously and those who only learned after receiving verbal instructions suggests that both ways of learning can potentially lead to the same solution in terms of the underlying neural representation. In complex, high-dimensional environments, learning abstract representations through trial and error becomes exponentially costly (the curse of dimensionality), and instructions can be used to steer attention towards previously undiscovered latent structure that can be explicitly represented and utilized for behavior. The process of instruction-dependent restructuring of hippocampal representations is likely cortical-dependent, given the role of the cortex in language comprehension, but the exact mechanism by which this process occurs remains to be explored^30,74^. Our findings suggest that when high-level instructions successfully alter behavior, the underlying neural representation is rapidly modified to resemble one learned through experience.

## Acknowledgements

We thank R. Adolphs for advice and support throughout all stages of the project and the members of the labs of R. Adolphs, U. Rutishauser, and M. Meister for discussion. We thank all subjects and their families for their participation and the staff and physisicans of the Cedars-Sinai and Toronto Western Epilepsy Monitoring Units for their support.

## Author contributions

Conceptualization: J.M., U.R., C.D.S., S.F., Task design: J.M. and D.L.K., Data collection: J.M., H.S.C, and A.R.C., Data analysis: H.S.C. and J.M., Writing: H.S.C., U.R., and S.F., Clinical care and experiment facilitation: C.M.R. Surgeries: A.N.M. and T.A.V., Supervision: U.R. and S.F.

## Data availability statement

Data will be made available on public repositories such as OSF or DANDI upon acceptance, as we commonly do for our publications.

## Code availability statement

Example code to reproduce the results will be made available on public repositories such as Github upon acceptance.

## Methods

### Participants

The study participants were 17 adult patients who were implanted with depth electrodes for seizure monitoring as part of an evaluation for treatment for drug -resistant epilepsy (see Table 1). 14 were monitored at Cedars-Sinai Medical Center (CSMC) and the other 3 were monitored at Toronto Western Hospital (TWH). All patients provided informed consent and volunteered to participate in this study. All research protocols were approved by the institutional review boards of CSMC, TWH, and the California Institute of Technology.

### Psychophysical Task and Behavior

Participants performed a serial reversal learning task. There were two possible static stimulus-response-outcome (SRO) maps, each of which was active in one of the two possible contexts. Context was latent and switches between context were uncued. Each recording session consisted of 280-320 trials grouped into 10-16 blocks of variable size (15-32 trials/block) with block transitions corresponding to a change in the latent context. Each trial consisted of a blank baseline screen, stimulus presentation, speeded response from the participant, followed by feedback after a brief delay (Fig. 1A). Responses were either “left” or “right” in every trial. In each session, stimuli were four unique images, each chosen from a different semantic category (human, macaque, fruit, car). If a patient performed multiple sessions, new images not seen before by the patient were chosen for each session. The task was implemented in MATLAB (The Mathworks, Inc., Natick, MA) using PsychToolbox-3^75^. Images were presented on a laptop positioned in front of the patient and subtended approximately 10 degrees of visual arc (300 px^2^, 1024×768 screen resolution, 15.6 inch (40 cm) monitor, 50 cm viewing distance). Patients provided responses using a binary response box (RB-844, Cedrus Inc.).

Receipt of reward in a given trial was contingent on the accuracy of the response provided. In each trial, either a high or low reward (25¢ or 5¢) was given if the response was correct, and no reward (0¢) if incorrect. Whether a given trial resulted in high or low reward if the response was correct was determined by the fixed SRO map (see Fig. 1C). Stimulus-response associations were constructed such that two out of four images (randomly selected) were assigned one response and the other two images were assigned the other (e.g. human and fruit = left, macaque and car = right). Thus, in each context, each stimulus was uniquely specified by a combination of its correct response (left/right) and reward value (high/low). Crucially, the SRO maps of the two possible contexts were constructed so that they were the opposite of each other from the point of view of the associated response (Fig. 1C). To fully orthogonalize also associated reward, half of the reward values stayed the same and the others switched. This structured relationship of stimuli across contexts led to the full orthogonalization of the response, context, and reward variables (Fig. 1B-C). Crucially, the stimulus-response map inversion across contexts provided the opportunity for patients to perform inferential reasoning about the current state of the SRO map, and therefore the latent context.

Since rewards were provided deterministically, participants could switch context upon receiving a single error. Therefore, if patients performed inference, they should be able to respond correctly after receiving a single error. The behavioral signature of inferential reasoning was thus the accuracy in the trials that occurred immediately after the first error trial. Specifically, we took a participant’s performance on the first instance of each of the three remaining stimuli in the new context is to measure a participants inference capabilities.

Patients completed multiple sessions of the task, in each of which new stimuli were chosen. After completion of the first session, the experimenter provided a standardized description of the latent contexts and SRO reversal to the patient (see below). These instructions were given regardless of how well the patient performed in the immediately preceding session. After this brief interlude, the participants completed the task again with a novel set of four stimuli.

#### Instructions given to patients

---------------------- Instruction set 1 (before first session) -----------------------

In this task, we will show you a series of images, 4 of them in total. Your objective is to learn the correct response for each image (either left or right). In the beginning, you will not know what the correct answer is, so take a guess. The correct answer for an image may occasionally change, so pay close attention. For every correct answer you will receive a reward of either 25 or 5 cents. For an incorrect answer you will receive 0 cents. This is real money that you will receive before you leave the hospital in the form of a gift card to your favorite place (ex. Starbucks). You will have the opportunity to take a break halfway through.

----------------------- Instruction set 2 (before second session) -----------------------

You may have noticed that some images have the same correct response and some images have the same reward. Even when the correct response changes, they usually change together. In this experiment, we are going to try a different strategy. Pay attention to which images go together (i.e. have the same correct response and similar reward). This should make it a lot easier to perform the task. To make the task a little more difficult, now the correct response for each image will change a little more frequently.

### Behavioral Control

We administered a control version of the task identical to the ‘first session’ described above to n=49 participants recruited on Amazon Mechanical Turk (MTurk). We then used this data to calibrate the difficulty of the task. A majority (∼75%) of the control subjects demonstrated proper inference performance, and the remaining 25% demonstrating slow updating of SROs after a context switch, consistent with a behavioral strategy where each stimulus is updated independently (see Fig. S1A).

### Electrophysiology

#### Electrode Placement and Recording

Extracellular electrophysiological recordings were conducted using microwires embedded within hybrid depth-electrodes (AdTech Medical Inc.). The patients we recruited for this study had electrodes implanted in at least the hippocampus, as well as in addition subsets of amygdala, dACC, pre-SMA, vmPFC, and VTC as determined by clinical needs (see Table 1). Implant locations were often bilateral but some patients only had unilateral implants as indicated by clinical needs. Broadband potentials (0.1Hz – 9kHz) were recorded continuously from every microwire at a sampling rate of 32kHz (ATLAS system, Neuralynx Inc.). All patients included in the study had well isolated single neuron(s) in at least one of the brain areas of interest.

#### Electrode Localization

Electrode localization was conducted using a combination of pre-operative MRI and post-operative CT using standard alignment procedures as previously described^19,63^. Electrode locations were co-registered to the to the MNI152-aligned CIT168 probabilistic atlas^76^ for standardized location reporting and visualization. Placement of electrodes in gray matter was confirmed through visual inspection of subject-specific CT/MRI alignment, and not through visualization on the atlas.

### Spike Detection and Sorting

Raw electric potentials were filtered with a zero-phase lag filter with a 300Hz-3kHz passband. Spikes were detected and sorted using the OSort software package^77^. All spike sorting outcomes were manually inspected and putative single-units were isolated and used in all subsequent analyses. We evaluated the quality of isolated neurons quantitatively using our standard set of metrics^69,78,79^ including proportion of inter-spike interval violations < 3ms, signal-to-noise ratio of the waveform, projection distance between pairs of isolated clusters, and isolation distance of each cluster relative to all other detected spikes.

### Selection of Neurons, Trials, and Analysis Periods

Activity of neurons was considered during two epochs throughout each trial: the baseline period (base), defined as -1s to 0s preceding stimulus onset on each trial, and the stimulus period (stim), defined as 0.2s to 1.2s following stimulus onset on each trial. Spikes were counted for every neuron on every trial during each of these two analysis periods. The resulting firing rate vectors were used for all encoding and decoding analyses. Tests of single-neuron selectivity were conducted using N-way ANOVAs with significance at P < 0.05, where N was either 2 for models of stim id (A, B, C, D) and context (1, 2), or 3 for models including outcome (High, Low), response (Left, Right), and context (1, 2). All variables were categorical, and all models were fit with all available interaction terms included.

### Population analysis – decoding

Single-trial population decoding analysis was performed on pseudo-populations of neurons assembled across all neurons recorded across all patients. We pooled across sessions within each anatomically specified recording area as described previously^19,20^. Decoding was conducted using support vector machines (SVM) with a linear kernel and L2 regularization as implement in matlab’s fitcsvm function. No hyperparameter optimization was performed. All decoding accuracies are reported for decoding accuracy for individual trials. Decoding accuracy is estimated out-of-sample using 5-fold cross-validation unless otherwise specified (e.g. cross-condition generalization). Many of the decoding analyses in this work consist of grouping sets of distinct task conditions into classes, then training an SVM to discriminate between those two groups of conditions. Neurons included in the analysis were required to have at least K correct trials of every unique condition in order to be included in the analysis (K = 15 trials unless otherwise stated). To construct the pseudopopulation, we then randomly sampled K trials from every unique condition and divided those trials into the groups required for the current decoding analysis for every neuron independently. Randomly sampling correct trials in this way allowed us to destroy noise-correlations that might create locally correlated sub-spaces from neurons recorded in the same area and session^19^.

To account for the variance in decoding performance that arose from this random sub-sampling procedure, all reported decoding accuracies are the average resulting from 1000 iterations of sub-sampling and decoder evaluation. A similar trial balancing and sub-sampling procedure was conducted for all analyses that report decoding accuracy on incorrect trials, but with K = 1 trial/condition required as incorrect for the neuron to be included in analysis. Various other analyses conducted throughout this work, including representation geometry measures, centroid distances, and coding direction variances, all rely on this procedure of balanced correct and incorrect trial sub-sampling, and averaging over 1000 iterations of the computed metric to study the relationships between task conditions in an unbiased manner. All reported values have been computed with this approach unless otherwise stated.

### Construction of Balanced Dichotomies

Our task has 8 possible states (Fig. 1B). We characterized how neurons represented this task space by assessing how a decoder could differentate between all possible “balanced dichotomies” of these 8 task conditions (Fig. 1B). The set of all possible balanced dichotomies is defined by all possible ways by which the 8 unique conditions can be split into two groups containing 4 of the conditions each (e.g. 4 points in context 1 vs 4 points in context 2 is the context dichotomy). There are 35 possible balanced dichotomies (nchoosek(8,4)/2). Some of the possible balanced dichotomies are easily interpretable because they correspond to variables that were manipulated in the task. We refer to these balanced dichotomies as the “named dichotomies”, which are: context, response, outcome, stimulus pair (stim pair), and parity. These dichotomies are shown individually in Fig. S2. The stim pair dichotomy corresponds to the grouping of stimuli for which the response is the same in either context (A&C vs. D& B; see Fig. S2). The parity dichotomy is the balanced dichotomy with the maximal non-linear interaction between the task variables (Fig. S2).

### Defining decoding difficulty of dichotomies

We quantify the relative degree of non-linear variable interactions needed by a neural population to classify a given dichotomy using a difficulty metric that rates dichotomies that require proximal task conditions to be placed on opposite sides of the decision boundary as more difficult. Note that proximity of task conditions in task space here is defined with respect to the variables that were manipulated to construct the task space. The conditions corresponding to (Response L, Outcome Low, Context 1) and (Response L, Outcome Low, Context 2) are proximal since their task specifications differ by a single variable (hamming distance 1) whereas (Response L, Outcome Low, Context 1) and (Response R, Outcome High, Context 2) are distal since their task specifications differ by all three variables (hamming distance 3). With this perspective, we can systematically grade the degree of non-linearity required to decode a given dichotomy with high accuracy as a function of the number of adjacent task conditions that are on opposite sides of the classification boundary for that dichotomy. For a set of 8 conditions specified by 3 binary variables, this corresponds to the number of adjacent vertices on the cube defined by the variables that are in opposing classes (See Fig. S7A). We define this number as the “difficulty” for a given dichotomy, and can compute it directly for every one of the 35 balanced dichotomies. The smallest realizable dichotomy difficulty is 4, and corresponds only to named dichotomies that align with the axis of one of the three binary variables used to specify the task space. The largest realizable dichotomy is 12, and this corresponds to the parity dichotomy since the dichotomy difficulty (number of adjacent conditions with opposing class membership) is maximized in this dichotomy by definition. All remaining dichotomies lie between these two extremes in difficulty, and computing average decoding accuracy over dichotomies of increasing difficulty gives a sensitive readout of the degree of non-linear task variable interaction present in a neural population.

### Geometric Analysis of Balanced Dichotomies

We used three measures to quantify the geometric structure of the neural representation^14^: shattering dimensionality (SD), cross-condition generalization performance (CCGP), and parallelism score (PS).

SD is defined as the average decoding accuracy across all balanced dichotomies. It is an index of the expressiveness of a representation, as representations with higher SD allow more dichotomies to be decoded. The content of a representation is assessed by considering which balanced dichotomies are individually decodable better than expected by chance.

CCGP assesses the extent to which training a decoder on one set of conditions generalized to decoding a separate set of conditions. Note that to compute CCGP, all trials from a set of conditions are held out from the training data, which is different from the “leave-one-out” type decoding used to estimate SD. The remaining held-in conditions are used to train the decoder, and performance is then evaluated on the held-out conditions (trial-by-trial performance). The CCGP for a given balanced dichotomy is the average over all possible 16 combinations of held-out conditions on either side of the dichotomy boundary. One of the 4 conditions on each side of the dichotomy are used for testing, whereas the remaining three on each side of the dichotomy are used for training. For each of the 16 possible train/test splits, the decoder is trained on all correct trials from the remaining six conditions, and performance is evaluated on the two held-out conditions.

PS assesses how coding directions for one variable are related to each other across values of other variables in a decoder agnostic manner. The PS is defined for every balanced dichotomy as the cosine of the angle between two coding vectors pointing from conditions in one class to conditions in the other for a given dichotomy. These vectors are computed by selecting four conditions (two on either side of the dichotomy), computing the normalized vector difference between the mean population response for each of the two pairs, then computing the cosine between said coding vectors.This procedure is repeated for all possible pairs of coding vectors, and the average over all cosines is reported. Since the correct way of “pairing” conditions on either side of the dichotomy is not known a-priori, we compute the cosine average for all possible configurations of pairing conditions on either side of the dichotomy, then report the PS as the maximum average cosine value over configurations.

### Null distribution for geometric measures

We used two approaches to construct null distributions for significance testing of the geometric measures SD, CCGP, and PS.

For the SD and decoding accuracy of individual dichotomies, the null distribution was constructed by shuffling trial labels between the two classes on either side of each dichotomy prior to training and testing the decoder. After shuffling the order of the trial labels, the identical procedures for training and testing were employed. This way of constructing the null distribution destroys the information content of the neural population while preserving single-neuron properties such as mean firing rate and variance.

For the CCGP and PS, we employed a geometric null distribution^14^. Prior to training, we randomly swapped the responses of pairs of neurons within a given condition. For example, for one task condition, all of neuron 1’s responses are assigned to neuron 2 and all of neuron 2’s responses are assigned to neuron 1, for another task condition, all of neuron 1’s responses are assigned to neuron 3, etc…). This way of randomly shuffling entire condition responses leads to the situation where neural population response statistics by-condition are held constant, but the systematic cross-condition relationships that exist for a given neuron are destroyed. This way of shuffling creates a maximally high dimensional representation, thereby establishing a conservative null distribution for the geometric measures CCGP and PS.

#### Analysis of Incorrect Trials

For determining decoding accuracy for trials in which subjects provided an incorrect response (“error trials”), decoders were trained and evaluated out of sample on all correct trials in IA and IP sessions (denoted as “IA-c” and “IP-c” trials respectively). The accuracy of the decoder was then evaluated on the left out incorrect trials in the IP sessions (denoted as “IP-i”" trials) that were balanced by task condition. Neurons from sessions without at least one incorrect trial for each of the 8 conditions were excluded. We did not estimate CCGP separately for correct and incorrect trials. The PS was estimated using only correct trials for IP-c and IA-c. For IP-i, parallelism was computed using one coding vector (difference between two conditions) from correct trials and one coding vector from incorrect trials. All other aspects of the PS calculation remained as described earlier. The very first trial after a context switch was excluded from analysis (it was incorrect but by design, as the subject cannot know when a context switch occurred).

#### Stimulus Identity Geometry Analysis (Fig. 4)

We repeated the geometric analysis described above for subsets of trials to examine specifically how the two variables context and stimulus interact with each other. To do so, we considered each possible pair of stimuli (AB, AC, AD, BC, BD, CD) separately. For each stimulus pair, we then examine the ability to decode and the structure of the underlying representation for two variables: stimulus identity (see Table 3) and context (see Table 4).

For stimulus identity, what is decoded is whether the stimulus identity is the first or second possible identity in each pair (i.e. “A vs. B” for the AB pair). Stimulus CCGP (Fig. 4B,D) is calculated by training a decoder to decide “A vs. B” in context 1 and testing the decoder in context 2 and vice-versa (the CCGP is the average between these two decoders). Stimulus PS (Fig. 4C,E) is the angle between the two coding vectors “A vs. B” in context 1 and 2.

For context, decoding accuracy is estimated by training two decoders to decide “Context 1 vs. Context 2” for each of the two stimuli in a stimulus pair. The reported decoding accuracy is the average between these two decoders (Fig. S11A,B). For example, for the stimulus pair AB, one such decoder each is trained for all “A” trials and all “B” trials. Context CCGP (Fig. 4G, S11C) is calculated by training a decoder to differentiate between Context 1 and 2 based on the trials in the first identity of the pair, and tested in the second pair and vice-versa. The reported Context CCGP value for a given stimulus pair is the average between the two. Similarly, context PS (Fig. 4H, S11D) is the angle between the two coding vectors Context 1 vs. Context 2 estimated separately for the first and second stimulus in a pair.

#### Distance/Variance Analysis (Fig. 5)

We computed a series of metrics to quantify aspects of the population response that changed between IA and IP sessions. We used (i) the firing rate, (ii) distance in neural state space between classes for balanced dichotomies and stimulus dichotomies (dichotomy distance), (iii) variance of neural spiking projected along the coding directions for those dichotomies (coding direction variance), and (iv) the condition-wise fano factor.

Firing rate (Fig. 5E) was the mean firing rate averaged across all neurons during the stimulus period, reported separately for correct trials of every unique task condition. Values reported during the baseline (Fig. S14G,H) are computed with an identical procedure using firing rates from before 1s prior to stimulus onset.

Dichotomy distance (Fig. 5G,H,K,L) was defined as the Euclidean distance in neural state space between the centroids of the two classes on either side of the decision boundary for that dichotomy. Centroids were computed by constructing the average response vector for each class using a balanced number of correct trials from every condition included in each class through a resampling procedure (described below). Null distributions reported for dichotomy distances are geometric null distributions.

Coding direction variance (Fig. 5I) was computed for a given balanced dichotomy by projecting individual held-out trials onto the coding vector of the decoder trained to differentiate between the two groups of the balanced dichotomy being evaluated. The coding direction was estimated by training a linear decoder on all trials except eight (one from each condition either side of the dichotomy). The vector of weights estimated by the decoder (one for each neuron) was normalized to unit magnitude to estimate the coding vector. The projection of the left out trial onto this coding vector was than calculated using the dot product. This process was repeated 1000 times, generating a distribution of single trial projections onto the coding vector for each dichotomy. The variance of the distribution of 1000 projected data point was then computed and reported as the variance for a given balanced dichotomy (Fig. 5I).

The condition-wise Fano factor (Fig. 5F) was computed separately for each neuron. We used all correct trials for a given balanced dichotomy to estimate the mean firing rate and standard deviation and then took the ratio between the two to calculate the Fano factor for each neuron. Reported fano factors are the average of all fano factors across all neurons from that area/behavioral condition. Fano factors are computed by-condition since grouping trials across conditions could lead to task variable coding (signal) contaminating the fano-factor measurement, which should ideally only reflect trial-by-trial variation around the mean for approximately poisson-distributed firing rates.

The context-modulation consistency (Fig. 5J) was also computed separately for each neuron. Context modulation consistency is the tendency for a neuron’s firing rate to shift consistently (increase or decrease) to encode context across stimuli. For each neuron, it was computed by deteriming the sign of the difference (+/-) between the mean firing rate for a given stimulus between the two contexts, and summing the number of stimuli that exhibit the same modulation (either increase or decrease) across the two contexts. This consistency can take on values between 0 (increase in firing rate to encode context for half of the stimuli, decrease in firing rate for the other half) and 4 (either increase or decrease in firing rate for all four stimuli).

#### Bootstrap Re-sampled Estimation of Measures and Null Distributions

All the measures described in the preceding sections were estimated using a trial and neuron-based re-sampling wmethod. This resampling strategy was used to assure that every measure reported is comparable between a set of conditions by assuring that the same number of neurons and data points are used to train and test classifiers. Metrics were re-computed 1000 times with resampling and all null distributions were computed with 1000 iterations of shuffling and re-computing. Plotted boundaries of null distributions correspond to the 5^th^ and 95^th^ percentiles as sestimated from the 1000 repetitions.

A single iteration of the re-sampling estimation procedure proceeds as follows. For all analyses that involved a comparison of a metric between two behavioral conditions (IA vs IP or pre vs. post instruction), the same number of neurons was included in both conditions by on a region by region basis. For a neuron to be included, at least 15 correct trials for each of the 8 unique task conditions had to exist (120 correct trials total). Across patients, the number of correct trials per condition varied: min = 10.9 ± 1.3 trials/condition, mean = 25.0 ± 0.6 trials/condition, max = 39.6 ± 1.2 trials/condition (mean ± s.e.m.). After identifying the neurons that met this inclusion criteria, an equal number were randomly sampled from both behavioral conditions. The number of considered neurons was set to the number of neurons available in the smallest group.

When constructing feature matrices for decoding, 15 trials were randomly selected from each unique condition that was included in the given analysis. Trial order was shuffled independently for every neuron within condition to destroy potential noise correlations between neurons that were simultaneously recorded. For decoding and SD, out-of-sample accuracy was estimated with 5-fold cross validation. For generalization analyses (CCGP), all trials were used in training since performance is evaluated on entirely held-out conditions. For vector-based measures (dichotomy distance, variance, PS), all trials in relevant conditions were used to compute condition centroids. In the case of variance estimation, all trials except one on either side of the dichotomy boundary were used to learn the coding axis, then the held-out trials were projected onto the coding axis. As previously stated, these procedures were repeated 1000 times with independent random seeds to ensure independent random sampling of neurons and trials across iterations.

