## Supplementary Information 1 for "Abstract representations emerge in human hippocampal neurons during inference behavior"

#### Supplementary Sections

##### S.1 Hippocampus encodes context abstractly at baseline during inference sessions.

We analyzed the baseline period preceding stimulus onset (Fig. 2, inset) to study the geometry of task representations that might persist from the previous trial (all labels were defined by the prior just completed trial for this analysis). Context was encoded as an abstract variable in the HPC in IP and not in IA sessions. Unlike the stim epoch, context was the only named dichotomy to emerge as significantly decodable in the HPC for IP sessions (Fig. 2D, red, IA vs. IP, $p_{RS} = 1.1x10^{-33}, p_{IA} = 0.35, p_{IP} = 6.4x10^{-5}$, S4B), indicating that any task condition information from the previous trial other than the context (i.e. outcome, response) was not significantly decodable from the population during this epoch. Context during baseline was also encoded abstractly as shown by the increase in CCGP (Fig. 2E, S4C, red, IA vs. IP, $p_{RS} = 2.4x10^{-34}$ , $p_{IA} = 0.45, p_{IP} = 7.7x10^{-6}$) and PS (Fig. S4D,E, red, $p_{IA} = 0.37, p_{IP} = 1.2x10^{-10}$). These results indicate that an abstract representation of context that persisted from the previous trial also emerged in the hippocampus in sessions where patients performed inference.

#### S.2 Hippocampal context representation does not rely on classically tuned neurons

To determine whether the geometric findings presented here arose as a consequence of strong classical (univariate linear) tuning, two ablation analyses were conducted in which all neurons that had significant univariate linear tuning to at least one task variable were excluded and all measures were re-computed. In the first analysis, neurons tuned to one or several of response, reward, or context (3-Way ANOVA, any main effect $p < 0.01$) were excluded. Of the 494 neurons in HPC, 41 were excluded for exhibiting classic tuning prior to recomputing all measures. Qualitatively, findings remained largely unchanged by the exclusion of this population of specialized neurons (Fig. S5A-E). The Parity dichotomy was no longer decodable above chance in IP sessions (Fig. S5A, orange, $p_{IP-c} = 0.056$), but the rise in SD (average over dichotomies) was still present (Fig. S5A, $p_{RS} = 0.0012$, Ranksum over dichotomies). A similar analysis was also conducted for the baseline representation of context, and again the findings qualitatively remained unchanged (Fig. S5F-J). Note that, although the decodability of context at baseline during IP sessions did decrease in significance when conducting the error trial analysis (Fig. S5I, red, $p_{IP-c} = 0.12$), it was still significantly increased compared to IA sessions ($p_{RS} = 5.7x10^{-19}$) and was significantly reduced in error trials ($p_{IP-i} = 0.37$, IP-c vs IP-i, $p_{RS} = 0.0002$). In the second analysis, neurons tuned to stimulus identity or context (2-Way ANOVA, any main effect $p < 0.01$) were excluded. In this analysis, 82 neurons were excluded, leading to a loss of decodability of stimulus dichotomies due to the removal of visually-selective neurons as expected (Fig. S5K-M). Context, however, remained decodable and present in an abstract format. Together, these analyses indicate that the abstract context representation in the hippocampus is a highly distributed variable whose geometry is not a simple consequence of strong univariate tuning of a small population of “context” neurons.

#### S.3 Hippocampal context and stimulus representations are not driven by SOZ neurons

To ensure that the reported findings were not influenced by pathology, in a further control analysis we excluded all HPC neurons (86/494) recorded from electrodes that were clinically identified to reside in medial temporal seizure onset zones (SOZs). Repeating our analyses on the neurons that remain revealed that results were qualitatively unchanged (Fig. S6), though the parallelism for context was now significantly above chance during IA sessions for both the stimulus (Fig. S6C, red, $p_{IA} = 3.5x10^{-7}$) and baseline (Fig. S6F, red, $p_{IA} = 0.0022$) periods. These findings suggests that neurons residing in hippocampal tissue with high disease burden do not meaningfully contribute to the task representation[17], thereby leading to marginally increased representation strength once these neurons are removed (note that the decodability and CCGP for context in IA are still not significantly different from chance, confirming our finding).

#### S.4 Hippocampal representations are not driven by differences in non-inference trial performance

To ensure that the reported geometric findings in the hippocampus did not arise due to non-inference performance differences between IA and IP sessions, an additional control analysis was performed where the non-inference trial performance of included sessions was distribution-matched between IA and IP. Pairs of sessions from IA and IP with at most 7.5% difference in non-inference trial performance were selected, prioritizing sessions with more hippocampal neurons. This matching process yielded 10 IA sessions (152 HPC neurons) and 10 IP sessions (187 HPC neurons) whose average non-inference performances did not statistically significantly differ (92.8% IA v.s. 94.7% IP, $p_{RS} = 0.58$, RankSum over sessions, Fig. S6M). All main geometric findings were recapitulated using this session split during both the stimulus (Fig. S6G-I) and baseline (Fig. S6J-L) periods, indicating that the learning-dependent changes in the hippocampal representational geometry we observe cannot be explained by differences in non-inference trial performance.

**S.5 Changes in hippocampal single-neuron tuning explain changes in representation with inference**

To determine if changes in univariate tuning could partially explain the observed changes in the hippocampal representation from IA to IP sessions, average tuning properties of single neurons were computed. Both the number of significant factors (main effects or interactions) per neuron (3-way ANOVA – Response, Context, Outcome, $p < 0.05$) and the tuning strength of neurons for those significant factors indexed by the log-p-value of those ANOVA factors were significantly elevated in IP compared to IA (Fig. S13E, $p_{RS} = 0.010$, S13H, $p_{RS} = 0.0089$). These effects were only observed in HPC, with either no change or a decrease in tuning observed in the other areas except for a significant increase in vmPFC (Fig. S13F,I). We also separately considered the linear and non-linear (interaction) terms for the 4 (Stimulus) x 2 (Context) ANOVA and found that, while the fraction of hippocampal neurons exhibiting significant ($p < 0.05$, Main Effect) stimulus identity tuning significantly decreased from IA to IP (Fig. S13J, 21.3% vs. 19.1% of neurons, $p = 0.002$), the fraction of neurons exhibiting significant context tuning increased from IA to IP (Fig. S13J, 9.5% vs. 15.7% of neurons, $p = 2.2x10^{-6}$), which could also partially explain the increase in centroid distance and resultant context decodability. Such "context neurons", however, cannot account for this effect fully as their removal from the pseudopopulation does not qualitatively alter the hippocampal population geometry (Fig. S5). As a control, this analysis was also performed for VTC neurons, in which we did not find a significant difference in the percentage of neurons with univariate context tuning (Fig. S13K).

#### S.6 Context representations outside of the hippocampus

The dorsal Anterior Cingulate Cortex (dACC) was the only other region to exhibit significant changes in its latent context representation as a function of patients' ability to perform inference. However, this context representation was limited to the baseline period, since geometric analysis for the stimulus period revealed an absence of significant context decodability and CCGP in IA and IP sessions (Fig. 2C, S3A). However, the PS for context in the dACC did increase significantly from IA to IP sessions (Fig. S3B, red, IA vs. IP, $p_{RS} = 2.4x10^{-19}, p_{IA} = 0.99, p_{IP} = 3.8x10^{-12}$), reflecting an increase in context parallelism in condition averages that was not detectable with the other metrics (which are based on single trial decoding).

During the baseline, however, context also emerged as the only decodable dichotomy in IP sessions in the dACC (Fig. S4A,H, red, $p_{IA} = 0.37, p_{IP} = 0.049$). We found this context variable was also represented in an abstract format, emerging as the dichotomy with the highest CCGP (Fig. S4F,I, red, $p_{IA} = 0.26, p_{IP} = 0.018$) and PS (Fig. S4G,J, red, $p_{IA} = 0.18, p_{IP} = 0.013$) in IP sessions, while being at chance for both metrics in IA sessions. As in HPC, the lack of decodability for previous trial outcome and response suggests that any variables encoded at earlier epochs during the previous trial (e.g. post-reply delay or outcome) were extinguished from the population by the onset time of the current baseline epoch.

Analysis of dichotomy centroid distances during the baseline period revealed context also emerged as the dichotomy with the greatest separation in the dACC in IP trials (3.9 vs. 5.7 Hz, $p_{IA} = 0.48, p_{IP} = 0.0065, p_{\Delta Dist} = 0.046$, Fig. S14C,D). However, this implementation of the context representation was achieved through significant increases in condition-wise firing rates (Fig. S14H, dACC IA vs IP, $p_{RS} = 0.049$) as opposed to the firing rate decrease observed in the hippocampus. Together, these analyses indicate that a weak, but nonetheless significant, representation of latent context encoded in an abstract format also emerged in the dACC, and that this variable was accommodated in the representation through a different implementational strategy.

#### Supplementary Figure Captions

**Figure S1. Task behavior and single-neuron responses across all recorded regions.**

**(A)** Task performance on individual sessions from 49 control subjects recruited through an online platform (Amazon MTurk). Accuracy is reported as an average for each subject over all non-inference trials (left) and inference trials (right). The horizontal gray dashed line corresponds to chance (50%). This task variant is equivalent to the first session of the task encountered by patients where they were given general instructions about learning stimulus-response mappings, but were not informed of the latent task structure. Subjects exhibited a variety of behaviors, with 46/49 subjects performing above chance on non-inference trials, indicating that the SRO maps were generally learnable despite the wide variation in performance on inference trials.

**(B)** Non-inference performance for context 1 is plotted against context 2 for each of the 36 sessions included in the analysis. Error bars correspond to SEM computed over blocks. The diagonal gray dashed line indicates identical block performance (y=x). The reported p-value is computed by paired t-test between the mean accuracies for Context 1 and Context 2 across all sessions.

**(C)** Same as (B), but with reaction time (RT), computed as time from stimulus onset to button press for every trial. Mean RT's are also computed by block.

**(D)** Truncated behavioral performance plot similar to Fig 1E. Plot shows performance on the last trial before the context switch, the first trial after the context switch, and for the first inference trial (Trial 2) averaged over all trials in each session (mean ± s.e.m. across sessions). Dashed line marks chance. Red and blue lines correspond to session performance before and after instructions detailing latent context are provided. This plot shows performance for the "instruction-successful" (IS) patient group – did not exhibit significant inference performance pre-instruction, but did exhibit inference performance post-instruction. Trial 2 pre/post performance was used to classify patients, so difference significance is not computed. All trials where pre/post performance difference was insignificant ($p > 0.05$) are shown with n.s.

**(E)** Same as (D), but for the "instruction-unsuccessful" (IU) patient group – did not exhibit significant inference performance during either pre-instruction or post-instruction sessions.

**(F)** Same as (D), but for the "session one inference" (S1) patient group – exhibited significant inference performance during both pre-instruction and post-instruction sessions.

**(G)** Normalized activity for all neurons recorded from the hippocampus is plotted as a heat map by region after computing the trial-averaged response to each unique condition (8 total, specified by unique Response-Context-Outcome combinations). Z-scored firing rates are computed from 0.2s to 1.2s after stimulus onset for every trial. Each row of the heat map corresponds to the activity of a single neuron, and columns correspond to each of the 8 conditions. Neurons are ordered such that adjacent rows (neurons) are maximally correlated in 8-dimensional condition response space. This approach would allow for modular tuning to visibly emerge in the heat map if groups of neurons were clustered in their response profiles. Clearly, the responses here are very diverse.

**(H)** Same as (G), but for amygdala.

**(I)** Same as (G), but for ventral temporal cortex.

**(J)** Same as (G), but for dorsal anterior cingulate cortex.

**(K)** Same as (G), but for pre-supplementary motor area.

**(L)** Same as (G), but for ventromedial prefrontal cortex.

**(M)** Percentage of neurons across all areas that exhibit tuning to each of the three binary variables manipulated in the experiment. Tuning was assessed by fitting either a 2x2x2 (Response-Context-Outcome) ANOVA for every individual neuron's firing rate during a 1s window during the stimulus presentation period. Significant neurons were counted as $p < 0.05$ for main effects (Linear) or interaction effects (Nonlinear) involving the stated variables (RxC = Response x Context interaction, RxO = Response x Outcome interaction, CxO = Context x Outcome interaction, RxCxO = Response x Context x Outcome interaction). Significance in the change of percentage of neurons exhibiting modulation to each factor is determined via z-test, where "*" indicates $p < 0.05$, "***" indicates $p < 0.005$, and "n.s." indicates "not significant".

**(N)** Same analysis as **(M)**, but for a 4x2 ANOVA for stimulus identity and context.

**(O)** Same analysis as **(M)**, but for a 4x2 ANOVA for stimulus identity and response.

Note: the corresponding analysis between stimulus identity and outcome cannot be conducted since those variables are correlated by task construction.

**Figure S2. Visual representation of all named balanced dichotomies.**

Named balanced dichotomies correspond to condition splits that have clearly interpretable meaning with respect to the construction of the task when evaluating the decodability or disentanglement (Cross Condition Generalization Performance, Parallelism Score) for that dichotomy. For example, the context dichotomy (top left), arises from assigning all conditions for context = 1 to one class and all conditions for which context = 2 to the other class. Performing binary classification on neural responses with trial labels arranged in this way corresponds to decoding context from the neural population. The specific assignment of class labels 1 and 2 is arbitrary, and inverting the labels still corresponds to the same meaning for the dichotomy. All named dichotomies shown here are color coded to reflect their value in all Shattering Dimensionality, CCGP, and PS plots, and this color code remains consistent throughout the paper whenever balanced dichotomies are considered.

The "stim pair" dichotomy corresponds to the special split of stimulus identities where the stimuli that have the same response in each context are grouped together (e.g. Stimuli A and C have Response L in Context 1 and Response R in Context 2, v.v. for Stimuli B and D). There are more balanced dichotomies that correspond to splits of stimulus identity (shown in Fig. S7). High decodability of any one of these balanced dichotomies reflects stimulus-id coding in the neural population.

The "parity" dichotomy is another special dichotomy that corresponds to the most difficult, or non-linear, dichotomy that can be constructed, and high decodability of this dichotomy is a signature of a high-dimensional representation. The term parity is in reference to the fact that, if task states were represented as 3-bit binary words where each bit corresponds to the value of the response, context, and outcome variable that describes the state (e.g. 000 for Left, Ctxt 1, Low, 111 for Right, Ctxt 2, High), then one class of the parity dichotomy corresponds to all states with an even number of ones, and the other class corresponds to all states with an odd number of ones. Note, for this dichotomy, no node of a given class shares an edge with another node of the same class. If one views the faces of the cube, one can see that the standard 2D XOR dichotomy between class 1 and 2 is present on every face. The notion of parity, can be generalized to arbitrarily many dimensions.

**Figure S3. Additional geometric analysis for hippocampus during stimulus processing.**

**(A)** Cross-condition generalization performance (CCGP) reported for other regions over balanced dichotomies. Each dot corresponds to the CCGP for a single dichotomy. The reported values are averages over 1000 repetitions of resampling of trials and neurons as described in the methods. For every region, the left column corresponds to inference absent (IA) sessions and the right column to inference present (IP sessions). Colored lines are drawn to connect IA and IP values for named dichotomies using the standard color-coding scheme. The gray background bars indicate the $5^{th}$ (bottom) to $95^{th}$ (top) percentile of the null distribution. Note that the null distribution differs by area due to the different number of neurons present in each area. Significant named dichotomies are marked when the dichotomies are: above $95^{th}$ pctle of null in IP (i.e. significantly above chance during inference), significantly different between IA and IP (RankSum $p < 0.01/35$, Bonferroni corrected for balanced dichotomies). Significant increases from IA to IP were observed in vmPFC for stim pair (purple, $p_{IA} = 0.45, p_{IP} = 0.014$) and preSMA for response (green, $p_{IA} = 0.045, p_{IP} = 0.0010$). Note that stim pair CCGP in AMY was above chance for both IA and IP sessions (purple, $p_{IA} = 0.050, p_{IP} = 0.039$).

**(B)** Same as (A), but for Parallelism Score (PS). Significant increases in PS were present for stim pair in amygdala (purple, $p_{IA} = 1.3x10^{-4}, p_{IP} = 9.0x10^{-8}$) and context in the dorsal anterior cingulate (red, $p_{IA} = 0.99, p_{IP} = 3.8x10^{-1}$ ).

**(C)** Change in decoding accuracy computed as IP – IA for every balanced dichotomy. The gray shaded bar again indicates $5^{th}$-$95^{th}$ pctle of null, which is populated by computing IP – IA for 1000 random pairs of dichotomies in the null distributions of IP and IA computed separately. In general, the null distribution for any "IP-IA" difference plot is computed by drawing such samples from the IA and IP null distributions of the associated area and metric (e.g. the null distribution here is computed using the IP and IA nulls in Fig. 2C, the null distribution for Fig. S3D is computed from Fig. 2D, the null distribution for Fig. S3F is computed from Fig. S3E, etc…).

**(D)** Same as (C), but for changes in CCGP from IA to IP.

**(E)** Parallelism score plot for hippocampus in the IA and IP condition. Coloring, plotting, and significance conventions are identical to those used for Decoding Accuracy and CCGP plots (e.g. Fig. 2C,D). Null distribution here (gray background bars) is computed using the geometric null, the same procedure as CCGP. We note that the $5^{th}$ and $95^{th}$ pctle of null is quite narrow since the rotation procedure used to generate the null distribution produces approximately orthogonal coding vectors in high dimensional spaces. Here, context was significantly elevated in IP compared to IA (red, $p_{IA} = 0.55, p_{IP} = 1.4x10^{-15}$), as was stim pair (purple, $p_{IA} = 0.17, p_{IP} = 1.7x10^{-8}$).

**(F)** Same as (C), but for changes in PS from IA to IP.

**Figure S4. Additional geometric analysis for hippocampus and anterior cingulate cortex during the baseline period.**

**(A)** Decoding accuracy reported for regions other than hippocampus, analogous to Fig. 2E, but for the baseline period instead of the stimulus period. Additionally, trial labels here correspond to the conditions (Response, Context, Outcome) encountered during the previous trial. Details regarding the reported decoding accuracies and null distributions are identical to those described in Fig. 2A. Decoding accuracy for all balanced dichotomies is reported for the inference absent (IA, Left) and inference present (IP, right) conditions, with color-coded lines for named dichotomies connecting IA and IP for each region. Significant increase from IA to IP was observed in dACC for context (red, $p_{IA} = 0.37, p_{IP} = 0.049$).

**(B)** Change in decoding accuracy for all balanced dichotomies in hippocampus associated with the presence of inference (i.e. IP – IA), again computed during the baseline period. Procedure for all reported accuracies and null distribution construction identical to that described in Fig. S3C, except that the analysis used baseline firing rates and condition labels from the previous trial instead of the current trial. Here, context is the only balanced dichotomy whose increase in decodability is significantly above null.

**(C)** Change in cross-condition generalization performance (CCGP) for all balanced dichotomies in hippocampus. See Fig. S3C for plotting details. Context is also the only dichotomy for which the CCGP rises significantly more than the 95$^{th}$ pctle of the geometric null distribution.

**(D)** Parallelism score (PS) for hippocampus in the IA and IP condition. Plot is analogous to Fig. S3E, but computed during the baseline with previous trial labels instead of during the stimulus with current trial labels. Context is the only named dichotomy for which the PS increased to significance above the geometric null in the IP condition. (red, $p_{IA} = 0.37, p_{IP} = 1.2x10^{-10}$)

**(E)** Same as **(B)** and **(C)**, but for parallelism score. Context is the only named dichotomy to increase significantly in IP, and the other un-named dichotomies that also significantly rise are correlated with context.

**(F)** CCGP for balanced dichotomies in the dorsal anterior cingulate cortex (dACC). Associated decoding accuracy for balanced dichotomies in IA and IP shown in (A). Here in the dACC, context (red, $p_{IA} = 0.26, p_{IP} = 0.018$) is also found to be in an abstract format.

**(G)** Parallelism score (PS) for balanced dichotomies in the acc. See Fig. S3E for plotting details. Here, context (red, $p_{IA} = 0.18, p_{IP} = 0.013$) ) emerges as significant in the IP condition.

**(H)** Change in decoding accuracy for balanced dichotomies in acc with inference (IP – IA). The associated plot for IA and IP is in (A). Though context is the dichotomy with the greatest increase in decoding accuracy, it is still below the null 95$^{th}$ pctle in this case.

**(I)** Same as (H), but for CCGP. Here, context is also the dichotomy with the greatest increase, but is still below null 95$^{th}$ pctle.

**(J)** Same as (H), but for PS. Increase in PS for parity is notably significant ($p_{\Delta} = 0.0016$). Context also significantly increases ($p_{\Delta} = 0.026$).

**(K)** Scatter plot of context decoder weights during baseline and stimulus periods. Each point in the scatter plot corresponds to the weights assigned to a single neuron. Weights are averages over 1000 repetitions of decoding with trial re-sampling as described in the methods. Weights are z-scored across neurons for every repetition. Circled neurons are those with weights in the top 25th pctle, separately identified for baseline and stimulus decoding, with neurons in the top 25th pctle for baseline context decoding alone circled red, top 25th pctle for stimulus context decoding alone circled blue, and those in the top 25th pctle of both stimulus and baseline context decoding circled green.

**(L)** Angle distribution for circled neurons shown in (K). Angles are computed as the angle generated by the vector $\langle \beta_{baseline}, \beta_{stimulus} \rangle$ with the vector $\langle 1,0 \rangle$ for each neuron. Thus, angle 0 corresponds to a neuron with $\beta_{stimulus} = 0$ and $\beta_{baseline} > 0$, angle 90 corresponds to a neuron with $\beta_{stimulus} > 0$ and $\beta_{baseline} = 0$, and a neuron with an intermediate angle contributes to both.

**Figure S5. Hippocampal representation geometry unchanged by removal of linearly tuned neurons and neurons recorded inside seizure onset zones.**

Identical analysis to the main geometric analysis shown in Fig. 2, except that hippocampal neurons are excluded from the analysis with the following criteria: in **(A-J)**, neurons with significant linear tuning for Context, Response, or Outcome (2x2x2 ANOVA, Any Main Effect p < 0.01), and in **(K-M)**, neurons with significant linear tuning for Stimulus Identity or Context (4x2 ANOVA, Any Main Effect p < 0.01).

Using the 3-Way ANOVA applied neuron-by-neuron, 455/494 neurons were retained for the stimulus period analysis **(A-C)** and 458/494 neurons were retained for the baseline period analysis **(D-F)**. All primary results for changes in hippocampal geometry were recapitulated apart from decodability of the parity dichotomy during the stimulus period **(A)**.

**(A)** Context decodability (red, $p_{IA} = 0.36, p_{IP} = 0.0001, p_{RS} = 1.6x10^{-31}$). Stim pair decodability (purple, $p_{IA} = 0.078, p_{IP} = 4.2x10^{-5}, p_{RS} = 6.6x10^{-31}$) during the stimulus presentation.

**(B)** Context CCGP (red, $p_{IA} = 0.63, p_{IP} = 0.0016, p_{RS} = 5.2x10^{-34}$). Stim pair CCGP (purple, $p_{IA} = 0.17, p_{IP} = 0.00095, p_{RS} = 5.3x10^{-34}$) during the stimulus presentation.

**(C)** Context PS (red, $p_{IA} = 0.40, p_{IP} = 3.7x10^{-13}$). Stim pair PS (purple, $p_{IA} = 0.83, p_{IP} = 1.2x10^{-7}$) during the stimulus presentation.

**(D)** Context decodability (red, $p_{IA-c} = 0.36, p_{IP-c} = 0.0029, p_{IP-i} = 0.64, p_{RS} = 1.5x10^{-20}$). Stim pair decodability (purple, $p_{IA-c} = 0.071, p_{IP-c} = 0.0021, p_{IP-i} = 0.062, p_{RS} = 2.0x10^{-5}$) during the stimulus presentation.

**(E)** Context PS (red, $p_{IA-c} = 0.40, p_{IP-c} = 4.6x10^{-15}, p_{IP-i} = 0.012$) during the stimulus.

**(F)** Context decodability (red, $p_{IA} = 0.37, p_{IP} = 0.013, p_{RS} = 2.2x10^{-26}$) during the baseline.

**(G)** Context CCGP (red, $p_{IA} = 0.31, p_{IP} = 0.0044, p_{RS} = 1.9x10^{-33}$) during the baseline.

**(H)** Context PS (red, $p_{IA} = 0.12, p_{IP} =0.0055$) during the baseline.

**(I)** Context decodability (red, $p_{IA-c} = 0.55, p_{IP-c} = 0.12, p_{IP-i} = 0.37$) during the baseline.

**(J)** Context PS (red, $p_{IA-c} = 0.66, p_{IP-c} = 8.5x10^{-9}, p_{IP-i} = 0.30$) during the baseline.

Using the 2-Way ANOVA applied neuron-by-neuron, 412/494 neurons were retained for the stimulus period analysis **(A-C)**. The stim-pair dichotomy is no longer decodable after removal of all stimulus-identity tuned neurons, but context is still present in an abstract format.

**(K)** Context decodability (red, $p_{IA} = 0.38, p_{IP} = 0.0088, p_{RS} = 4.1x10^{-2}$ ) during the stimulus.

**(L)** Context CCGP (red, $p_{IA} = 0.51, p_{IP} = 6.0x10^{-4}, p_{RS} = 2.5x10^{-34}$) during the stimulus presentation.

**(M)** Context PS (red, $p_{IA} = 0.77, p_{IP} = 2.3x10^{-6}$) during the stimulus presentation.

**Figure S6. Additional control analyses for Hippocampal representational geometry.**

**(A-F)** Seizure onset zone exclusion analysis. Identical analysis to the main geometric analysis shown in Fig. 2, except that hippocampal neurons neurons recorded in seizure onset zones (SOZs, post-hoc identification) were removed. 410/494 neurons were retained for analysis. The exclusion neurons recorded from SOZ hippocampi led to the full hippocampal geometric analysis being effectively identical to that reported in Fig. 2, with every significant named dichotomy increase during stimulus **(A-C)** and baseline **(D-F)** periods being recapitulated in the absence of SOZ hippocampal neurons.

**(A)** Context decodability (red, $p_{IA} = 0.12, p_{IP} = 0.00044, p_{RS} = 1.0x10^{-26}$). Stim pair decodability (purple, $p_{IA} = 0.034, p_{IP} = 2.0x10^{-7}, p_{RS} = 3.5x10^{-32}$). Parity decodability (purple, $p_{IA} = 0.74, p_{IP} = 0.019, p_{RS} = 2.4x10^{-30}$).

**(B)** Context CCGP (red, $p_{IA} = 0.74, p_{IP} = 0.019, p_{RS} = 1.1x10^{-31}$). Stim pair CCGP (purple, $p_{IA} = 0.084, p_{IP} = 2.7x10^{-5}, p_{RS} = 1.2x10^{-33}$).

**(C)** Context PS (red, $p_{IA} = 3.5x10^{-7}, p_{IP} = 0$). Stim pair PS (purple, $p_{IA} = 0.027, p_{IP} = 1.6x10^{-7}$)

**(D)** Context decodability (red, $p_{IA} = 0.35, p_{IP} = 0.0025, p_{RS} = 1.1x10^{-3}$ ).

**(E)** Context CCGP (red, $p_{IA} = 0.20, p_{IP} = 0.00018, p_{RS} = 2.5x10^{-3}$ ).

**(F)** Context PS (red, $p_{IA} = 0.0022, p_{IP} = 2.0x10^{-5}$).

**(G-M)** Non-inference performance control analysis. Identical analysis to the main geometric analysis shown in Fig. 2, except that IA and IP sessions were distribution-matched for non-inference trial performance. Pairs of sessions from IA and IP with at most 7.5% difference in non-inference trial performance were selected, prioritizing sessions with more hippocampal neurons. This matching process yielded 10 IA sessions (152 neurons) and 10 IP sessions (187 neurons) whose average non-inference performances did not statistically significantly differ (92.8% IA v.s. 94.7% IP, $p_{RS} = 0.58$, RankSum over sessions). All main geometric findings were recapitulated for the stimulus **(G-I)** and baseline **(J-L)** periods. Distribution-matched behavior shown in **(M)** using conventions from Fig. 1, S1.

**(G)** Context decodability (red, $p_{IA} = 0.12, p_{IP} = 0.00051, p_{RS} = 7.6x10^{-7}$). Stim pair decodability (purple, $p_{IA} = 0.014, p_{IP} = 1.2x10^{-5}, p_{RS} = 3.3x10^{-7}$). Parity decodability (purple, $p_{IA} = 0.27, p_{IP} = 0.057, p_{RS} = 5.6x10^{-5}$).

**(H)** Context CCGP (red, $p_{IA} = 0.52, p_{IP} = 0.044, p_{RS} = 6.7x10^{-8}$). Stim pair CCGP (purple, $p_{IA} = 0.17, p_{IP} = 0.0021, p_{RS} = 6.7x10^{-8}$).

**(I)** Context PS (red, $p_{IA} = 0.54, p_{IP} = 0$). Stim pair PS (purple, $p_{IA} = 0.15, p_{IP} = 2.7x10^{-15}$)

**(J)** Context decodability (red, $p_{IA} = 0.32, p_{IP} = 0.0036, p_{RS} = 4.4x10^{-7}$).

**(K)** Context CCGP (red, $p_{IA} = 0.27, p_{IP} = 0.0013, p_{RS} = 6.7x10^{-8}$).

**(L)** Context PS (red, $p_{IA} = 0.015, p_{IP} = 0$).

**Figure S7. Effect of inference and errors on shattering dimensionality as a function of dichotomy difficulty.**

The signature for a high-dimensional representation is a greater degree of non-linear mixing of task variables. "Dichotomy difficulty" is a systematic measure that quantifies the relative amount of non-linear interaction of task variables needed in a population of neurons to make a given dichotomy decodable (see methods for detailed description). **(A)** Example schematics of dichotomies of increasing difficulty. The cubes here represent different unique task conditions realized by three binary variables, and node coloring represents membership of a condition to one of two arbitrary classes assigned for the purposes of dichotomy decoding (identical to Fig. S2). Note: the difficulty 4 dichotomy corresponds to context and difficulty 12 dichotomy corresponds to parity (Fig. S2). **(B-G)** Decoding accuracy as a function of dichotomy difficulty for different regions. Reported values (mean +/- SEM) are computed over dichotomy decoding accuracies, where the average decoding accuracy for each dichotomy is computed with 1000 repetitions of re-sampled estimation (see methods). The blue, red, and green curves correspond to correct IA trials, correct IP trials, and incorrect IP trials respectively. Black dashed lines indicate chance level (50% for binary decoding), horizontal black lines indicate the 5$^{th}$ and 95$^{th}$ pctle of the null distribution. P-values are computed by conducting a one-way ANOVA over dichotomies independently for every dichotomy difficulty (Bonferroni MCC). This value is not meaningfully computable for difficulty 12, which contains a single dichotomy (the parity dichotomy), and is therefore not reported. Hippocampus alone **(B)** exhibits an increase in decoding accuracy from IA to IP sessions, with more difficulty dichotomies rising above 95$^{th}$ pctle null in IP. A collapse in representational dimensionality on incorrect trials (purple curves) is present in the hippocampus **(B)**, and is also present in other areas, most prominently in the ventral temporal cortex **(D)** and the amygdala **(E)**.

**Figure S8. Balanced dichotomy analysis of neural geometry in ventral temporal cortex.**

Ventral temporal cortex (VTC) strongly encodes high-level features of visual stimuli, necessitating the introduction of two new dichotomies that capture stimulus identity, while not directly corresponding to any of the principal manipulated variables in the task (Response, Context, Outcome). The AB vs CD and AD vs BC dichotomies are "stimulus" dichotomies in that they represent systematic differences in coding between unrelated stimuli arbitrarily paired together, unlike the AC vs BD dichotomy which pairs stimulus identities for which responses are identical across the two contexts. That is, A/C response is L in Context 1 and R in Context 2, and v.v. B/D response is R in Context 1 and L in Context 2. These are the images whose correct responses "switch together" across contexts. Note that the new stimulus dichotomies are correlated with other named dichotomies: AB vs CD is correlated with outcome and AD vs BC is correlated with the parity dichotomy. Thus, high AB vs CD decodability will lead to increased outcome decodability. However, CCGP is robust to these dichotomy correlations, and will be low for correlated dichotomies even if decodability is increased. These three dichotomies (AB vs CD, AC vs BD, and AD vs BC) are particularly relevant to vtc given its strong stimulus representations.

**(A)** Dichotomy decodability during pre-stimulus baseline. None of the balanced dichotomies are decodable during IA or IP ($p > 0.05$ for all dichotomies).

**(B)** Dichotomy decodability during the stimulus presentation period. All three named stimulus dichotomies are highly decodable both during IA and IP. Correlated dichotomies also demonstrated above-chance decodability. Dichotomies: purple, $p_{IA} = 6.8x10^{-13}, p_{IP} = 6.6x10^{-1}$ , brown, $p_{IA} = 2.2x10^{-9}, p_{IP} = 6.0x10^{-14}$, pink, $p_{IA} = 1.1x10^{-13}, p_{IP} = 6.7x10^{-14}$. Notably, context is not significantly decodable in either IA or IP (red, $p_{IA} = 0.24, p_{IP} = 0.38$).

**(C)** Dichotomy CCGP for VTC during the stimulus presentation period. Two stimulus dichotomies are in an abstract format in IA and all three are in an abstract format in IP (purple, $p_{IA} = 0.0054, p_{IP} = 0.0036$, brown, $p_{IA} = 0.057, p_{IP} = 0.0029$ , pink, $p_{IA} = 0.0030, p_{IP} = 0.0032$). All remaining dichotomy CCGP values are at chance apart from response and context, which are significantly below chance in IP (green, $p_{IA} = 0.93, p_{IP} = 0.96$, red, $p_{IA} = 0.84, p_{IP} = 0.94$).

**(D)** Dichotomy PS for VTC during the stimulus presentation period. Again, two stimulus dichotomies are in an abstract format in IA, and all three are in an abstract format in IP (purple, $p_{IA} = 0, p_{IP} = 4.3x10^{-13}$, brown, $p_{IA} = 0.73, p_{IP} = 0$, pink, $p_{IA} = 0, p_{IP} = 5.9x10^{-7}$).

**(E)** Dichotomy decodability analysis for incorrect trials during the stimulus presentation period. Decoders are trained on correct trials and evaluated on incorrect trials (balanced by condition) in IP sessions. Plotting conventions are identical to those described in Fig. 3,S6. Note that all three stimulus identity-related dichotomies are still highly significantly decodable during incorrect trials in IP sessions (purple, $p_{IP-i} = 7.8x10^{-11}$, brown, $p_{IP-i} = 1.1x10^{-13}$, pink, $p_{IP-i} = 8.7x10^{-11}$).

**Figure S9. Cross-condition generalization performance for stimulus identity and context defined over stimulus pairs.**

To fully disentangle and study the interaction between stimulus coding and context, geometric analysis of balanced dichotomies must be replaced by new analyses that are defined for pairs of individual stimuli, thus allowing for the study of stimulus coding un-ambiguously without arbitrarily grouping together stimuli as is necessary in the balanced dichotomy approach. When considering a pair of stimuli (e.g. A and B) across two contexts (e.g. 1 and 2), there are four possible task conditions (A1, B1, A2, B2). On these points, stimulus (A1A2 vs B1B2) and context (A1B1 vs A2B2) can be decoded in a straightforward manner, but is not informative about the format in which stimulus and context are encoded. The CCGP for stimulus across contexts **(A-C)** and for context across stimuli **(D-F)** provide information about the structure of the two variables and how they interact.

Consider within-context training/testing. The procedure is summarized in **(A)**, which shows a linear decoder (green bar) trained between stimuli A and B in context 1 (green + and – correspond to class labels for training). The decoder is then generalized to context 2, where stimulus identity is decoded (purple bar, + and – for class labels). This procedure is broken down step-by-step for training in **(B)** and testing in **(C)**. In addition, arrows showing persistent stimulus and context coding vectors (black/dashed arrows) have been drawn alongside the vector orthogonal to the hyperplane learned during train/test (colored arrow passing through the bar). Note that, for this formulation of Stimulus CCGP, the stimulus coding vector and the normal vector to the hyperplane are parallel in **(B)** and **(C)**. Thus, in cases with high within-context train/test Stimulus CCGP, stimulus information is present in an abstract format across contexts.

The same procedures for computing CCGP can be applied for studying the format of context organizing across pairs of stimuli **(D-F)**, with schematic details identical to those described above for Stimulus CCGP. Here, high Context CCGP indicates that context is encoded abstractly across the different stimuli.

**Figure S10. Additional geometric stimulus-pair analysis for hippocampus and ventral temporal cortex.**

**(A)** Average distances between stimulus representations in hippocampus (HPC) in IA and IP. All plotted points correspond to named, interpretable groups of conditions defined by pairs of stimuli presented in both contexts. For example, the green dot in IA indicates the average distance (~5.1Hz) between the condition centroids for stimulus A and stimulus D (averaged over contexts). Distance is computed as Euclidean distance between the stimulus centroids, each of which is an N (# of neurons) dimensional vector of average firing rates during stimulus presentation. Neuron counts are balanced between IA and IP to allow for direct distance comparisons. Null distributions here are geometric nulls, and are identical to those used for CCGP and PS. Significance of the difference between IA and IP inter-stimulus distances is established by RankSum test computed over stimulus pairs, and n.s. indicates $p > 0.05$.

**(B)** Same as **(A)**, but for ventral temporal cortex (VTC).

**(C)** Decodability of stimuli also did not significantly change between IA and IP for HPC. Here, decoding accuracies are reported for each unique pair of stimuli with 1000 repetitions of trial sub-sampling. Null distributions are constructed with trial-label shuffling, and the gray bars correspond to the boundary of the $5^{th}$ to $95^{th}$ pctle of the null. Significance of the difference between IA and IP decodability is also established by Ranksum test over average decoding accuracies and n.s. indicates $p > 0.05$.

**(D)** Same as **(C)**, but for VTC.

**Figure S11. Additional context CCGP analysis over stimulus pairs for hippocampus and ventral temporal cortex (stimulus period).**

Change in context decoding accuracy from IA to IP evaluated over individual stimulus pairs is shown for the hippocampus **(A)** and ventral temporal cortex **(B)**. Individual points correspond to context decoding accuracy averaged over 1000 repetitions of decoding/CCGP/Parallelism Score estimation with trial re-sampling. All plotting conventions for geometric plots are identical to those used in Fig. 4,S10.

**(C)** and **(D)** show Context CCGP and Context Parallelism Score over stimuli for VTC. Analogous plots for HPC are Fig. 4G,H.

Two exemplar neurons are shown, one from HPC **(E,G)** and one from VTC **(F,H)** that feature both stimulus tuning and context modulation. Plotting conventions are identical to previous raster/psth plots apart from the colors and conditions plotted, which here are two stimuli (A and B) in the two contexts. Responses to task conditions for the two neurons are summarized in **(G,H)**, which show mean ± s.e.m. firing rates by condition for spikes counted on individual trials during the stimulus period (0.2s to 1.2s after stimulus onset). The same trials used to compute **(G)** and **(H)** are shown in **(E)** and **(F)** respectively, and condition colors are matched between the two sets of plots. Black arrows indicate the direction in which the firing rate for a stimulus is modulated by a shift in context. The HPC neuron **(G)** shows consistent modulation by context since both arrows point downward, whereas the VTC neuron **(H)** shows inconsistent modulation by context since one arrow points downwards and the other points upward.

**(I)** Change in the consistency of context-modulation for stimuli averaged over all neurons in VTC and HPC. Context modulation consistency is the tendency for a neuron's firing rate to shift consistently (increase or decrease) to encode context across stimuli. This consistency can take on values between 0 (increase in firing rate to encode context for half of the stimuli, decrease in firing rate for the other half) and 4 (either increase or decrease in firing rate for all four stimuli). An interaction effect is observed between context modulation consistency for HPC neurons and VTC neurons in IA and IP in the absence of main effects (2 x 2 ANOVA, $p_{Area} = 0.36$, $p_{IA/IP} = 0.64$, $p_x = 4.5x10^{-5}$), revealing significant increases in context modulation consistency in HPC from IA to IP with concurrent decreases in VTC.

**Figure S12. Hippocampal MDS plots showing increases in context and stimulus parallelism for all stimulus pairs.**

MDS plots analogous to that shown in Fig. 4K, but plotted in 2D for individual stimulus pairs. Colored points represent the mean condition response of all HPC neurons during IA or IP to a given stimulus in a given context. Stimuli are color coded according to identity (e.g. in A, red points are condition responses to stimulus A and blue points are condition responses to stimulus B), and are connected by a line of the same color to reflect the context coding direction for that stimulus. Stimuli in the same context are connected by shaded lines that are blue for context 1 and red for context 2. Since MDS is conducted independently for IA and IP, individual MDS axes are not directly comparable between IA and IP, but the relative distances are comparable since the number of neurons is matched between IA and IP, and both are dimensionally reduced to the same number of MDS dimensions ($N_{dim} = 2$). Condition averages are computed using only correct trials. Evidence of disentangling of context and stimulus identity is evident across most stimulus pairs, with the notable exception of the B/D stimulus pair **(E)**, which is perfectly correlated with outcome and therefore cannot be dissociated from outcome using CCGP. The emergence of quadrilaterals with approximately parallel sides for all other stimulus pairs **(A-D, F)** is a signature of disentangling of stimulus identity and context, and clearly demonstrates the abstract format of both variables.

**Figure S13. Implementation of geometric changes in hippocampal representation – Stimulus period.**

**(A)** Distances between centroids for balanced dichotomies shown for all regions other than HPC. Plotting conventions are identical to those used in Fig. 5E. Note: neuron counts were only balanced across IA/IP within-region, so distances in different regions are computed in spaces with different dimensionality and are therefore not meaningfully comparable. Significant change in average dichotomy separation determined through Bonferroni MCC RankSum where * indicates $p < 0.05/35$, and n.s. otherwise.

**(B)** Changes in inter-centroid distance for balanced dichotomies. Points in these plots are the differences between IP and IA distances shown in **(A)**. Null distributions are computed No distances for named dichotomies increased or decreased more than would be expected by chance (outside $5^{th}$-$95^{th}$ pctle null).

**(C)** Firing rates for individual task conditions (8 total) for all regions other than HPC. Plotting conventions identical to those used in Fig. 5I. Task conditions are color coded based on the identity of the presented stimulus (same as Fig. 1B. 5I,J). Significant change in average dichotomy separation determined through Bonferroni MCC RankSum where * indicates $p < 0.05$, and n.s. otherwise.

**(D)** Changes in hippocampal firing rates for 3 different sub-groups of session pairs: instruction-successful sessions (IS, n = 5 patients, 10 sessions), instruction-unsuccessful sessions (IU, n = 4 patients, 8 sessions), and instances of first-session inference (S1, n = 3 patients, 6 sessions). These session groups were constructed to compare the effect of inference acquisition with the passage of time (see Methods for details). Firing rate changes here are computed during the stimulus presentation period (0.2s to 1.2s after stim onset) from consecutive pre-instruction and post-instruction sessions. Points are average changes in condition-averaged firing rates (8 unique conditions). Changes in firing rate that significantly differed from zero (t-test, $p < 0.05/3$) are indicated with a "*". IS group alone exhibited significant decrease in firing rate. IU group exhibited an increase in firing rate.

Changes in hippocampal single-neuron tuning quantified by 3-way ANOVA (Response, Context, Outcome) with interactions. Significant factors ($p < 0.05$) were identified for every neuron and averages of both the number of factors per neuron **(E)** and the depth of tuning of those factors quantified through $-\log_{10}(p_{ANOVA})$ **(H)** reported (mean ± s.e.m. across neurons) for the IA (red) and IP (blue) groups. Significance of difference between IA and IP for both the number of factors ($\mathbf{E}, p_{RS} = 0.041$) and the tuning strength ($\mathbf{H}, p_{RS} = 0.027$) was assessed by RankSum test over neurons between the two groups, and "*" indicates $p_{RS} < 0.05$.

**(F)** same as **(E),** but for all regions other than HPC.

**(I)** same as **(H),** but for all regions other than HPC.

**(G)** The change in the distribution of trials projected along the coding direction for context was visualized during IA (above) and IP (below). The red and blue histograms are the distribution of projected trials from context 1 and 2 respectively, with the red and blue vertical lines indicating the mean of each distribution. Positive and negative values for projection were arbitrarily established by computing the coding vector as (ctxt1 – ctxt2).

**(J)** Plot showing the fraction of hippocampal neurons that exhibit task selectivity for IA (red) and IP (blue) sessions. Selectivity is determined independently for every neuron using a 4x2 ANOVA (Stimulus Identity, Context), with a per-factor significance threshold of $p < 0.05$. Significant differences in tuned fractions between IA and IP assessed with z-test. **(J)** Plot showing the fraction of hippocampal neurons that exhibit task selectivity for IA (red) and IP (blue) sessions. Selectivity is determined independently for every neuron using a 4x2 ANOVA (Stimulus Identity, Context), with a per-factor significance threshold of $p < 0.05$. Significant differences in tuned fractions between IA and IP assessed with z-test.

**(K)** same as **(J)**, but for VTC.

**Figure S14. Implementation of geometric changes in hippocampal representation – Baseline period.**

**(A)** Average variance of individual trials projected onto the coding direction for every dichotomy. Plotting conventions identical to those in Fig. 5I. Average variance along coding directions decreased significantly between IA and IP sessions ($p_{RS} = 6.5x10^{-13}$, RankSum over dichotomies).

**(B)** Changes in variance between IP and IA for all dichotomies shown in **(A)**. No named dichotomies fall outside the $5^{th}$-$95^{th}$ pctle of the null distribution.

**(C)** Population distance between dichotomy centroids for dACC at baseline. All plotting conventions identical to those used in Fig. 5G. Average distance between dichotomy centroids increased when comparing IA to IP sessions ($p_{RS} = 2.9x10^{-8}$, RankSum over dichotomies). Notably, context centroids emerged as significantly separated in IP than expected by chance ($p_{IA} = 0.48$, $p_{IP} = 0.0065$)

**(D)** Changes in distance between IP and IA sessions for all dichotomies shown in **(C)**. Context alone (red, $p_{\Delta} = 0.047$) exhibited a greater increase in distance than expected by chance.

**(E)** Same as **(A)**, but for projective variance in dACC during the baseline. Average variance along coding directions increased significantly between IA and IP sessions ($p_{RS} = 6.0x10^{-3}$, RankSum over dichotomies).

**(F)** Same as **(B)**, but for differences in dACC variance during the baseline computed using **(E)**.

**(G)** Baseline firing rate averaged by condition (8 total) for the hippocampus. Plotting conventions are identical to those in Fig. 5I, S13C. Reduction from IA to IP is significant ($p < 0.05$, RankSum over conditions).

**(H)** Baseline firing rates averaged by condition for all regions other than hippocampus. Significance of change in firing rate also assessed by RankSum over conditions ("*" indicates $p < 0.05$, n.s. otherwise). Note: most regions (apart from AMY) exhibit slight, but significant increases in baseline firing rate during in IP compared with IA.

**Figure S15. Additional analysis of the effect of instructions on behavior.**

Task performance shown as a moving average over blocks for the instruction-successful (IS) subject group **(A,B)**, the instruction unsuccessful (IU) subject group **(C,D)**, and the session one inference (S1) subject group **(E,F)**. Plots are similar to those shown in Fig. 1F,G, except Pre and Post plots show performance over time for sessions recorded immediately preceding and immediately following verbal instructions describing latent task structure. For example, for the IS group, pre-instruction performance is plotted in **(A)** for non-inference trials (black) and inference trials (gray). Average performance is computed as a moving average with a 3-block window on the last three trials before a context switch (non-inference) and on the first inference trial after a switch (inference). Error bars are standard errors computed over subjects. Chance performance is indicated with the dashed line at $y = 0.5$.

**Figure S16. Additional analysis of the effect of instructions on hippocampal neural geometry.**

Geometric metrics shown are computed over balanced dichotomies, and are plotted using the same conventions as discussed previously (see Fig. 2, methods for details), except for the left and right columns of each plot correspond to pre-instruction and post-instruction geometry respectively. Plots in the figure are segregated according to subject group (instructions successful - IS, instructions unsuccessful IU, session one inference - S1) and analysis period (Stimulus, Baseline). Only context is plotted as a named dichotomy for visual clarity.

**(A)** CCGP (context, red, $p_{pre} = 0.27, p_{post} = 0.046, p_{RS} = 1.4x10^{-31}$) and **(B)** PS (context, red, $p_{pre-c} = 0.029,\ p_{post-c} = 3.5x10^{-6}, p_{post-}\ = 0.0028$) for the IS group during the stimulus period.

**(C)** Decoding accuracy (context, red, $p_{pre-c} = 0.35, p_{post-}\ = 0.0014, p_{post-i} = 0.55, p_{RS} = 1.4x10^{-20}$), **(D)** CCGP (context, red, $p_{pre} = 0.33, p_{post} = 0.0037, p_{RS} = 3.0x10^{-34}$), and **(E)** PS (context, red, $p_{pre-c} = 0.017, p_{post-c} = 7.5x10^{-8}, p_{post-}\ = 0.40$) for the IS group during the baseline period.

**(F)** CCGP (context, red, $p_{pre} = 0.56, p_{post} = 0.39, p_{RS} = 0.004$) and **(G)** PS (context, red, $p_{pre} = 0.81, p_{post} = 0.95$) for the IU group during the stimulus period.

**(H)** Decoding accuracy (context, red, $p_{pre} = 0.45, p_{post} = 0.45, p_{RS} = 0.68$), **(I)** CCGP (context, red, $p_{pre} = 0.45, p_{post} = 0.47, p_{RS} = 0.15$), and **(J)** PS (context, red, $p_{pre} = 0.93, p_{post} = 0.30$) for the IU group during the baseline period.

**(K)** CCGP (context, red, $p_{pre} = 0.23, p_{post} = 0.19, p_{RS} = 0.0045$) and **(L)** PS (context, red, $p_{pre} = 6.3x10^{-8}, p_{post} = 4.5x10^{-7}$,) for the S1 group during the stimulus period.

**(M)** Decoding accuracy (context, red, $p_{pre} = 0.37, p_{post} = 0.47, p_{RS} = 0.036$), **(N)** CCGP (context, red, $p_{pre} = 0.30, p_{post} = 0.50, p_{RS} = 5.9x10^{-7}$), and **(O)** PS (context, red, $p_{pre} = 1.7x10^{-5}, p_{post} = 0.029$) for the S1 group during the baseline period.

**Table S1. Tabulation of Patients, Sessions, Behavior, and Neurons.**

Summary of patient information, the number of sessions performed, the behavioral classification at the patient and session level, and the number of recorded neurons per region per session. Patient behavior is defined with respect to the first instance of high-level verbal instructions (see Fig. 6), where: IS – "instructions successful", IU – "instructions unsuccessful", S1 – "first session inference", and N/A – "did not qualify for analysis". Session behavior is defined with respect to performance on the first available inference trial, where: IA – "inference absent", IP – "inference present", X – "at or below chance non-inference performance".

**Table S2. Definition of all balanced dichotomies.**

Class assignment and name of all 35 balanced dichotomies used in geometric balanced dichotomy analysis. Dichotomies where the name is "N/A" do not have a clear interpretation with respect to task construction. Identity of the task conditions participating in the balanced dichotomies is shown to the right.

**Table S3. Definition of all stimulus dichotomies.**

Task condition assignment for stimulus dichotomies. These dichotomies are used in Fig. 4B-E and associated supplements whenever there is a reference to "Stimulus CCGP" or "Stimulus Parallelism Score". Partial and full correlations with other task variables are noted for each stimulus dichotomy.

**Table S4. Definition of all context dichotomies.**

Task condition assignment for context dichotomies. These dichotomies are used in Fig. 4G,H and associated supplements whenever there is a reference to “Context CCGP” or “Context Parallelism Score”. Partial and full correlations with other task variables are noted for each context dichotomy.

**Supplementary Video 1. Transformation of hippocampal geometry shown with MDS of real data.**

Visualization of the transformation of the representational geometry in the hippocampus shown using MDS of condition-averaged responses for all recorded hippocampal neurons during the stimulus period. Plotting conventions and data are identical to those used in Fig. 4I. Here, we use linear interpolation between the starting and ending geometry shown in the video, which correspond to inference absent and inference present sessions respectively. The video is meant to provide intuition for how the task conditions are represented differently in neural state space in the presence and absence of inference behavior.

### Figure S1

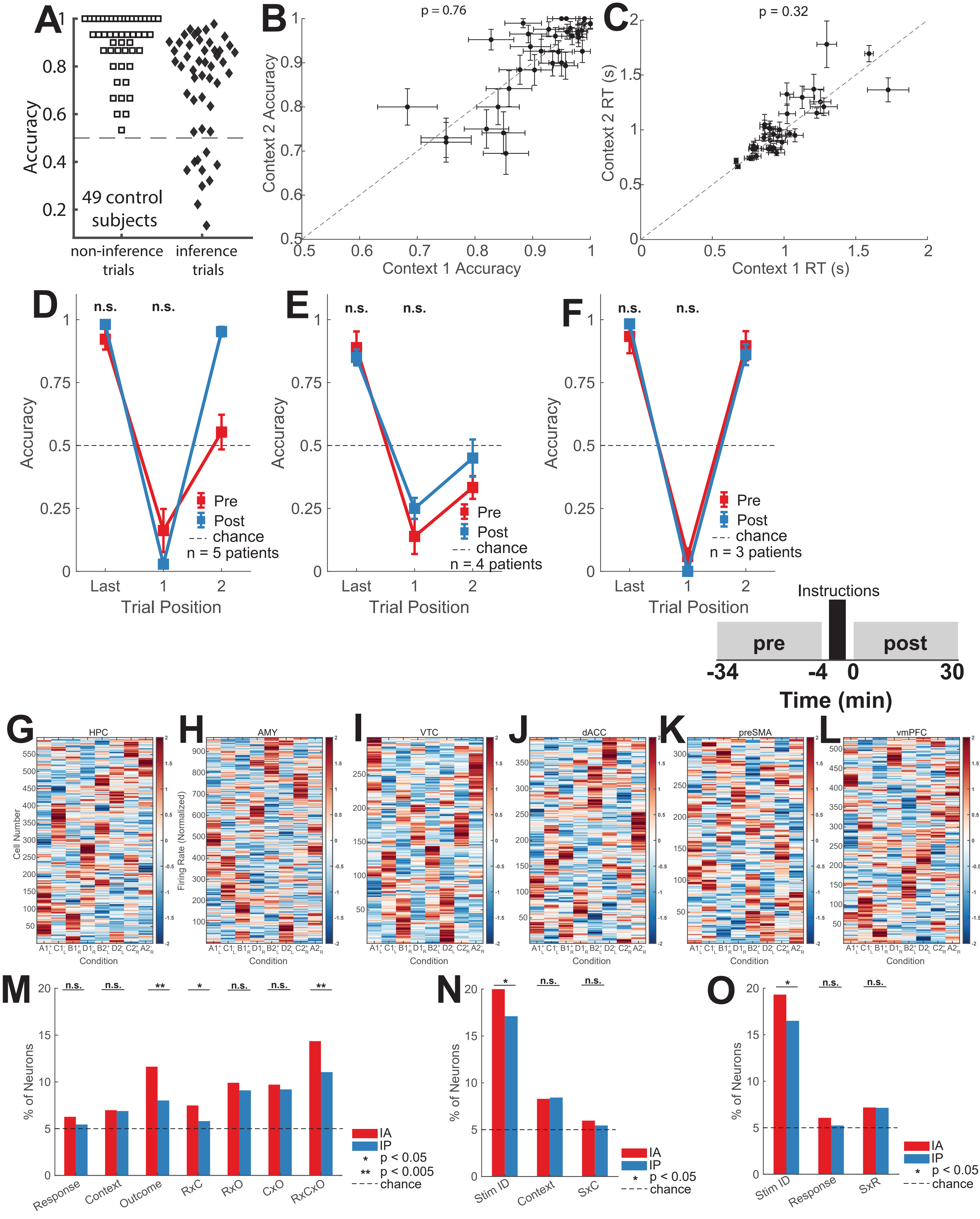

A
Accuracy
49 control subjects
non-inference trials
inference trials
B
p = 0.76
Context 2 Accuracy
Context 1 Accuracy
C
p = 0.32
Context 2 RT (s)
Context 1 RT (s)
D
n.s.
n.s.
Accuracy
Pre
Post
chance
n = 5 patients
Last
Trial Position
E
n.s.
n.s.
Accuracy
Pre
Post
chance
n = 4 patients
Trial Position
F
n.s.
n.s.
Accuracy
Pre
Post
chance
n = 3 patients
Trial Position
Instructions
pre
post
-34
-4
0
30
Time (min)
G
HPC
Cell Number
Condition
H
AMY
Firing Rate (Normalized)
Condition
I
VTC
Condition
J
dACC
Condition
K
preSMA
Condition
L
vmPFC
Condition
M
% of Neurons
Response
Context
Outcome
RxC
RxO
CxO
RxCxO
n.s.
**
*
IA
IP
* p < 0.05
** p < 0.005
chance
N
% of Neurons
Stim ID
Context
SxC
*
n.s.
IA
IP
* p < 0.05
chance
O
% of Neurons
Stim ID
Response
SxR
*
n.s.
IA
IP
* p < 0.05
chance

### Figure S2

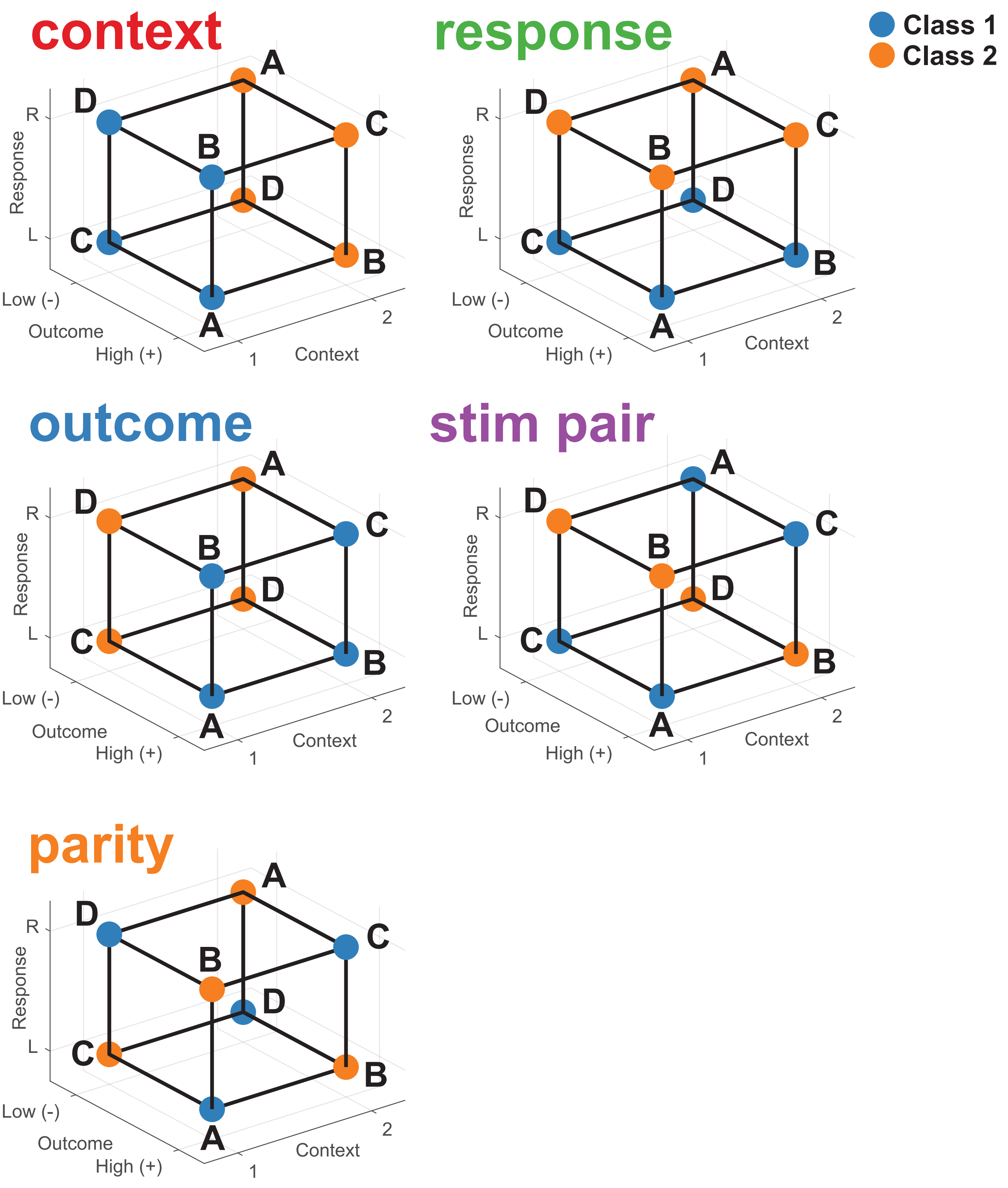

context
response
outcome
stim pair
parity
Class 1
Class 2
A
B
C
D
R
L
Response
Low (-)
Outcome
High (+)
1
2
Context

### Figure S3

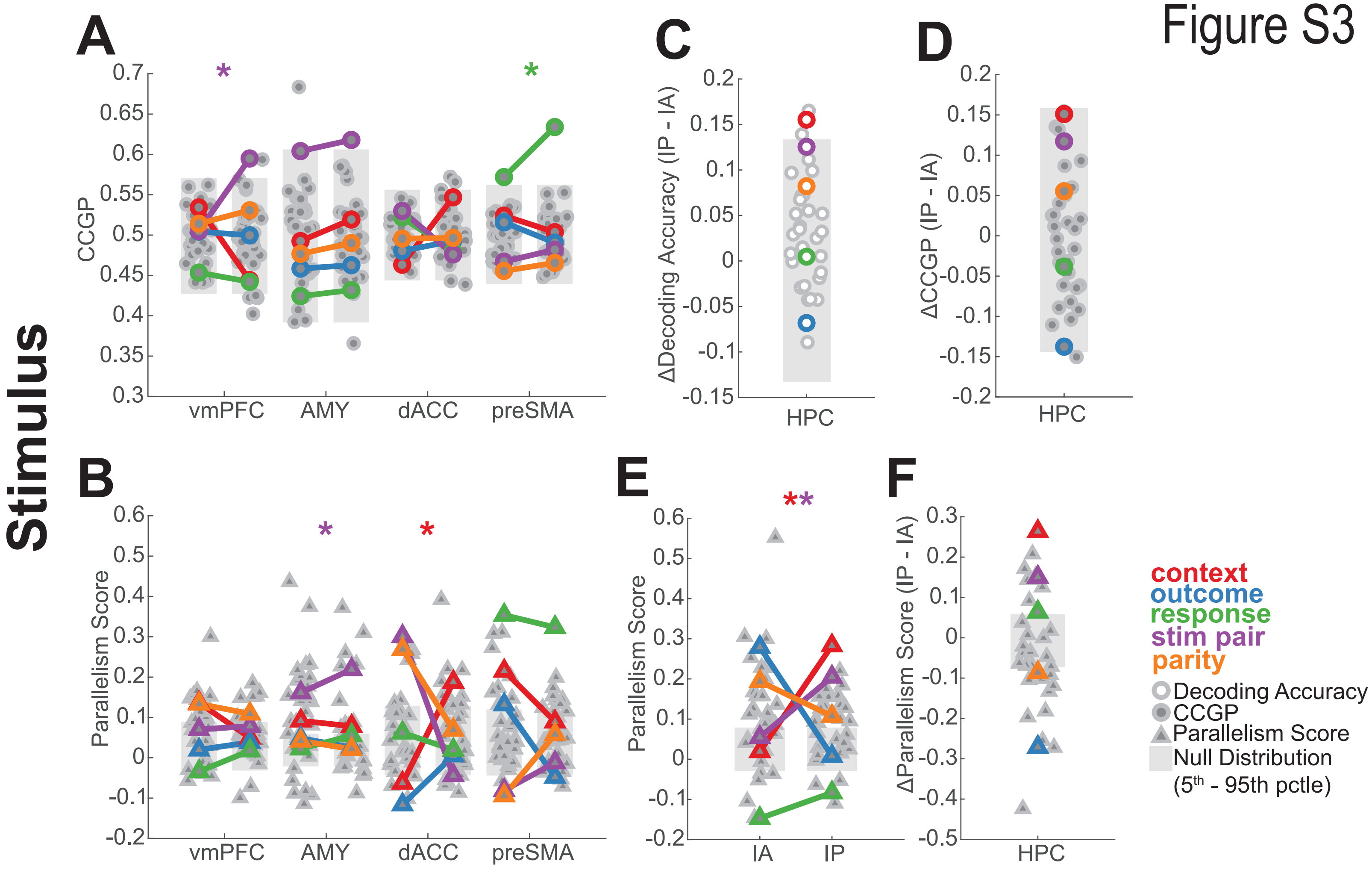

Stimulus
A
CCGP
vmPFC
AMY
dACC
preSMA
C
ΔDecoding Accuracy (IP - IA)
HPC
D
ΔCCGP (IP - IA)
HPC
B
Parallelism Score
vmPFC
AMY
dACC
preSMA
E
Parallelism Score
IA
IP
F
ΔParallelism Score (IP - IA)
HPC
context
outcome
response
stim pair
parity
Decoding Accuracy
CCGP
Parallelism Score
Null Distribution
(5th - 95th pctle)

### Figure S4

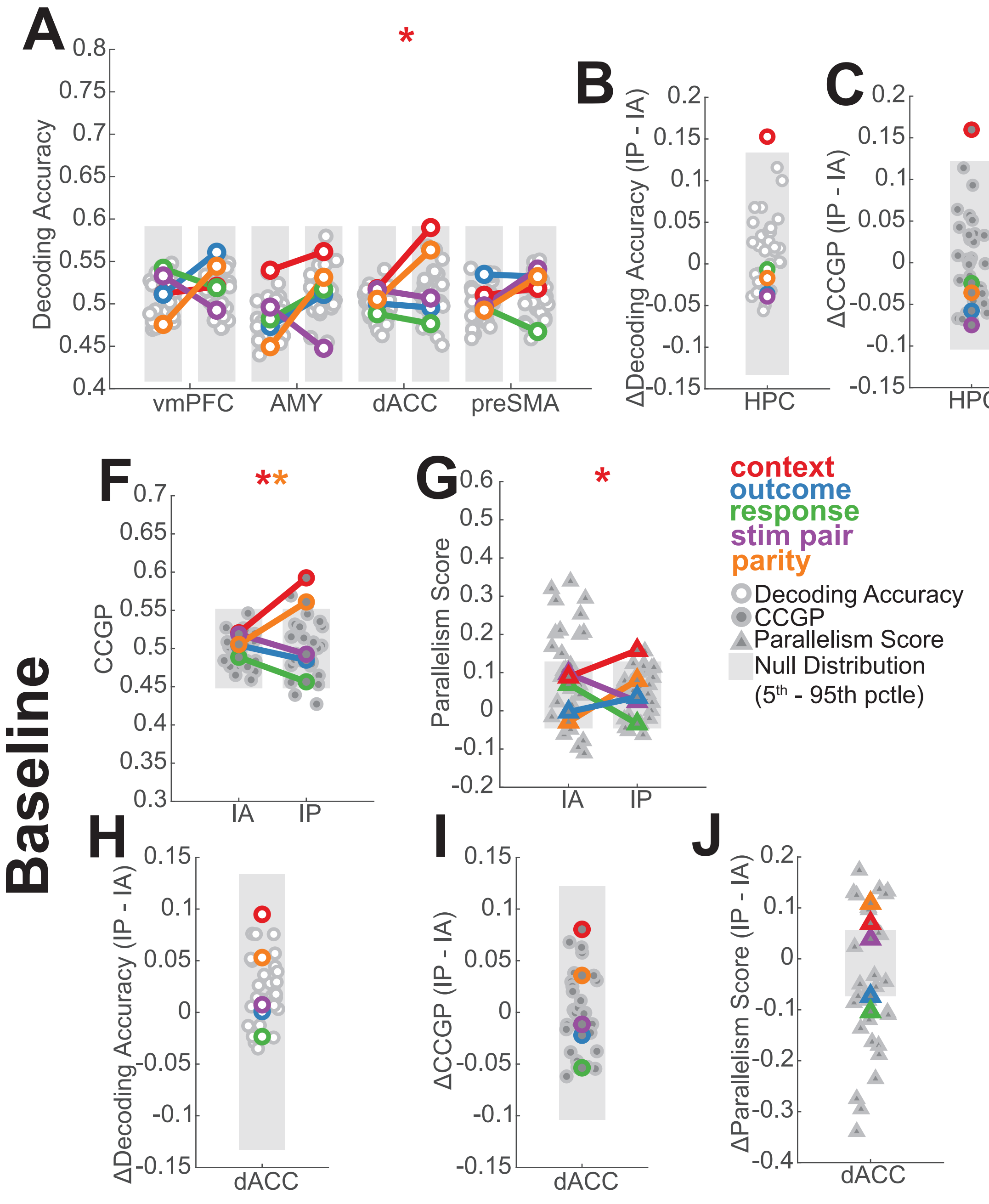

A
B
C
D
E
F
G
H
I
J
Baseline
Decoding Accuracy
vmPFC
AMY
dACC
preSMA
ΔDecoding Accuracy (IP - IA)
ΔCCGP (IP - IA)
Parallelism Score
ΔParallelism Score (IP - IA)
HPC
IA
IP
CCGP
context
outcome
response
stim pair
parity
Decoding Accuracy
CCGP
Parallelism Score
Null Distribution
(5th - 95th pctle)

Figure S5

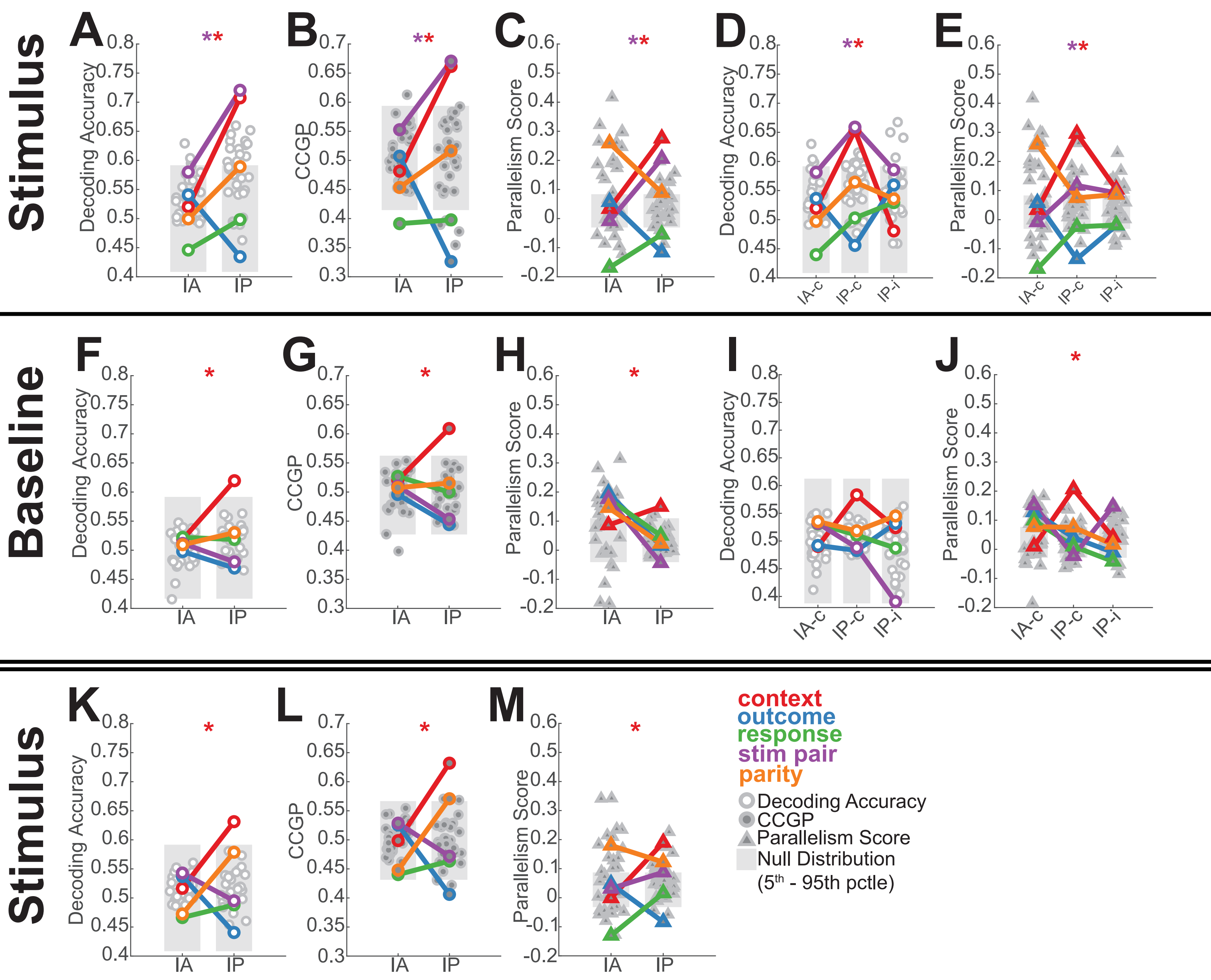
Stimulus
Baseline
Stimulus
A
B
C
D
E
F
G
H
I
J
K
L
M
Decoding Accuracy
CCGP
Parallelism Score
IA
IP
IA-c
IP-c
IP-i
**
*
context
outcome
response
stim pair
parity
Decoding Accuracy
CCGP
Parallelism Score
Null Distribution
(5th - 95th pctle)

### Figure S6

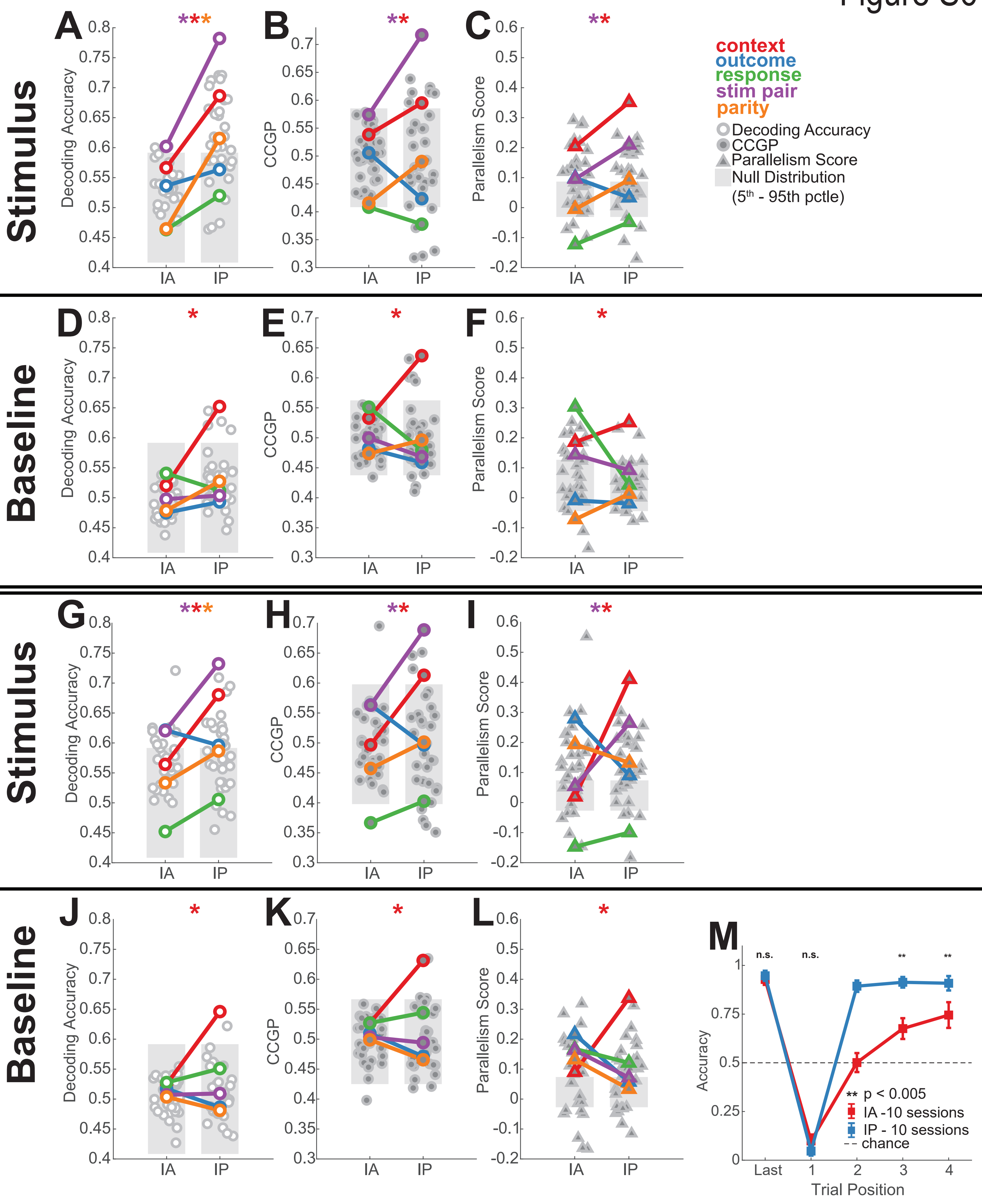

Stimulus
Baseline
Stimulus
Baseline
A
B
C
D
E
F
G
H
I
J
K
L
M
Decoding Accuracy
CCGP
Parallelism Score
IA
IP
context
outcome
response
stim pair
parity
Decoding Accuracy
CCGP
Parallelism Score
Null Distribution
(5th - 95th pctle)
n.s.
n.s.
**
**
Accuracy
Trial Position
Last
** p < 0.005
IA -10 sessions
IP - 10 sessions
chance

Figure S7

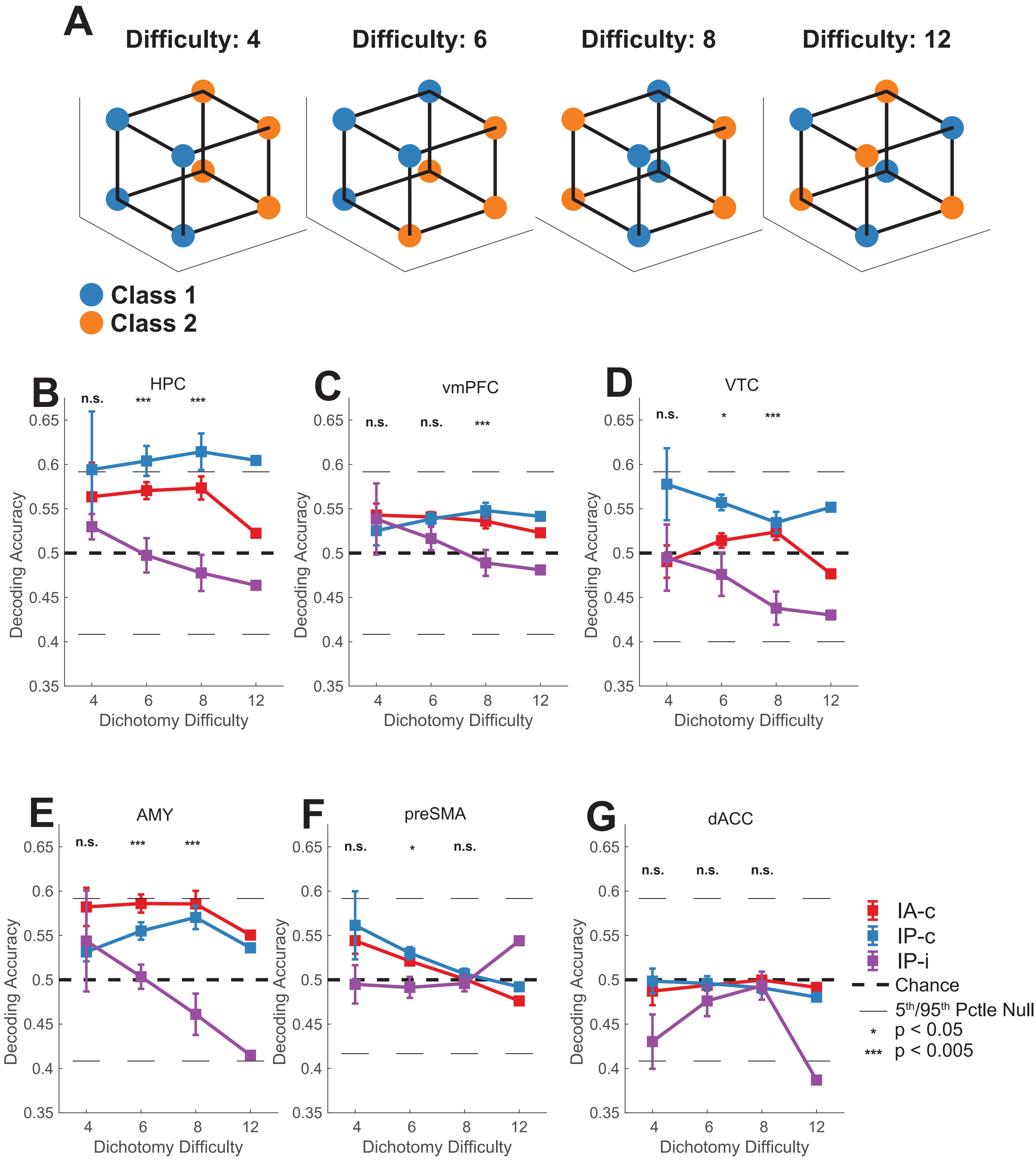
A
Difficulty: 4
Difficulty: 6
Difficulty: 8
Difficulty: 12
Class 1
Class 2
B
HPC
C
vmPFC
D
VTC
E
AMY
F
preSMA
G
dACC
n.s.
*
***
Decoding Accuracy
Dichotomy Difficulty
0.35
0.4
0.45
0.5
0.55
0.6
0.65
4
6
8
12
IA-c
IP-c
IP-i
Chance
5th/95th Pctle Null
* p < 0.05
*** p < 0.005

Figure S8

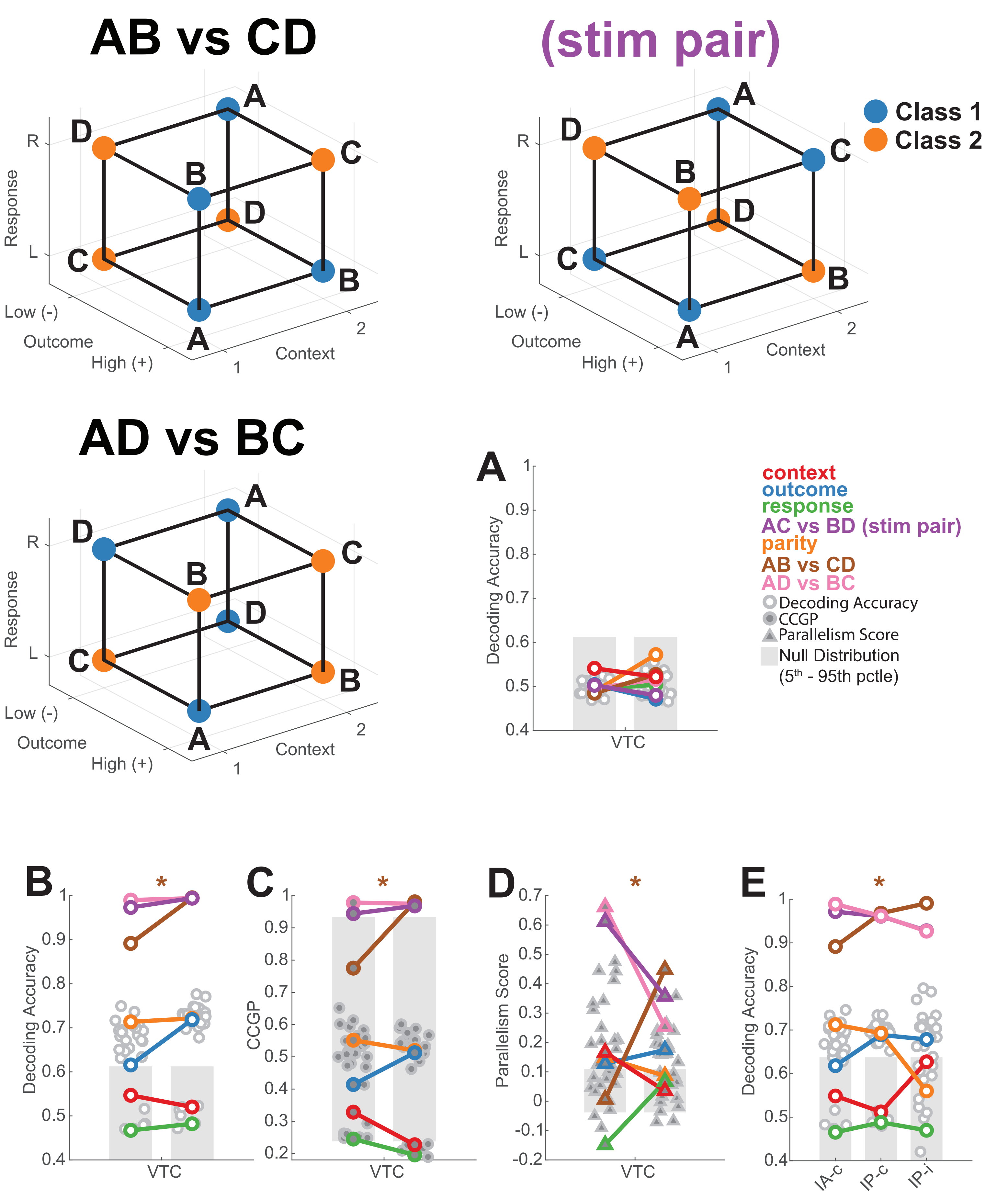

AB vs CD
AC vs BD
(stim pair)
AD vs BC
Class 1
Class 2
A
B
C
D
Response
R
L
Low (-)
Outcome
High (+)
Context
1
2
A
B
C
D
E
Decoding Accuracy
CCGP
Parallelism Score
VTC
IA-c
IP-c
IP-i
context
outcome
response
AC vs BD (stim pair)
parity
AB vs CD
AD vs BC
Decoding Accuracy
CCGP
Parallelism Score
Null Distribution
(5th - 95th pctle)
*

Figure S9

### Stimulus Decoding/CCGP (train/test within context)

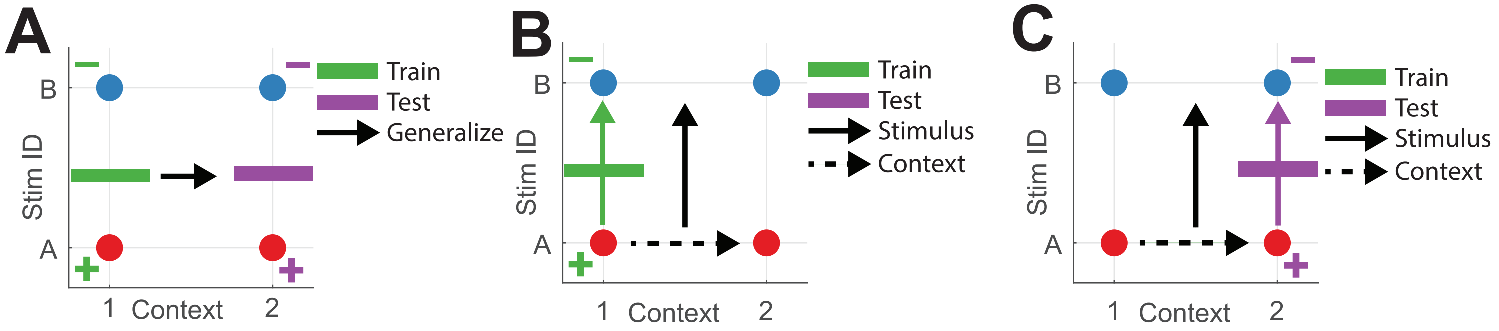


### Context Decoding/CCGP (train/test within stimulus)

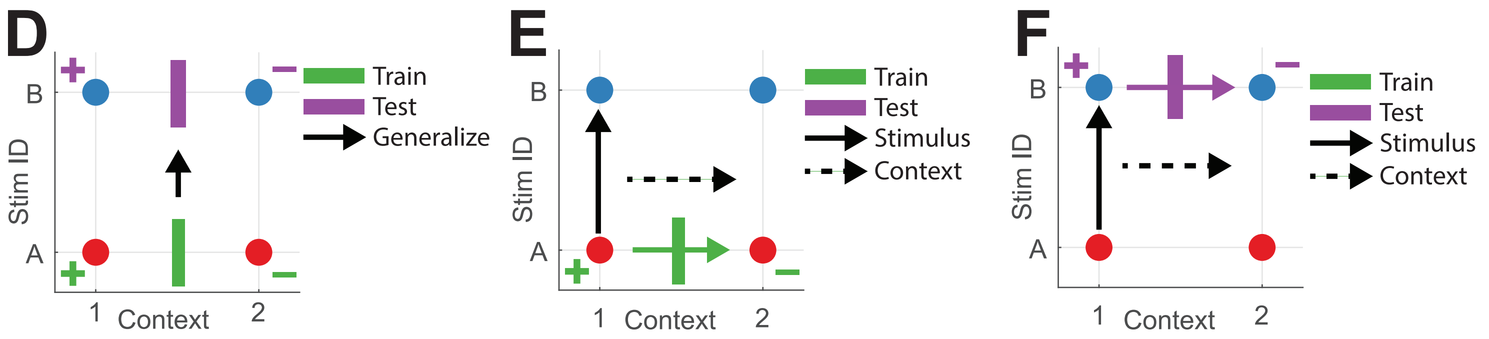

### Figure S10

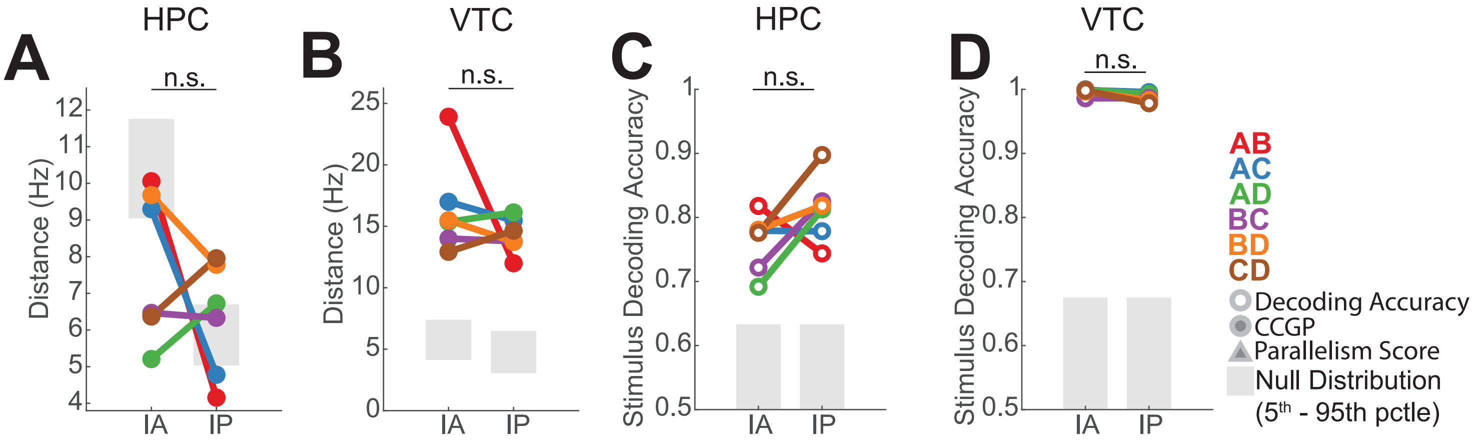
HPC
A
n.s.
Distance (Hz)
IA
IP
VTC
B
n.s.
Distance (Hz)
IA
IP
HPC
C
n.s.
Stimulus Decoding Accuracy
IA
IP
VTC
D
n.s.
Stimulus Decoding Accuracy
IA
IP
AB
AC
AD
BC
BD
CD
Decoding Accuracy
CCGP
Parallelism Score
Null Distribution
(5th - 95th pctle)

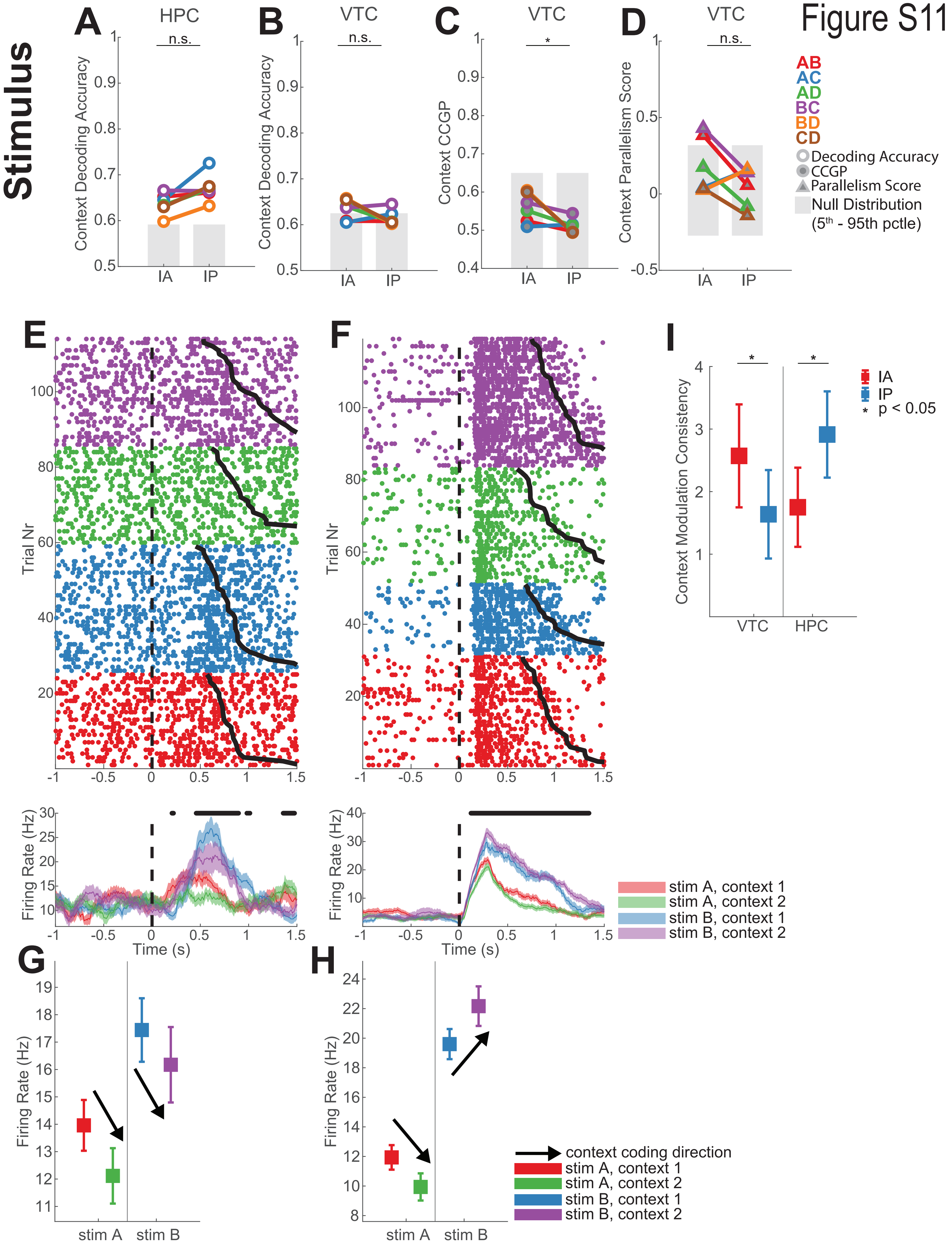
Figure S11
Stimulus
A
HPC
n.s.
Context Decoding Accuracy
IA
IP
B
VTC
n.s.
Context Decoding Accuracy
C
VTC
*
Context CCGP
D
VTC
n.s.
Context Parallelism Score
AB
AC
AD
BC
BD
CD
Decoding Accuracy
CCGP
Parallelism Score
Null Distribution
(5th - 95th pctle)
E
F
Trial Nr
Firing Rate (Hz)
Time (s)
stim A, context 1
stim A, context 2
stim B, context 1
stim B, context 2
I
Context Modulation Consistency
IA
IP
* p < 0.05
VTC
HPC
G
H
Firing Rate (Hz)
stim A
stim B
context coding direction
stim A, context 1
stim A, context 2
stim B, context 1
stim B, context 2

Figure S12

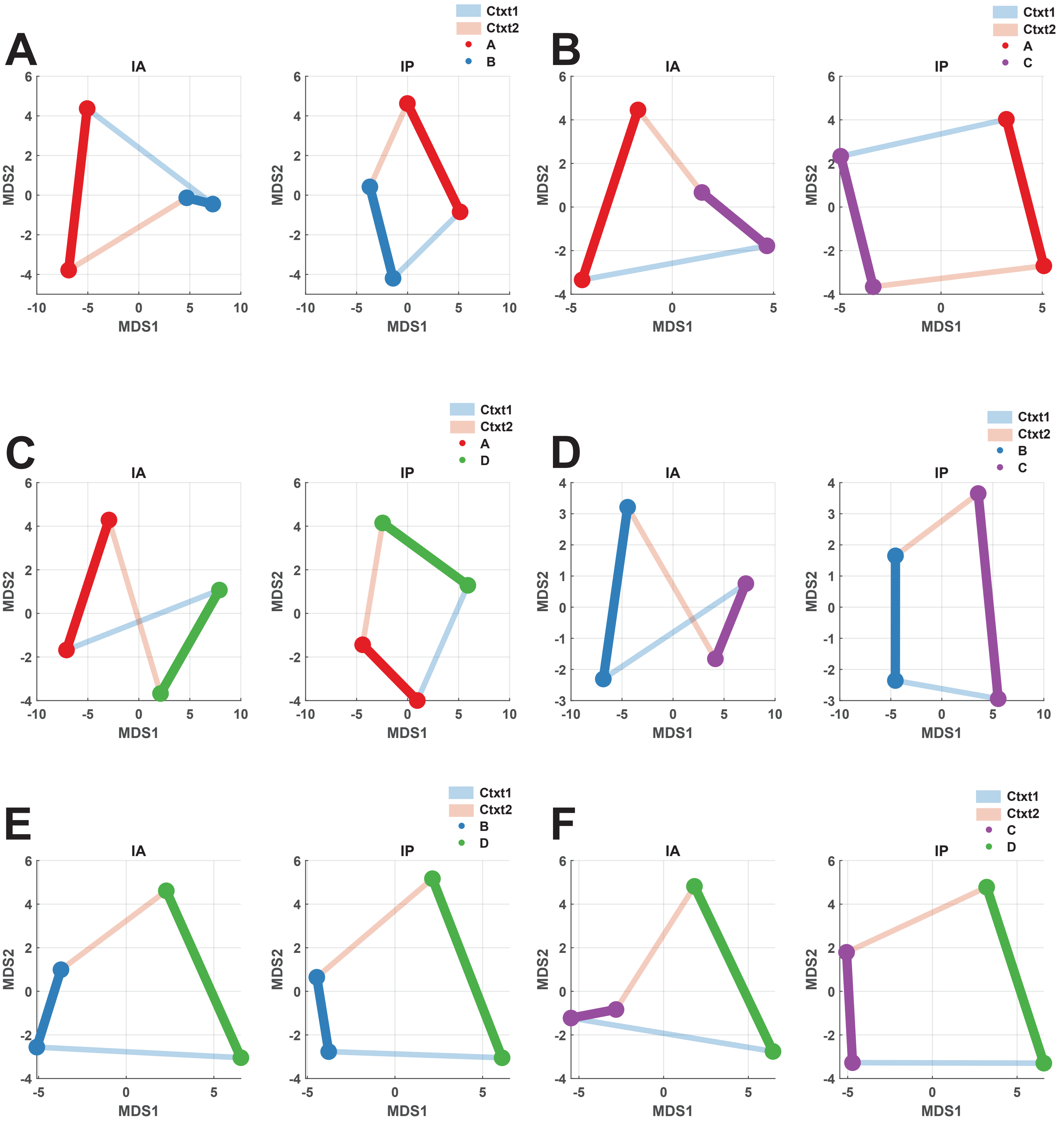

Figure S13

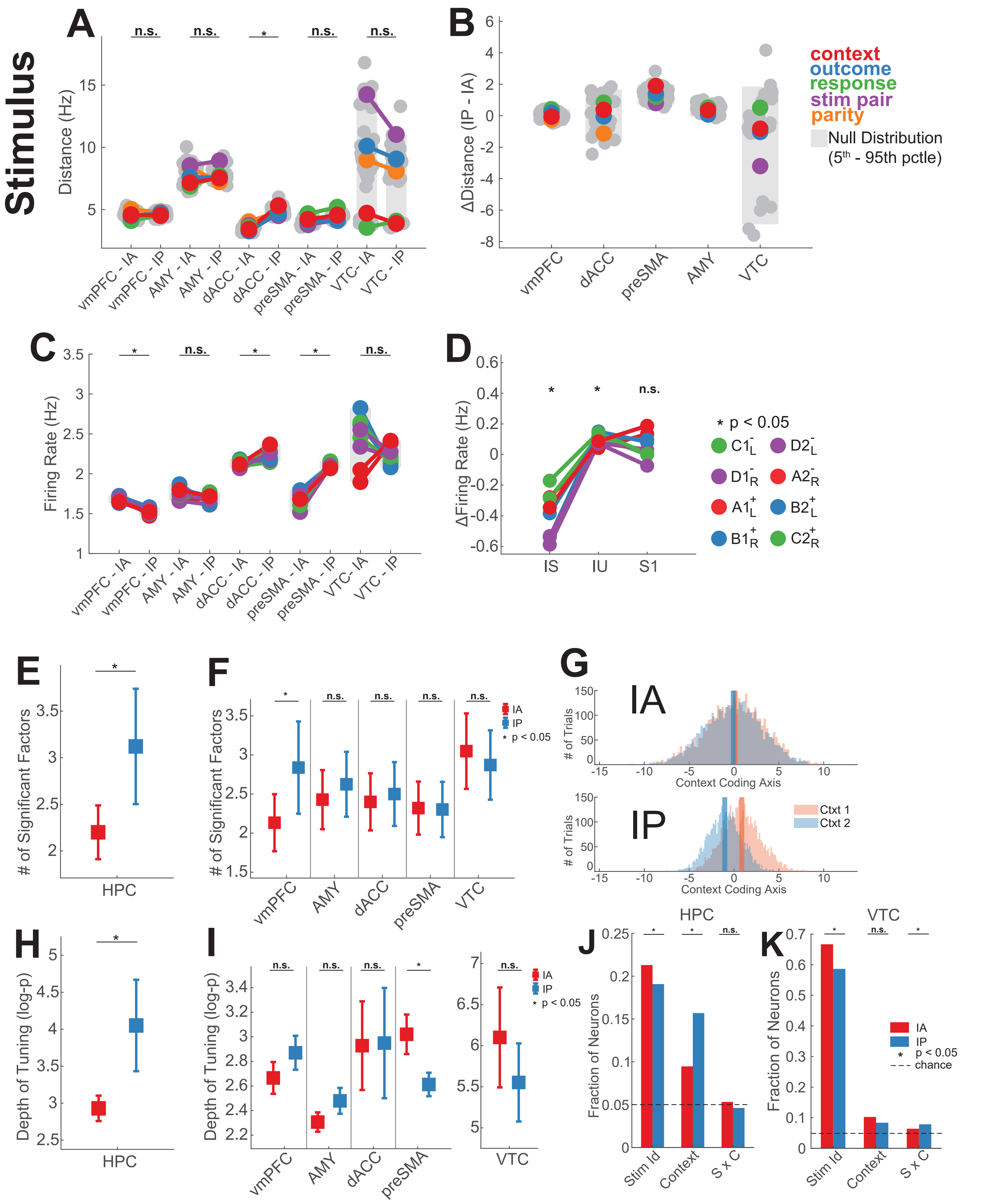
Stimulus
A
n.s.
n.s.
*
n.s.
n.s.
Distance (Hz)
vmPFC - IA
vmPFC - IP
AMY - IA
AMY - IP
dACC - IA
dACC - IP
preSMA - IA
preSMA - IP
VTC- IA
VTC - IP
B
ΔDistance (IP - IA)
vmPFC
dACC
preSMA
AMY
VTC
context
outcome
response
stim pair
parity
Null Distribution
(5th - 95th pctle)
C
Firing Rate (Hz)
D
ΔFiring Rate (Hz)
IS
IU
S1
* p < 0.05
E
### of Significant Factors
HPC
F
IA
IP
* p < 0.05
G
IA
IP
### of Trials
Context Coding Axis
Ctxt 1
Ctxt 2
H
Depth of Tuning (log-p)
I
J
HPC
VTC
K
Fraction of Neurons
Stim Id
Context
S x C
chance

### Baseline

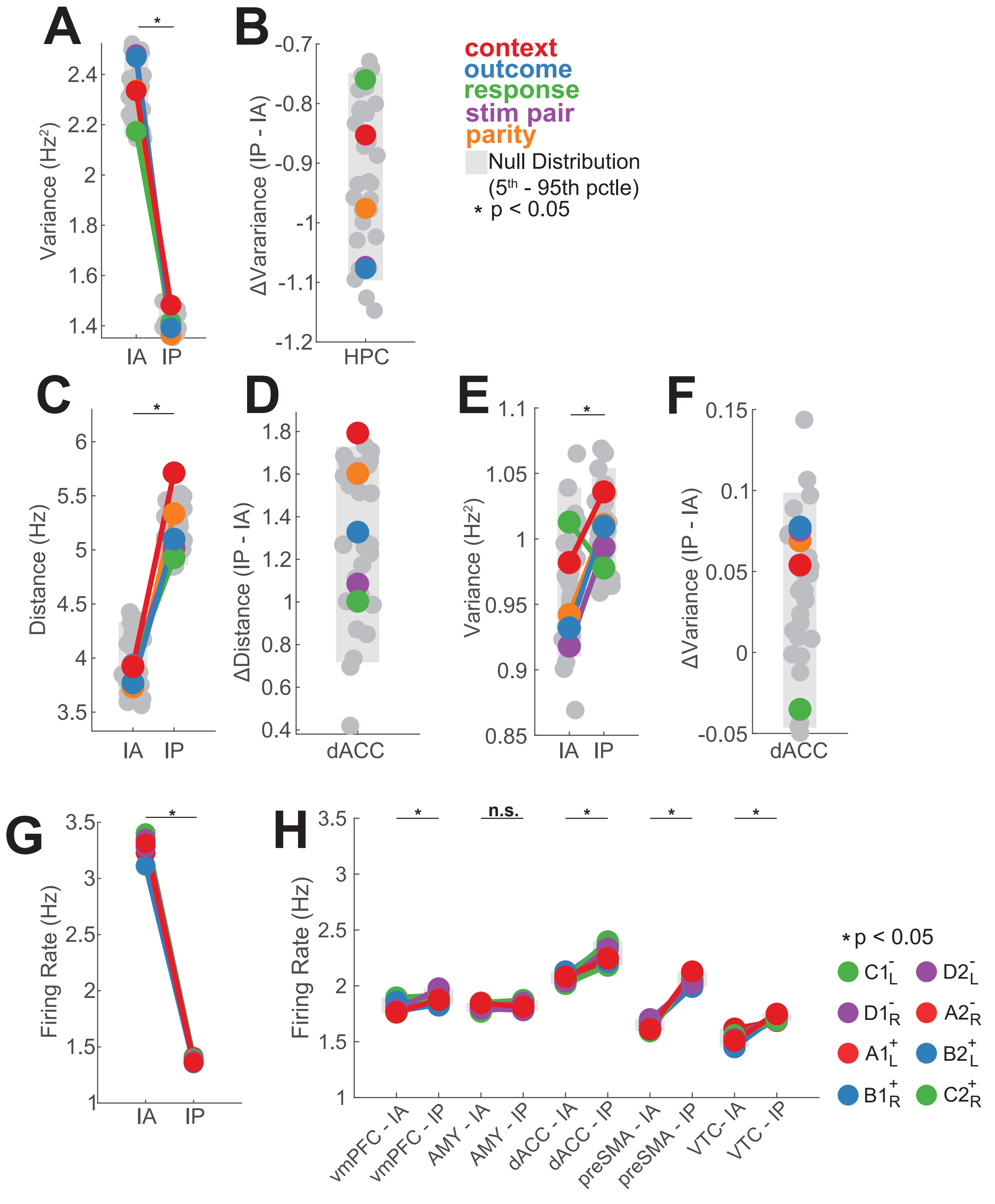
A
B
C
D
E
F
G
H
context
outcome
response
stim pair
parity
Null Distribution
(5th - 95th pctle)
* p < 0.05
Variance (Hz²)
ΔVarariance (IP - IA)
Distance (Hz)
ΔDistance (IP - IA)
Variance (Hz²)
ΔVariance (IP - IA)
Firing Rate (Hz)
Firing Rate (Hz)
IA IP
HPC
dACC
n.s.
vmPFC - IA
vmPFC - IP
AMY - IA
AMY - IP
dACC - IA
dACC - IP
preSMA - IA
preSMA - IP
VTC- IA
VTC - IP
C1
D2
D1
A2
A1
B2
B1
C2

Figure S15

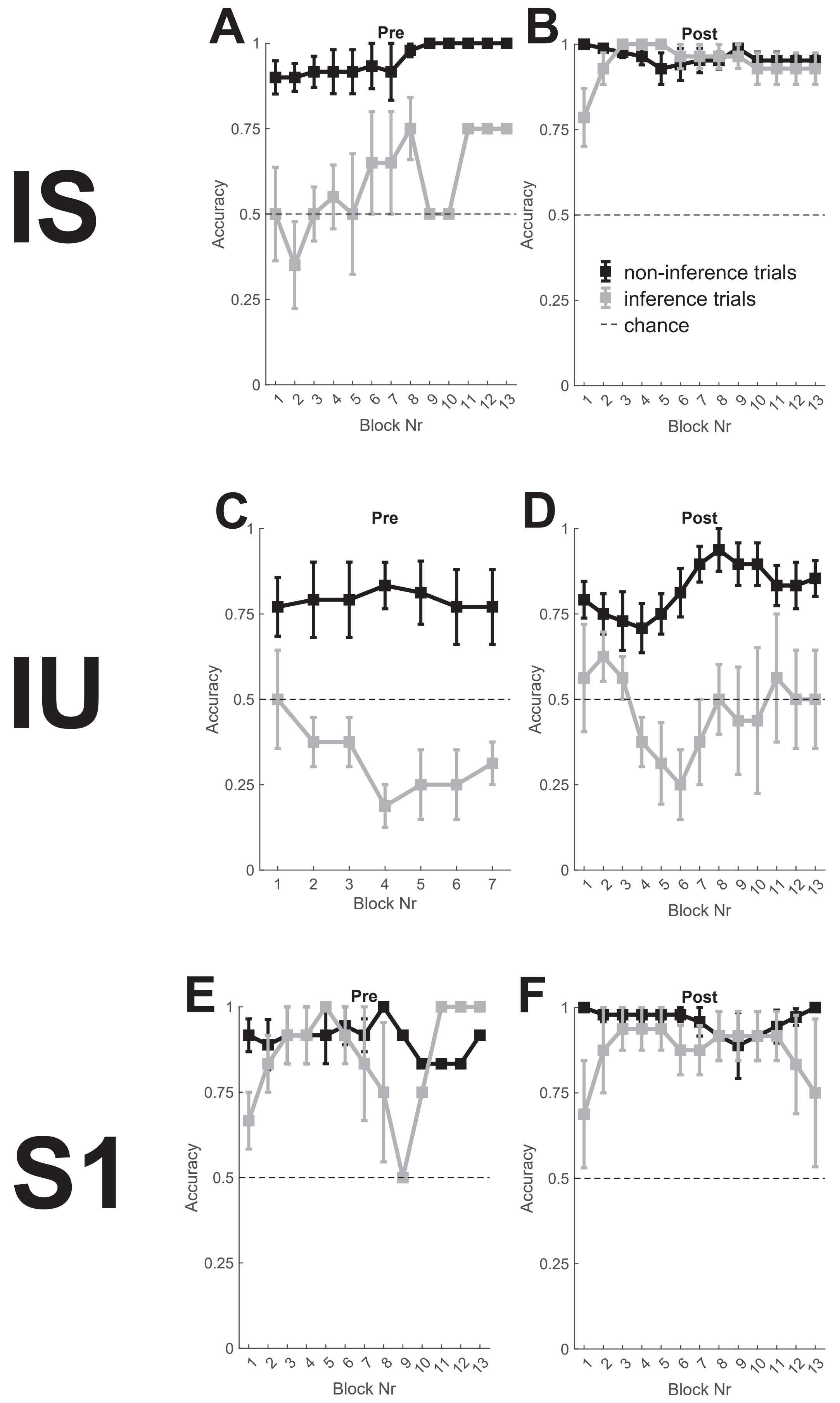

IS
IU
S1
A
B
C
D
E
F
Pre
Post
Accuracy
Block Nr
non-inference trials
inference trials
chance

Figure S16

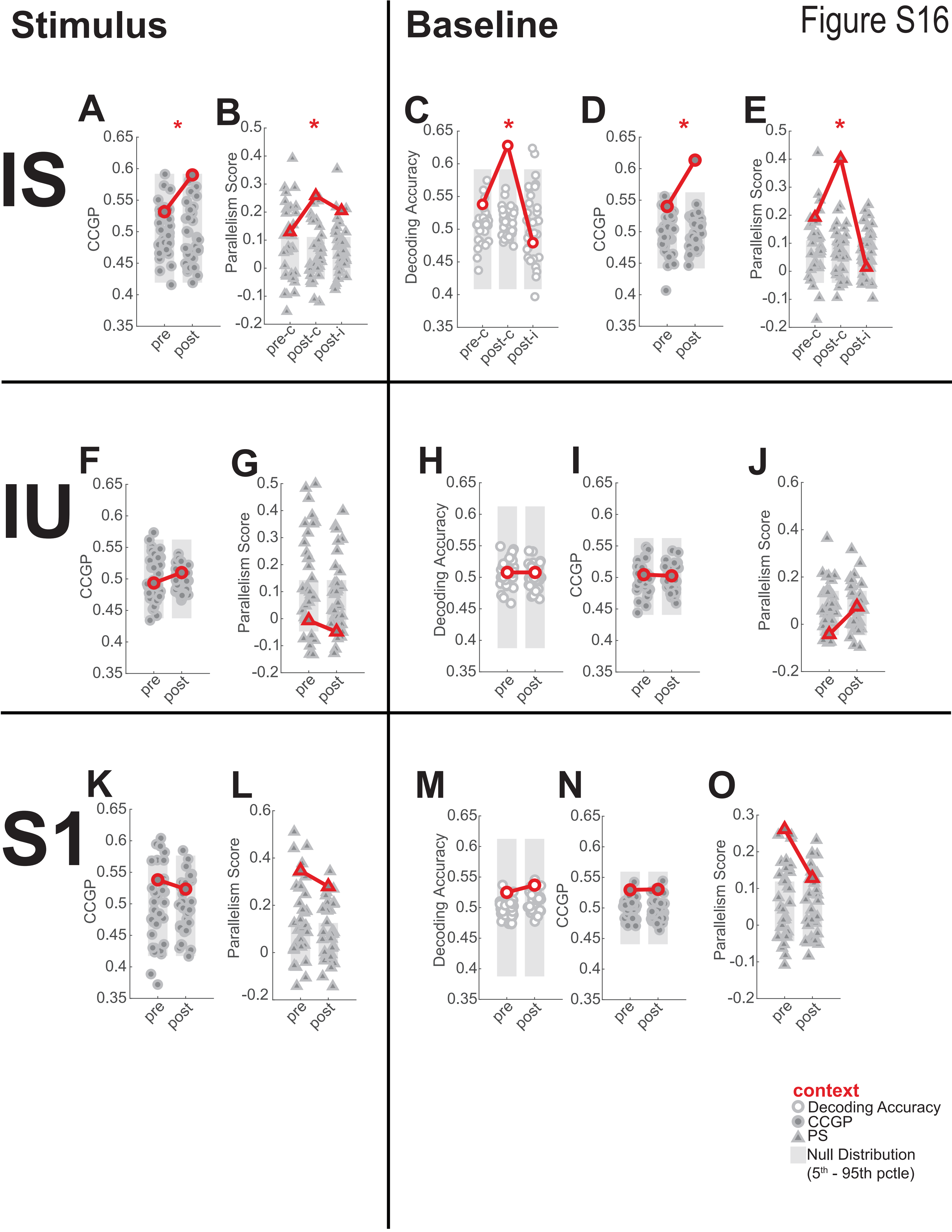

Stimulus
Baseline
IS
IU
S1
A
B
C
D
E
F
G
H
I
J
K
L
M
N
O
CCGP
Parallelism Score
Decoding Accuracy
pre
post
pre-c
post-c
post-i
context
Decoding Accuracy
CCGP
PS
Null Distribution
(5th - 95th pctle)
